# Harnessing full-text publications for deep insights into *C. elegans* and *Drosophila* connectomes

**DOI:** 10.1101/2024.04.13.588993

**Authors:** Karthick Raja Arulprakasam, Janelle Wing Shan Toh, Herman Foo, Mani R Kumar, An-Nikol Kutevska, Emilia Emmanuelle Davey, Marek Mutwil, Guillaume Thibault

**Affiliations:** School of Biological Sciences, Nanyang Technological University, Singapore, 637551; Mechanobiology Institute, National University of Singapore, Singapore 117411

## Abstract

In the rapidly expanding domain of scientific research, tracking and synthesizing information from the rapidly increasing volume of publications pose significant challenges. To address this, we introduce a novel high-throughput pipeline that employs ChatGPT to systematically extract and analyze connectivity information from the full-texts and abstracts of 24,237 and 150,538 research publications concerning *Caenorhabditis elegans* and *Drosophila melanogaster*, respectively. This approach has effectively identified 200,219 and 1,194,587 interactions within the *C. elegans* and *Drosophila* connectomes, respectively. Utilizing Cytoscape Web, we have developed comprehensive, searchable online connectomes that link relevant keywords to their corresponding PubMed IDs, thus providing seamless access to an extensive knowledge network encompassing *C. elegans* and *Drosophila*. Our work highlights the transformative potential of integrating artificial intelligence with bioinformatics to deepen our understanding of complex biological systems. By revealing the intricate web of relationships among key entities in *C. elegans* and *Drosophila*, we offer invaluable insights that promise to propel advancements in genetics, developmental biology, neuroscience, longevity, and beyond. We also provide details and discuss significant nodes within both connectomes, including the insulin/IGF-1 signaling (IIS) and the notch pathways. Our innovative methodology sets a robust foundation for future research aimed at unravelling complex biological networks across diverse organisms.

## INTRODUCTION

The landscape of biological research has experienced a significant transformation over the past two decades, marked by an exponential surge in the volume of scientific publications. This trend is particularly pronounced in the study of model organisms such as *Caenorhabditis elegans* and *Drosophila melanogaster*, which have long served as pivotal systems for understanding fundamental biological processes. A PubMed search reveals the magnitude of this surge: the number of publications related to *C. elegans* has escalated from 6,166 in 2000 to over 40,000 in 2023, while *Drosophila* research has expanded from 40,793 to more than 120,000 publications within the same timeframe. This proliferation of data, while testament to the fields’ dynamism and the research community’s productivity, presents a formidable challenge. Researchers are now faced with the Herculean task of staying abreast of emerging insights and effectively synthesizing vast amounts of information. The critical need for sophisticated tools to navigate, manage, and interpret this growing knowledge base has never been more apparent. Without such innovations, the research community’s capacity to forge novel connections and draw meaningful insights from the wealth of available data may be substantially hindered, potentially slowing the trajectory of scientific progress.

Existing connectome and pathway databases, such as BioGRID, the Gene Ontology (GO), and Reactome, offer valuable insights into the complex networks of gene interactions and biological pathways. BioGRID is an extensive repository for interaction datasets, facilitating the exploration of protein and genetic interactions in a variety of organisms (Stark et al., 2006). However, like many databases, it may not capture real-time research dynamics due to the inevitable delay in data curation (Oughtred et al., 2019). The Gene Ontology (GO) provides a comprehensive framework for the representation of gene function across species, yet it can be constrained by its static consensus terminology and may not capture the full spectrum of gene functionality or recent discoveries (Gaudet & Dessimoz, 2017). Reactome, while detailing pathways of numerous biological processes, could also potentially miss species-specific functions and unique cellular conditions (Milacic et al., 2024). Despite the undeniable utility of these tools, they are not without limitations. Their reliance on curated data ensures accuracy but can result in updates lagging behind the latest literature due to the labor-intensive nature of manual curation. Moreover, these databases might not fully encapsulate the multifaceted relationships between genes, such as epistatic interactions, genetic modifiers, and context-dependent effects. Such relationships are essential for a comprehensive understanding of complex phenotypes and diseases. This underscores a crucial gap: the need for an advanced tool capable of dynamically incorporating the latest findings and representing the intricate web of genetic interactions with both depth and breadth.

In the fields of *C. elegans* and *Drosophila* research, gold-standard organism-specific databases such as WormBase and FlyBase have been instrumental in collating vast amounts of genetic information (Gramates et al., 2022; Harris et al., 2010). These platforms offer insights into gene function, protein-protein interactions, and phenotypic data crucial for understanding development, aging, and disease in these model organisms. WormBase, for instance, has been a pivotal tool for *C. elegans* researchers, providing curated genetic, genomic, and biological information. Similarly, FlyBase serves the *Drosophila* community by compiling data on genetic and molecular attributes of *Drosophila* genes and genomes. However, even these comprehensive repositories may not fully capture the dynamic and rapidly evolving insights emerging from current literature. Key information on context-specific gene interactions, the influence of environmental factors on genetic pathways, and the subtleties of temporal and spatial gene expression patterns are often more thoroughly detailed in individual studies. As such, there exists a gap between curated databases and the nuanced, high-resolution data that can be mined from full-text publications, which often contain rich, yet uncurated, insights into gene function and regulation.

In this work, we unveil a high-throughput text-mining pipeline designed to systematically extract and analyze gene-related information from a vast array of research publications concerning *C. elegans* and *Drosophila*. Utilizing the capabilities of natural language processing technologies, this pipeline transcends the confines of traditional databases to offer a dynamic, enriched view of the genetic interaction landscapes of these model organisms. We detail the development and deployment of our pipeline, showcasing its unparalleled ability to uncover and visualize intricate networks of gene interactions and biological pathways. Through this effort, we aim to equip researchers with a robust tool to navigate the growing wealth of genetic and biological information in *C. elegans* and *Drosophila*, thereby catalyzing significant advances in our systemic understanding of biology. Our study underscores the feasibility and the transformative impact of integrating advanced computational methods with bioinformatics to enhance our grasp of complex biological systems. The *C. elegans* and *Drosophila* Connectomes, developed as a result of this endeavor, are accessible at http://worm.connectome.tools and http://drosophila.connectome.tools, respectively, serving as comprehensive portals to an array of interactions and pathways.

## RESULTS

### Semantic Analysis of Thousands of Abstracts and Full-Text Papers

In our comprehensive analysis, we processed a substantial amount of literature pertaining to gene function within *C. elegans* and *Drosophila*. To construct the *C. elegans* and *Drosophila* Connectomes, we systematically searched scientific research papers for occurrences of *C. elegans* or *Drosophila* genes in their titles and abstracts. We prioritized full-text papers from open-access publications, those freely available in PubMed Central (PMC), and accessible through Elsevier via the NTU Library. Alternatively, it went back to fetch only titles and abstracts if full-text unavailable. This approach ensured that the selected papers were directly relevant to the genetic makeup and biological processes of these model organisms. The result was a curated selection of articles that form the backbone of our *C. elegans* or *Drosophila* Connectomes databases. Initially, we extracted 24,237 *C. elegans*-related articles, comprising 9,904 full-text articles and 14,332 abstracts. For *Drosophila*, the amount included an even larger set of 150,538 articles, with 71,226 full-text articles and 79,311 abstracts. Articles were initially categorized based on gene nomenclature, which necessitated a subsequent deduplication step due to multiple occurrences of the same articles across different categories. Following this curation process, we filtered the dataset to a final tally of 12,062 articles for the *C. elegans* connectome and 36,372 articles for the *Drosophila* connectome. These articles span a wide distribution across 925 journals for *C. elegans* and 1,815 journals for *Drosophila* (Table S1). For a more detailed visualization, we have compiled the top 40 most frequently cited journals in both fields, demonstrating the quantitative distribution of the articles (Figures 1A and 1B). These articles form the foundation of our subsequent analysis using GPT, enabling a refined exploration of the genetic and molecular interplays that define the complex connectomes of these model organisms.

**Figure 1.**
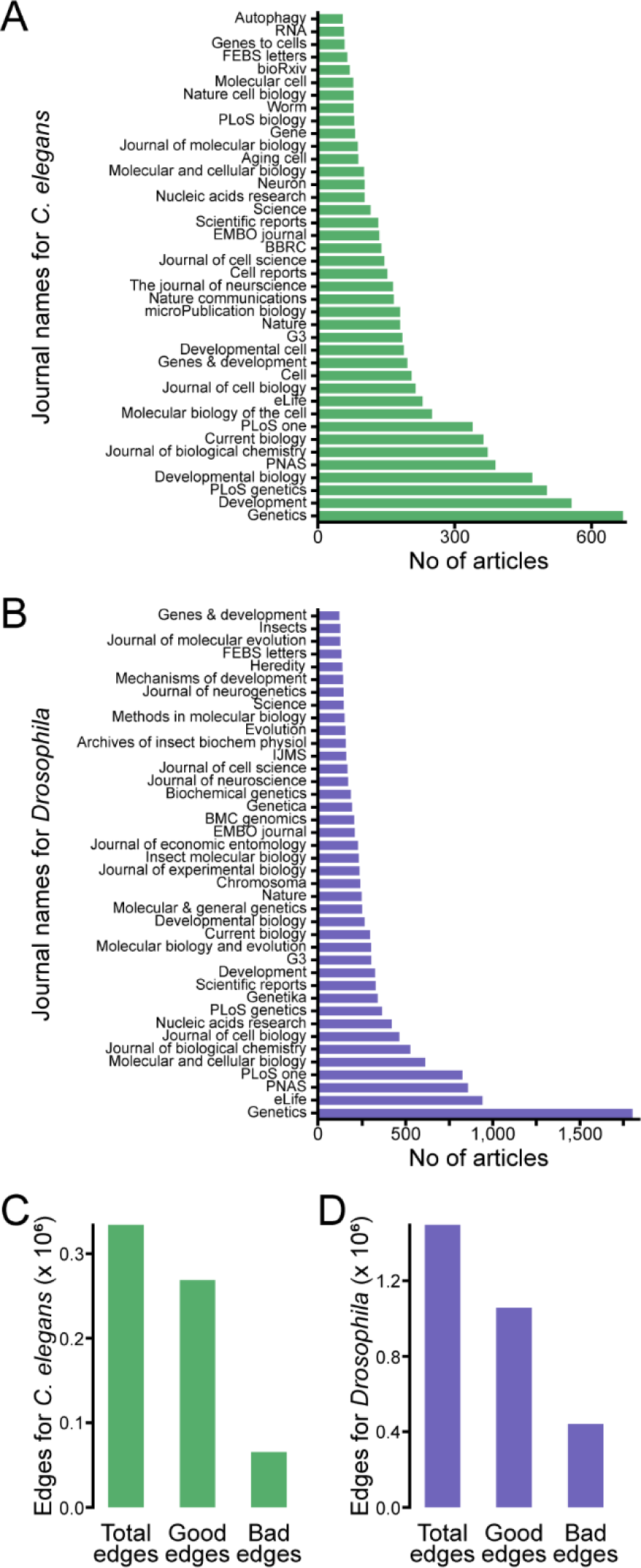
Comprehensive meta-analysis of full-text articles in *C. elegans* and *Drosophila*. (**A-B**) Quantitative distribution of articles from the top 40 journals featuring *C. elegans*-(**A**) and *Drosophila*-related (**B**) research. (**C-D**) Profile of *C. elegans* (**C**) and *Drosophila* (**D**) interaction data showing total, validated (good), and erroneous (bad) edges (in millions).

We exploited the capabilities of OpenAI’s GPT API to extract functional relationships between entity pairs (e.g., gene A interacts with gene B), infer functional annotations of genes (e.g., gene A is implicated in dauer formation), and delineate abbreviations (e.g., FOXO for FORKHEAD BOX O). To optimize our query efficiency, we iteratively tested different prompts with ChatGPT until we refined a distinct prompt (Table S2). The prompt was systematically applied across all abstracts and full-text articles, revealing 200,219 functional relationships for the *C. elegans* connectome and 1,194,587 for the *Drosophila* connectome, along with 112,128 gene function annotations and 73,591 abbreviation identifications. To evaluate the quality of GPT’s output, we examined the relationships or ‘edges’ that it identified between two genes and/or entities within the literature network. The GPT analysis generated statements in the format of “Entity1! Relationship! Entity2”. We classified edges as “good” when at least one of the entities appeared in the GPT analysis, and as “bad” when neither entity was found. For *C. elegans*, out of 334,104 total edges, 268,632 were designated as “good”, while 65,472 were “bad” (Figure 1C). In parallel, the *Drosophila* connectome revealed 1,498,156 edges, with 1,056,937 categorized as ‘good’ and 441,319 as “bad” (Figure 1D). Such results lend confidence to the integrity of GPT’s output, affirming that most relationships identified align with the expected analytical format. Next to evaluate the precision of our text-mining process, we conducted a manual accuracy assessment on a random sample of 50 abstracts. The outcomes indicated that GPT predominantly identified relationships correctly (Figure 2, Documents S2 and S3). However, we did observe instances where relationships were either missed or inaccurately characterized. Such inaccuracies were notably prevalent in abstracts and full-text articles that did not mention specific gene names, leading to instances of GPT “hallucinating” entities, incorrectly designating them as “gene”.

**Figure 2.**
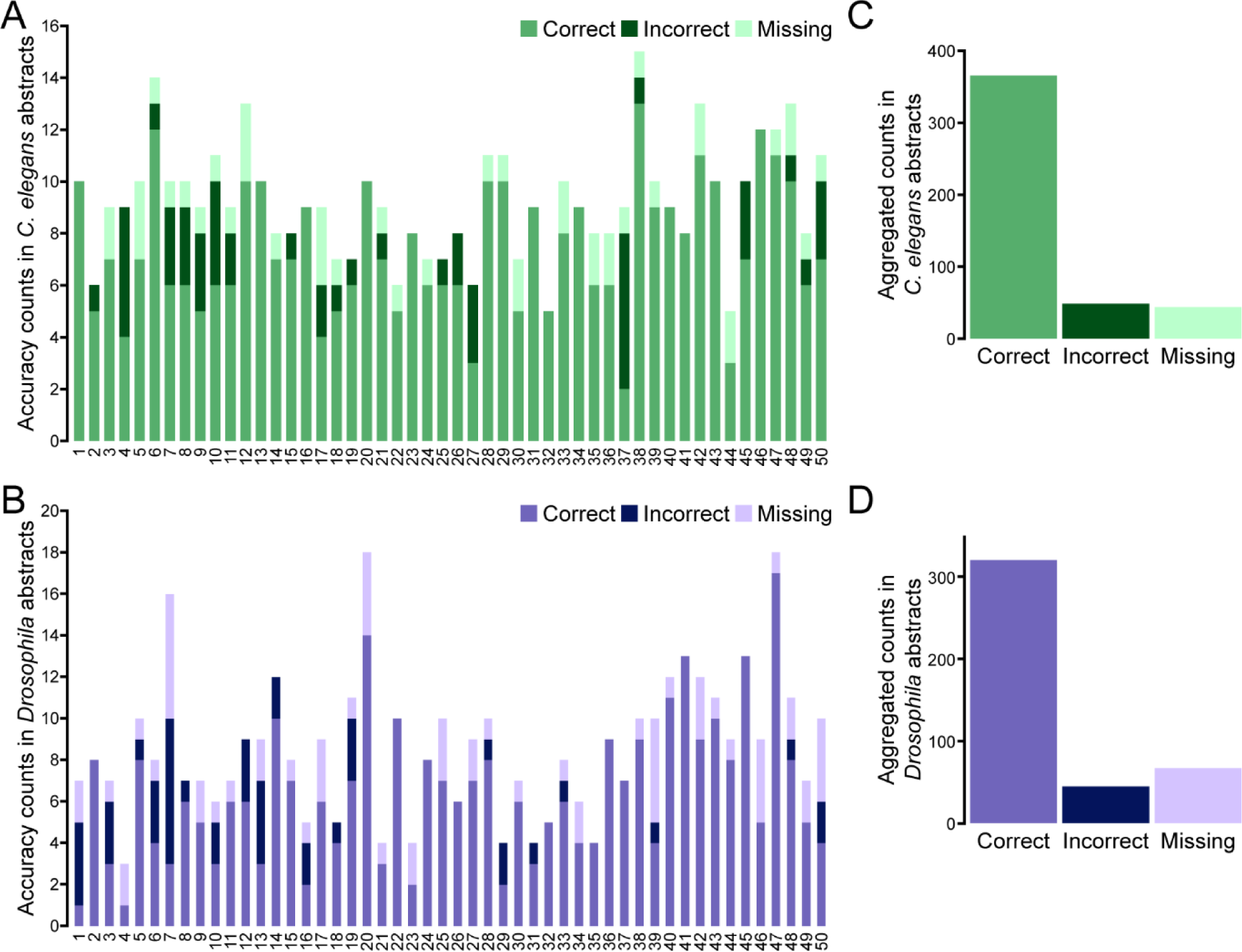
Accuracy assessment of GPT-processed abstracts for model organisms. (**A-B**) Analysis accuracy for 50 *C. elegans* (**A**) and 50 *Drosophila* (**B**) abstracts, categorized by the incidence of correct, incorrect, and missing information. (**C-D**) Aggregated counts of accurate (correct), erroneous (incorrect), and overlooked (missing) statements from *C. elegans* (**C**) and *Drosophila* (**D**) abstracts shown in (A) and (B), respectively.

### Expanding the *C. elegans* Interaction Map for a Connectome with Comprehensive Coverage

Leveraging the outputs from GPT analysis, which consist of pairwise relationships between entities, we crafted a network that encapsulates the full spectrum of functional relationships within the *C. elegans* biological system. Despite GPT’s instruction to prioritize genes, (Table S2) the analysis yielded interactions that included not only gene-gene interactions but also connections to biological functions, pathways, and phenotypes (Figure 3). During the curation process, we encountered numerous instances where gene functions were discussed generically, using placeholders such as “gene”, or the organism names “*Caenorhabditis elegans*” and “worm” were utilized as proxy for specific genetic entities. These non-specific terms were subsequently removed to sharpen the focus on the top 20 most meaningful entities within the *C. elegans* pool. The analysis spotlighted “*daf-16*” and “LIFESPAN” as the most prevalent topics in *C. elegans* literature, underscoring their significance in the field (Figure 3B). Moreover, examining the types of relationships revealed that “regulated”, “requires”, and “interacts with” emerged as the most common edges, delineating the primary modes of genetic and molecular interactions in *C. elegans* (Figures 3A and 3D). To understand the pattern of mostly investigated genes we pulled out the information of top 5,000 entities, edges between entities, genes as entities, and edges between genes that are appeared in both *C. elegans* and *Drosophila* connectome analysis (Table S3-S6). This analysis showed that “*lin-12*”, “*let-23*“, and “*par-3*” are mostly investigated genes in *C. elegans* (Figure 3C). The comparative frequency of these edges demonstrates the thematic congruencies within the biological research of these model organisms, reflecting shared foundational processes that are central to understanding their complex biology. The connectome thus constructed offers a comprehensive view of the intricate web of interactions that define *C. elegans* biology, serving as a valuable resource for researchers navigating this model organism’s extensive genetic landscape.

**Figure 3.**
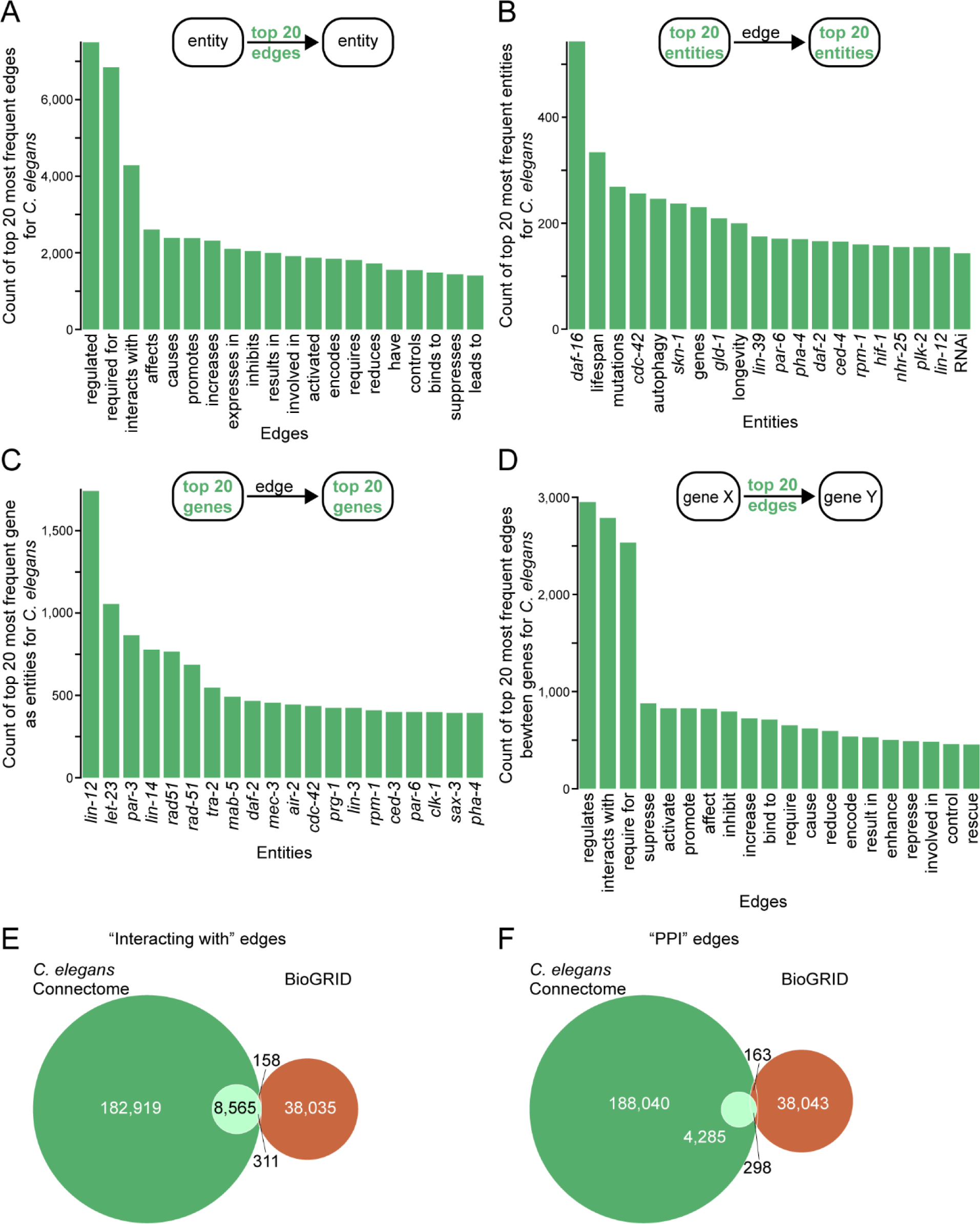
Network analysis highlights in *C. elegans* Connectome. (**A**) Frequency distribution of the top 20 interaction edges in the *C. elegans* connectome. (**B**) Frequency distribution of the top 20 interaction entities in the *C. elegans* connectome. (**C**) Frequency distribution of the top 20 interaction entities as genes in the *C. elegans* connectome. (**D**) Frequency distribution of the top 20 interaction edges between genes in the *C. elegans* connectome. (**E-F**) Venn diagram illustrating the commonalities “interacting with” (**E**) and protein-protein interaction (PPI) (**F**) edges in the *C. elegans* Connectome and the interacting proteins listed in BioGRID.

Further analysis was conducted to assess the comprehensiveness of the *C. elegans* Connectome in relation to the established BioGRID database, focusing on interacting edges which includes “interact”, “bind”, and “phosphorylate” relationships from the whole connectome database and the protein-protein interaction (PPI network) only from connectome database (Figures 3E and 3F). We identified 8,565 interacting edges in the connectome. Notably, the Connectome and BioGRID share 311 identical interacting partners. Additionally, there are 158 interactions catalogued in BioGRID that overlap with the Connectome; these, however, are not specifically defined as interacting. In terms of the PPI network, the *C. elegans* Connectome demonstrated an overlap of 298 PPI interacting edges with BioGRID, yet it also identified an additional 4,285 interacting edges not catalogued by BioGRID (Figure 3F). These findings highlight the Connectome’s strength in detecting not only the commonly recognized interactions but also a broader spectrum of biological relationships. Furthermore, the Connectome’s capability to capture more nuanced interaction types beyond interacting— such as “colocalize with” and “inhibits”—affirms its utility in providing a more detailed and extensive mapping of molecular interactions (Figure 3).

In our comprehensive mapping of the *Drosophila* connectome, we delineated a network equally rich and intricate as its *C. elegans* counterpart. The extent of interaction types identified among *Drosophila* genes, with “regulates” and “interacts with” emerging as particularly common, indicating frequent protein-protein and gene regulatory interactions (Figure 4A). Broader biological entities and processes that are recurrently discussed, such as “apoptosis” and “lifespan”, highlight their fundamental importance in *Drosophila* studies (Figure 4B). The genes that dominate the *Drosophila* Connectome, such as “*VIA*”, “*cycle*” and “*actin*”, underscoring their prominence and frequent investigation within the species’ genetic research (Figure 4C). It is important to note that terms like “VIA” and “cycle” cover both specific gene names and instances where these words do not refer to genes in the prompts/connectomes. Due to this, such terms cannot be distinctly identified as gene-related by ChatGPT without additional contextual analysis. Complementary , the most common gene-to-gene edges include “has”, “regulates” and other informative terms on the type of interactions such as “is required for”, illustrating the extensive interconnectivity within the *Drosophila* genome (Figure 4D). Our Connectome contained 44,382 edges categorized as “interacting” (Figure 4E). Of these, 289 were found to directly correspond with interactions listed in BioGRID. Additionally, the Connectome shared another 132 interactions with BioGRID, though these were not explicitly categorized under the “interacting” descriptor. Regarding the protein-protein interaction (PPI) network, a comparison revealed that 19 PPI edges were common between the *Drosophila* Connectome and BioGRID (Figure 4F). Nevertheless, our Connectome unveiled 9,101 additional PPI edges, absent in BioGRID’s catalog. Together, the analysis of both the *C. elegans* and *Drosophila* connectomes illuminates their potential to not only complement but substantially augment existing genetic databases. By unveiling a multitude of previously unavailable interactions, these connectomes serve as unconventional resources that encapsulate the evolving complexity of biological research. They provide researchers with dynamic and current tools essential for exploring into the genetic and molecular fabric of these model organisms.

**Figure 4.**
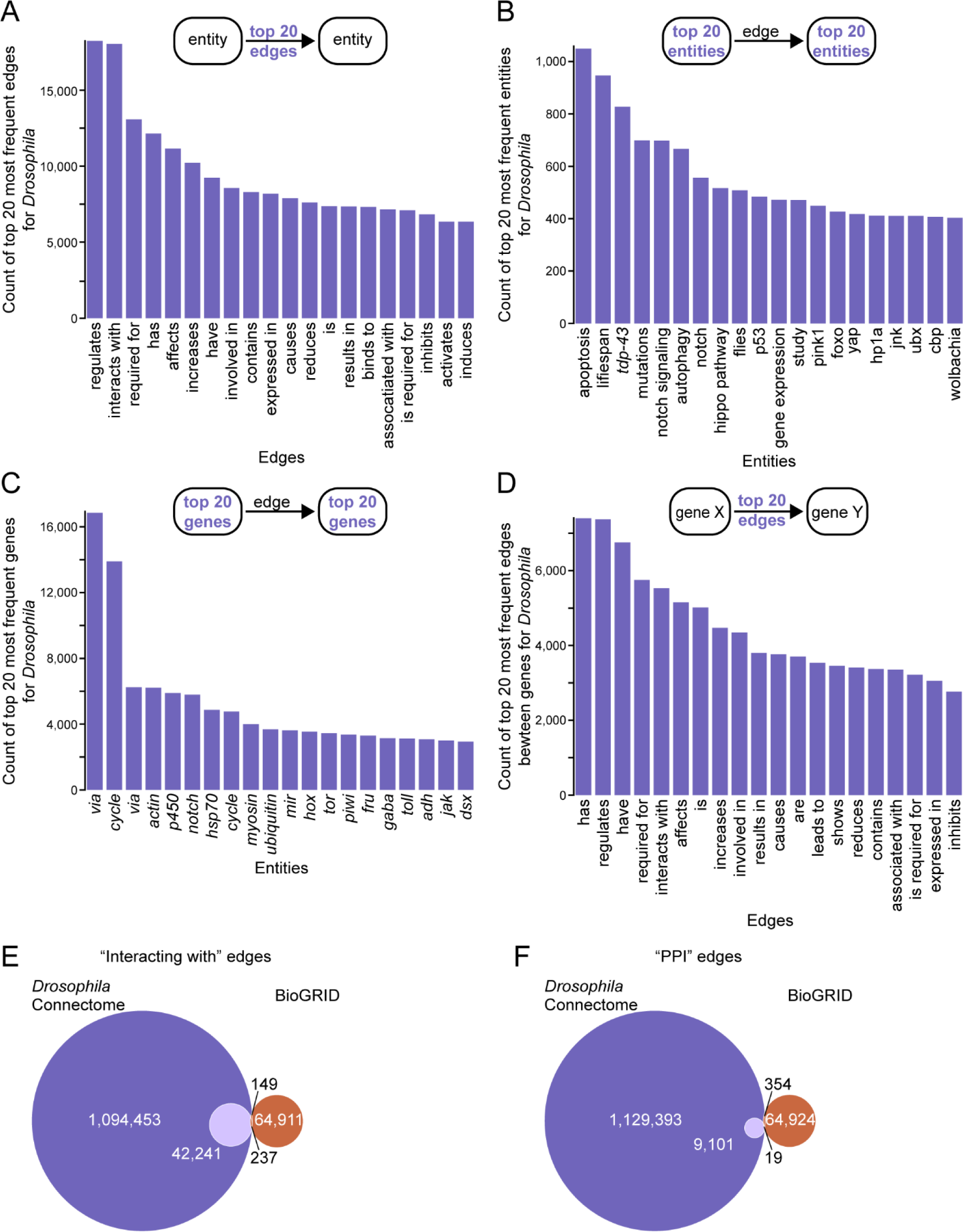
Network analysis highlights in *Drosophila* connectome. (**A**) Frequency distribution of the top 20 interaction edges in the *Drosophila* connectome. (**B**) Frequency distribution of the top 20 interaction entities in the *Drosophila* connectome. (**C**) Frequency distribution of the top 20 interaction entities as genes in the *Drosophila* connectome. (**D**) Frequency distribution of the top 20 interaction edges between genes in the *Drosophila* connectome. (**E-F**) Venn diagram illustrating the commonalities “interacting with” (**E**) and protein-protein interaction (PPI) (**F**) edges in the *Drosophila* Connectome and the interacting proteins listed in BioGRID.

### Interactive Connectome Platforms for *C. elegans* and *Drosophila*

The *C. elegans* and *Drosophila* Connectomes offer an interactive gateway to a vast array of genetic interactions, parallel to a digital atlas for biological functions within these model organisms. Through an intuitive interface, users can query genes, proteins, and other entities, obtaining detailed information pages that include GPT-generated abbreviations, functional annotations, and direct links to scientific articles. The platforms feature KnowledgeNetworks, visual representations of the connectome that allow users to customize the view, isolate nodes, and even download data for advanced analysis. A comprehensive summary accompanies each network, providing immediate insights into the most connected nodes and their respective literature sources. For those requiring programmatic access, an API delivers the connectome’s array of data in a structured format. Both Connectomes have been constructed with the same dedication to accessibility and depth of information as the PlantConnectome that we recently reported (Fo et al., 2023).

In the space of model organisms, few genes have garnered as much attention as *daf-16* in *C. elegans*, a gene whose conservation extends to its *Drosophila* counterpart, the fokhead box protein O (*foxo*). These genes pivotal connections within the complex networks of *C. elegans* and *Drosophila* biology, influencing essential processes like development, aging, metabolism, and stress response. DAF-16 and FOXO, transcription factors that interact with genes containing daf-16/FOXO binding elements (DBE), play crucial roles in the Insulin/IGF-1-like signaling (IIS) pathway. A query for “*daf-16*” and “*foxo*” within the *C. elegans* and *Drosophila* Connectomes maps a network sourced from 514 and 213 papers, respectively. Refinement of the search to “regulates” within the “Layout Options” reveals a more focused network from 61 and 71 papers for *C. elegans* and *Drosophila* (Figures 5A and 5B), respectively, featuring *daf-16* and *foxo*’s roles in regulating lifespan, feeding behaviors, dauer development, and stress resistance among other processes.

**Figure 5.**
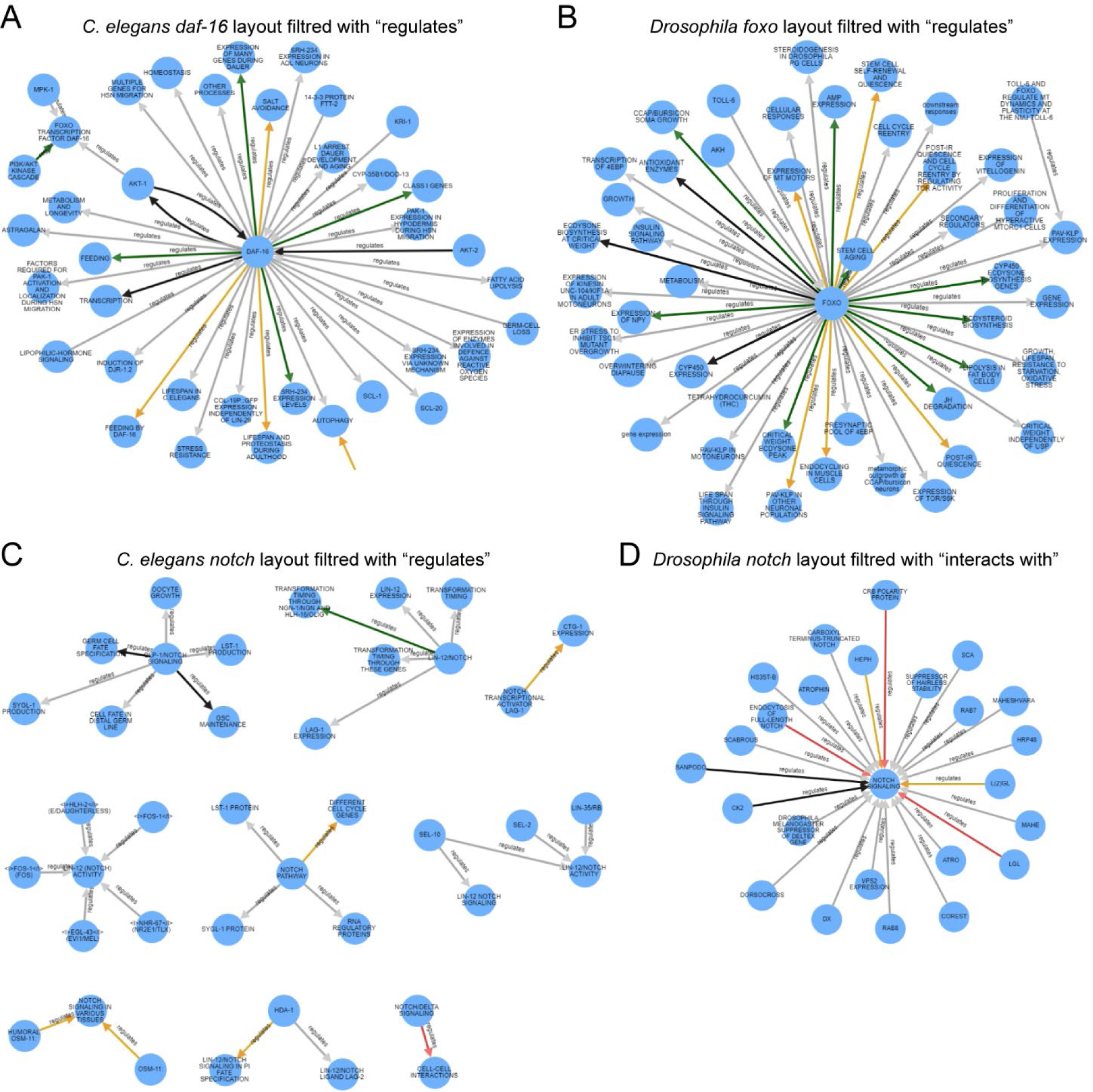
Connectome visualization for key genes in *C. elegans* and *Drosophila*. (**A**) Knowledge network of *C. elegans* gene *daf-16* with Layout Options “regulates”. (**B**) knowledge network of *Drosophila* gene *foxo* Layout Options “regulates”. (**C**) Knowledge network of *C. elegans* “notch” genes with Layout Options “regulates”. (**D**) Knowledge network of *Drosophila* gene *notch* with Layout Options “interacts with”.

The *C. elegans* Connectome elucidates *daf-16’*s regulation of genes such as “CYP-35B1/DOD-13” (Iser et al., 2011), “SCL-1” (Ookuma, Fukuda & Nishida, 2003), “PAK-1” (Kennedy, Pham & Grishok, 2013), “COL-19P::GFP” (Wirick et al., 2021), and “SRH-234” (Gruner et al., 2014). Similarly, it highlights how *daf-16* and *foxo* regulate specific phenotypes in *C. elegans* and *Drosophila*, such as “FATTY ACID LIPOLYSIS” (Antebi, 2013) or “LIPOLYSIS IN FAT BODY CELLS” (Roy & Palli, 2018) and “L1 ARREST, DAUER DEVELOPMENT, AND AGING” (Kaplan & Baugh, 2016) or “STEM CELL AGING” (Artoni et al., 2017). These connections and their respective publications are directly accessible through one-click links, demonstrating the tool’s efficacy in providing a rapid, comprehensive overview of protein interactions.

The highly conserved Notch signaling pathway, pivotal in cell fate determinations, was initially discovered in *Drosophila*, marked by its role in wing morphology. Subsequently, the *C. elegans* counterparts, *lin-12* and *glp-1*, were identified, spotlighting their essential functions in developmental processes (Priess, Schnabel & Schnabel, 1987; Greenwald, Sternberg & Horvitz, 1983; Austin & Kimble, 1987). This pathway plays a crucial role in a multitude of biological processes (Kopan & Ilagan, 2009; Suarez Rodriguez, Sanlidag & Sahlgren, 2023), operating through cell-cell interactions initiated by transmembrane ligands that activate Notch receptors on adjacent cells. The activation leads to the cleavage of the Notch receptor’s cytosolic domain, which then moves to the nucleus to regulate gene expression.

Leveraging the “notch” query within both the *C. elegans* and *Drosophila* Connectomes yielded networks sourced from 225 and 684 papers, respectively. Further refinement using “regulates” or “interacts with” in the “Layout Options” revealed more focused networks, sourced from 30 and 116 papers for *C. elegans* and *Drosophila*, respectively (Figures 5C and 5D). This analysis underscored the Notch pathway’s regulation of “LIN-11 EXPRESSION (Marri & Gupta, 2009), “GERM CELL FATE SPECIFICATION” (Lee et al., 2016), and “C.ELEGANS BEHAVIOR” (Chao et al., 2005) in *C. elegans*. In contrast, the *Drosophila* Connectome illustrated a richer interaction network for the Notch pathway, indicating a more extensive exploration of its interacting partners in *Drosophila* or its suitability as a model organism for studying these interactions. Among the highlighted interactions were “NOTCH” interacts with “MKK4” (Zhou et al., 2021), “DMYC EXPRESSION” (Sun et al., 2008), “MORE THAN 300 GENES” (Ho, Pallavi & Artavanis-Tsakonas, 2015), and “AKAP200” (Bala Tannan et al., 2018), showcasing the pathway’s broad influence across *Drosophila*’s genetic landscape.

## DISCUSSION

In this study, we unveiled the comprehensive Connectomes for *C. elegans* and *Drosophila*, charting a vast landscape of genetic interactions and biological functions pivotal to these model organisms. Our analysis, underpinned by the innovative application of GPT technology, has facilitated the identification and cataloging of hundreds of thousands of genetic relationships, encompassing both well-documented and previously unexplored interactions. Notably, genes such as *daf-16*/*foxo* in *C. elegans* and its functional counterpart in *Drosophila*, alongside the Notch signaling pathway, emerged as significant nodes within these networks. These hubs not only underscore the genetic complexity inherent in biological processes like development, aging, and stress response but also highlight the Connectomes’ capacity to unearth interactions that span across a broad spectrum of biological research. The creation of these Connectomes marks a significant stride in our ability to navigate the genetic intricacies of *C. elegans* and *Drosophila*, offering an enriched resource that advances our comprehension of their genetic frameworks and sets the stage for future discoveries in genetic regulation and function.

The introduction of our *C. elegans* and Drosophila Connectomes represents a significant augmentation to databases like BioGRID, which catalogs nearly 1.6 million interactions across various species through detailed literature annotations (Stark et al., 2006; Oughtred et al., 2019). Our connectomes, by mining full-text publications via computational techniques, offer a complementary approach. This methodology not only enriches the database with the latest research findings but also unravels biological contexts and intricate details of interactions, facilitating a more nuanced understanding of genetic networks. While BioGRID’s manual curation process ensures the accuracy and reliability of its data (Oughtred et al., 2019), it may encounter challenges in rapidly integrating new discoveries. Our connectomes aim to mitigate this gap, leveraging natural language processing technologies to capture and incorporate emerging insights directly from the expansive volume of research articles. However, it’s essential to recognize the foundational role of databases in the bioinformatics field. Their rigorously vetted information provides a valuable cornerstone that our computational approach seeks to extend, not supplant. The inclusion of CRISPR screen datasets into BioGRID signifies a notable expansion in the types of data curated, reflecting an evolution that our connectomes parallel through the adoption of advanced data mining techniques (Salwinski et al., 2009; Murugesan, Abdulkadhar & Natarajan, 2017). By integrating the strengths of manual curation with the scalability of automated text-mining, we aspire to create a synergistic resource. This combined approach aims to offer researchers a rounded view of the biological landscape, enabling a deeper understanding and facilitating discoveries in the genetics of *C. elegans* and *Drosophila*.

In constructing our *C. elegans* and *Drosophila* Connectomes, we aimed to address some of the challenges inherent in manual curation processes. Our strategy prioritized articles accessible at no cost, including open-access publications and those available through PubMed Central (PMC) and Elsevier via the NTU Library. While this approach has allowed us to compile a vast and comprehensive database, it inherently limits our ability to immediately incorporate the latest research findings beyond titles and abstracts. This delay in integrating new studies poses a challenge in maintaining the most current view of complex biological networks. Recognizing this limitation, we are exploring innovative strategies to further enhance the timeliness and comprehensiveness of our database. Future directions could include forming collaborations with publishers to secure earlier access to research findings and developing automated text-mining tools that can more rapidly identify and incorporate relevant studies. By augmenting our current resources with these advanced methodologies, we will aim to minimize delays and ensure our databases remain at the forefront of biological research. These steps should not only improve the immediacy of data curation but also reinforce our commitment to providing a dynamic and cutting-edge resource for the scientific community.

In conclusion, the advent of the *C. elegans* and *Drosophila* Connectomes represents an advancement in the field of biological databases, transcending traditional limitations through computational mining and dynamic data incorporation. By leveraging full-text publications, these connectomes offer an enriched, contextually detailed exploration of biological interactions, including both the depth and breadth necessary for decoding complex biological systems. They are not just repositories of information but active platforms for discovery, enabling insights into the intricate interplay of genes and pathways. Moving forward, our focus should remain on expanding accessibility and enhancing data comprehensiveness, ensuring that the *C. elegans* and *Drosophila* Connectomes continue to evolve as indispensable resources in the quest to unravel the complexities of life’s fundamental processes.

## LIMITATION OF THE STUDY

While our approach significantly advances the scope of interaction data available by leveraging computational techniques to mine full-text publications, it is dependent upon the accessibility of these publications. Reliance on publicly accessible or institutionally available literature means that some recent studies, especially those behind paywalls or subject to embargo periods, may not be immediately integrated into our database. This could introduce delays in reflecting the most current research findings and innovations within the connectomes. Furthermore, while automated data extraction techniques offer scalability, they may not always achieve the accuracy of manual curation, potentially affecting the precision of interaction data. Efforts are ongoing to refine these methodologies and expand our access to the latest scientific publications, ensuring our databases not only grow in volume but also in the quality and timeliness of the information they provide. Future directions will include exploring collaborations for wider access to publications and enhancing our algorithms for data mining to mitigate these limitations, continually striving to present the most accurate and comprehensive view of the biological landscapes we aim to model.

## METHODS

### KEY RESOUCES TABLE

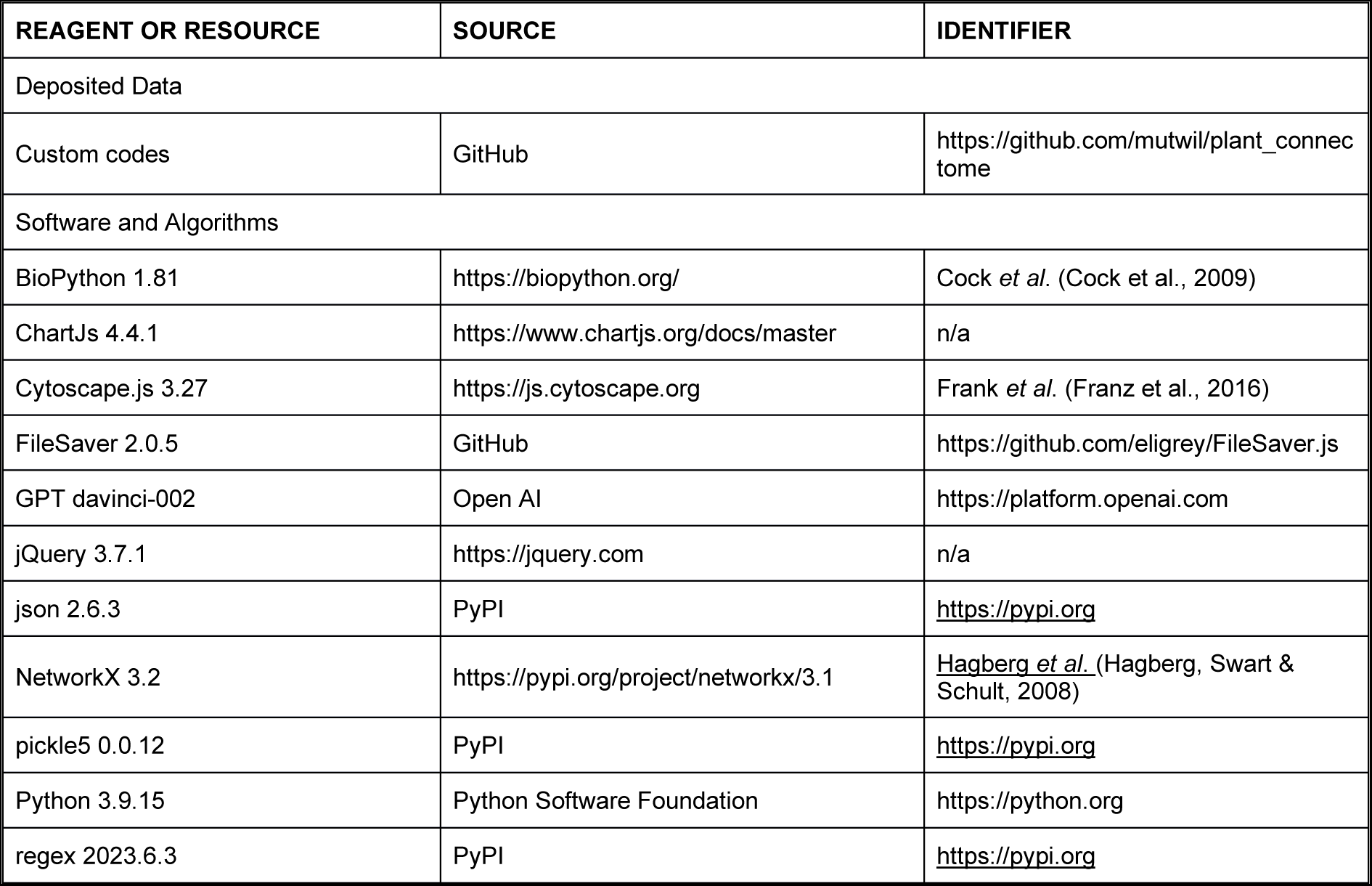

KEY RESOURCES TABLE
| REAGENT OR RESOURCE | SOURCE | IDENTIFIER |
| --- | --- | --- |
| Deposited Data |  |  |
| Custom codes | GitHub | https://github.com/mutwil/plant_connectome |
| Software and Algorithms |  |  |
| BioPython 1.81 | https://biopython.org/ | Cock *et al.* (Cock et al., 2009) |
| ChartJs 4.4.1 | https://www.chartjs.org/docs/master | n/a |
| Cytoscape.js 3.27 | https://js.cytoscape.org | Frank *et al.* (Franz et al., 2016) |
| FileSaver 2.0.5 | GitHub | https://github.com/eligrey/FileSaver.js |
| GPT davinci-002 | Open AI | https://platform.openai.com |
| jQuery 3.7.1 | https://jquery.com | n/a |
| json 2.6.3 | PyPI | https://pypi.org |
| NetworkX 3.2 | https://pypi.org/project/networkx/3.1 | Hagberg *et al.* (Hagberg, Swart & Schult, 2008) |
| pickle5 0.0.12 | PyPI | https://pypi.org |
| Python 3.9.15 | Python Software Foundation | https://python.org |
| regex 2023.6.3 | PyPI | https://pypi.org |

## RESOURCE AVAILABILITY

### Lead Contact

Further information and requests for resources should be directed to and will be fulfilled by the lead contacts, Marek Mutwil, at and Guillaume Thibault, at.

### Material Availability

This study did not generate new reagents.

### Data and Code Availability

The custom codes to generate the connectomes is available at GitHub (https://github.com/mutwil/plant_connectome).

## METHOD DETAILS

### Retrieval of full-text papers

Using BioPython version 1.81, we downloaded all full-text articles freely available in PubMed. This was followed by the acquisition of institutional token access to Elsevier (NTU Library) and by downloading open-access full-texts. Additionally, we included abstracts from PubMed to ensure a robust dataset. The full-texts included titles, abstracts, introductions, results, and discussions. For analysis, each article was processed using OpenAI’s Python API for the davinci 3.5 model, guided by specifically crafted prompts (Table S2). The output underwent further refinement to eliminate single-letter entities (e.g., removing ‘Gene !affects! X’) and to reframe passive edges into active ones (e.g., converting ‘daf-16 ! can restore ! Secretory protein metabolism’ to ‘daf-16 ! restores ! Secretory protein metabolism’). Furthermore, edges with synonymous meanings were consolidated to enhance clarity and consistency. The model operated under default settings, with the exception of setting the temperature parameter to zero, promoting deterministic outcomes. In our final tally, a total of 24,237 articles for the *C. elegans* connectome and 36,372 articles for the *Drosophila* connectome were processed, encompassing both full-texts and abstracts. This comprehensive collection was assembled and analyzed within a span of two weeks, as outlined in the supplementary abstract.

### Construction of *C. elegans* and *Drosophila* Connectome databases

Both connectomes are hosted on a Google Cloud server. The backend was implemented using the Python framework Flask and the Python packages networkx version 3.1, pickle version pickle5 0.0.12, json version 2.6.3, and regex version 2023.6.3. We used JavaScript dependencies jQuery v3.7.1, Cytoscape.js v3.27, ChartJS v4.4.1, and FileSaver v2.0.5 to visualize the KnowledgeNetwork graphs.

### API for *C. elegans* and *Drosophila* Connectomes

*C. elegans* and *Drosophila* Connectomes are equipped with an Application Programming Interface to ease conducting search queries remotely by users. For each successful call to Connectome’s API, a JSON object is returned, containing the functional abbreviations, GO terms, other nodes, and text summaries associated with the search query. To perform searches using the API, users can add “/api/<SEARCH type>/<SEARCH query>” to the web address, where “<SEARCH type>” and “<SEARCH query=” are placeholders representing the type of search and user’s query, respectively.

## ACKNOWLEDGEMENTS

We thank members of Thibault lab for critical reading of the manuscript. This work was supported by funds from the Singapore Ministry of Education Academic Research Fund Tier 1 (RG96/22 to G.T.) and Tier 3 (MOET32022-0002 to M.M.) as well as the Research Scholarship to K.R.A [predoctoral fellowship from Singapore Ministry of Education Academic Research Fund Tier 3 (MOE-MOET32020-0001)].

## Author contributions

Conceptualization: M.M. and G.T.; Methodology: M.M. and G.T.; Formal analysis: K.R.A. and J.W.S.T.; Investigation: K.R.A., J.W.S.T., and M.R.K.; Writing - original draft: G.T. and K.R.A.; Writing - review & editing: K.R.A., M.M., and G.T.; Supervision: E.E.D., M.M. and G.T.; Project administration: M.M. and G.T.; Funding acquisition: M.M. and G.T.

## DECLARATION OF INTERESTS

The authors declare no competing financial interests.

## ADDITIONAL FILES

**Table S1, Related to Figure 1.** List of journals curated for the *C. elegans* and *Drosophila* Connectome. Excel Spreadsheet.

**Table S2, Related to Figure 2.** An example of an abstract, prompts and outputs from GPT.

**Table S3, Related to Figures 3A and 4A**. List of the top 5,000 most frequent edges for the *C. elegans* and *Drosophila* Connectome. Excel Spreadsheet.

**Table S4, Related to Figures 3B and 4B**. List of the top 5,000 most frequent entities for the *C. elegans* and *Drosophila* Connectome. Excel Spreadsheet.

**Table S5, Related to Figures 3C and 4C**. List of the top 5,000 most frequent genes as entities for the *C. elegans* and *Drosophila* Connectome. Excel Spreadsheet.

**Table S6, Related to Figures 3D and 4D**. List of the top 5,000 most frequent edges between genes for the *C. elegans* and *Drosophila* Connectome. Excel Spreadsheet.

**Document S1**. Supplementary File 1.

**Document S2**. Manual accuracy assessment on a random sample of 50 abstracts for the *C. elegans* Connectome.

**Document S3**. Manual accuracy assessment on a random sample of 50 abstracts for the *Drosophila* Connectome.

## Notes

http://worm.connectome.tools

http://drosophila.connectome.tools

https://github.com/mutwil/plant_connectome

### Competing Interest Statement

The authors have declared no competing interest.

### Summary of Updates

Author correction including spelling errors and one author missing in the submitted manuscript file.

