## Supplemental Information for "Harnessing full-text publications for deep insights into *C. elegans* and *Drosophila* connectomes"

**SUPPLEMENTAL TABLES**

**Table S1, Related to Figure 1.** List of journals curated for the *C. elegans* and *Drosophila* Connectomes.

Excel Spreadsheet

**Table S2, Related to Figure 2.** An example of an abstract, prompts and outputs from GPT.

| | |
|---|---|
| Abstract | The insulin/IGF receptor homolog DAF-2 regulates the aging in C. elegans. Decreasing daf-2 activity causes fertile adults to remain active much longer than normal and to live more than twice as long. A more severe decrease in daf-2 function causes young larvae to enter a state of diapause rather than progressing to adulthood. We have asked which cells require daf-2 gene activity in order for the animal to develop to adulthood and to age normally. We found that daf-2 functions cell nonautonomously in both processes. Our findings imply that the life span of C. elegans is determined by a signaling cascade in which the DAF-2 receptor acts in multiple cell lineages to regulate the production or activity of a secondary signal (or signals), which, in turn, controls the growth and longevity of individual tissues in the animal.<br>[Abstract from Apfeld and Kenyon (Apfeld & Kenyon, 1998)] |
| Prompt | "You are data scientist tasked with identifying entities and relationships in scientific text. Please provide only one statement per line, and ensure that each line contains exactly two entities. If a relationship involves more than two entities, please break it down into multiple separate lines".<br><br>For example:<br>Input: "CLas inhibits callose deposition in the sieve pores and the accumulation of reactive oxygen species to favor its cell-to-cell movement. CESA1 interacts with CESA4 and 7. HIS3 mutant inhibits proline production."<br><br>Output: {"CLas: !inhibits! callose deposition\nCLas: !inhibits! accumulation of reactive oxygen species\ncallose deposition: !inhibits! cell-to-cell movement\nCESA1!interacts with!CESA4\nCESA1!interacts with!CESA7\nHIS3 mutant!inhibits!proline production".}<br><br><ABSTRACT><br><br>VERY SHORT, CONCISE SUMMARY CONTAINING ALL INFORMATION WITH TWO ACTORS PER LINE. |
| Output | C.elegans daf-2!regulates!diapause<br>C.elegans daf-2!regulates!life span<br>DAF-2!regulates!aging in C.elegans<br>Decreasing daf-2 activity!causes!longer activity and longer lifespan<br>Severe decrease in daf-2 function!causes!diapause in young larvae<br>daf-2 gene activity!required by!cells for development and normal aging<br>daf-2 functions!cell nonautonomously!in development and aging<br>DAF-2 receptor!acts in!multiple cell lineages |

**Table S3, Related to Figures 3A and 4A**. List of the top 5,000 most frequent edges for the *C. elegans* and *Drosophila* Connectomes.

Excel Spreadsheet

**Table S4, Related to Figures 3B and 4B**. List of the top 5,000 most frequent entities for the *C. elegans* and *Drosophila* Connectomes.

Excel Spreadsheet

**Table S5, Related to Figures 3C and 4C**. List of the top 5,000 most frequent genes as entities for the *C. elegans* and *Drosophila* Connectomes.

Excel Spreadsheet

**Table S6, Related to Figures 3D and 4D**. List of the top 5,000 most frequent edges between genes for the *C. elegans* and *Drosophila* Connectomes.

Excel Spreadsheet

### SUPPLEMENTAL DOCUMENTS

**Document S2, Related to Figures 2A and 2C**. Manual accuracy assessment on a random sample of 50 abstracts for the *C. elegans* Connectome.

**Document S3, Related to Figures 2B and 2D**. Manual accuracy assessment on a random sample of 50 abstracts for the *Drosophila* Connectome.
