## Supplementary material for "Harnessing full-text publications for deep insights into *C. elegans* and *Drosophila* connectomes": Document S2

**Document Supplemental 2**

**Manual accuracy assessment on a random sample of 50 abstracts for the *C. elegans* Connectome**

1. Pubmed ID: 9790527

The insulin/IGF receptor homolog DAF-2 regulates the aging in C. elegans. Decreasing daf-2 activity causes fertile adults to remain active much longer than normal and to live more than twice as long. A more severe decrease in daf-2 function causes young larvae to enter a state of diapause rather than progressing to adulthood. We have asked which cells require daf-2 gene activity in order for the animal to develop to adulthood and to age normally. We found that daf-2 functions cell nonautonomously in both processes. Our findings imply that the life span of C. elegans is determined by a signaling cascade in which the DAF-2 receptor acts in multiple cell lineages to regulate the production or activity of a secondary signal (or signals), which, in turn, controls the growth and longevity of individual tissues in the animal.

Output from GPT:

1. C.elegans daf-2!regulates!diapause
2. C.elegans daf-2!regulates!life span
3. DAF-2!regulates!aging in C.elegans
4. Decreasing daf-2 activity!causes!longer activity and longer lifespan
5. Severe decrease in daf-2 function!causes!diapause in young larvae
6. daf-2 gene activity!required by!cells for development and normal aging
7. daf-2 functions!cell nonautonomously!in development and aging
8. DAF-2 receptor!acts in!multiple cell lineages
9. DAF-2 receptor!regulates!production or activity of secondary signal(s)
10. Secondary signal(s)!controls!growth and longevity of individual tissues in C.elegans

Statements GPT missed:

Correct, Incorrect, Missing:

10, 0, 0

Remarks:

2. Pubmed ID: 9434769

The chaperonin containing TCP-1 (CCT) from the free-living nematode Caenorhabditis elegans was purified and shown to contain at least seven subunit species ranging from 52-65 kDa. SDS-gel electrophoresis and Western blot analyses with antibodies against C. elegans CCT-1 and CCT-5 and an antibody which recognizes a conserved region in vertebrate CCT subunits confirm that the subunit compositions of CCTs from distantly related organisms (C. elegans and bovine species) are remarkably similar. Surprisingly, the co-purified HSP60 chaperonin present in the C. elegans CCT preparation has the greatest binding activity for denatured actin. Expression of a reporter gene under the control of the C. elegans cct-1 promoter is found to be mainly restricted to neuronal and muscle tissues, an observation which is consistent with the participation of CCT in actin and tubulin folding.

Output from GPT:

1. Caenorhabditis elegans chaperonin containing TCP-1!contains!at least seven subunit species
2. Caenorhabditis elegans chaperonin containing TCP-1!purified!
3. gene encoding CCT-1!expresses!reporter gene (misinterpreted by GPT)
4. gene encoding CCT-1!controls!expression in neuronal and muscle tissues
5. CCT-1!similar to!CCT subunits in bovine species
6. CCT-1!participates in!actin and tubulin folding

Statements GPT missed:

Correct, Incorrect, Missing:

5, 1, 0

Remarks:

3. Pubmed ID: 15247923

Temporal control of cell division is critical for proper animal development. To identify mechanisms involved in developmental arrest of cell division, we screened for cell-cycle mutants that disrupt the reproducible pattern of somatic divisions in the nematode C. elegans. Here, we show that the cdc-14 phosphatase is required for the quiescent state of specific precursor cells. Whereas budding yeast Cdc14p is essential for mitotic exit, inactivation of C. elegans cdc-14 resulted in extra divisions in multiple lineages, with no apparent defects in mitosis or cell-fate determination. CDC-14 fused to the green fluorescent protein (GFP-CDC-14) localized dynamically and accumulated in the cytoplasm during G1 phase. Genetic interaction and transgene expression studies suggest that cdc-14 functions upstream of the cki-1 Cip/Kip inhibitor to promote accumulation of CKI-1 in the nucleus. Our data support a model in which CDC-14 promotes a hypophosphorylated and stable form of CKI-1 required for developmentally programmed cell-cycle arrest.

Output from GPT:

1. CDC-14 phosphatase!controls!developmental cell-cycle arrest
2. CDC-14 phosphatase!required for!quiescent state of specific precursor cells
3. C.elegans cdc-14!results in!extra divisions in multiple lineages
4. CDC-14!localizes dynamically and accumulates!in cytoplasm during G1 phase
5. cdc-14!functions upstream of!cki-1 Cip/Kip inhibitor
6. cdc-14!promotes accumulation of!CKI-1 in the nucleus
7. CDC-14!required for!developmentally programmed cell-cycle arrest

Statements GPT missed:

1. Temporal control of cell division is critical for proper animal development.
2. Cdc14p is essential for mitotic exit

Correct, Incorrect, Missing:

7, 0, 2

Remarks:

4. Pubmed ID: 10511554

Ras-mediated signaling is required for induction of vulval cell fates during Caenorhabditis elegans development. By screening for suppressors of the multivulva phenotype caused by constitutively active let-60 ras, we identified the mutation n2527. To clone the gene affected by n2527, we developed a method for high-resolution mapping. We took advantage of the genomic DNA sequence of the N2 strain by using DNA sequencing to scan for single-nucleotide polymorphisms (SNPs) at defined genomic positions of the RC301 strain. An average of one polymorphism per 1.4 kb was detected in predicted intergenic regions. Because of this high frequency, DNA sequencing is an efficient method to scan for SNPs. By alternating between identifying SNPs and mapping n2527 using selected recombinants, we generated an SNP map of progressively higher density. An intensive search for SNPs resulted in a local map with an average marker spacing of approximately 4 kb. This was used to map n2527 to a 9.6-kb interval. The small size of this interval made it feasible to use DNA sequencing to identify the molecular lesion. In principle, this approach can be used for high-resolution mapping of any C. elegans mutation. Furthermore, this approach can be applied to other species as the genomic sequence becomes available. The n2527 mutation affects a previously uncharacterized gene that we named cdf-1, as it encodes a predicted protein with significant similarity to members of the cation diffusion facilitator family.

Output from GPT:

1. local SNP map!used to clone!Caenorhabditis elegans cdf-1
2. Ras-mediated signaling!required for induction of!vulval cell fates
3. Caenorhabditis elegans development!requires!Ras-mediated signaling
4. approach!can be applied to!other species (misinterpreted by GPT)
5. genomic sequence!becomes available!approach can be applied (misinterpreted by GPT)
6. n2527 mutation!affects!previously uncharacterized gene (misinterpreted by GPT)
7. n2527 mutation!affects!cdf-1 gene
8. cdf-1 gene!encodes!predicted protein
9. predicted protein!has similarity to!cation diffusion facilitator family members

Statements GPT missed:

Correct, Incorrect, Missing:

4,5,0

Remarks:

5. Pubmed ID: 15096503

The Ras-extracellular signal-regulated kinase (ERK) cascade is a critical intracellular signaling pathway that regulates growth, survival, and differentiation. Previous work established that Ras-GTP binds to, and facilitates the activation of, the protein kinase Raf-1. Recently, it was demonstrated that the cation diffusion facilitator (CDF) proteins are involved in Ras-ERK signaling by use of a Caenorhabditis elegans genetic screen that identified suppressors of activated Ras. In the current work, we demonstrate that CDF proteins may function downstream of Ras, but upstream of Raf-1 in Xenopus oocytes. We also show that the C. elegans protein CDF-1 and its mammalian homologue ZnT-1 bind to the amino-terminal regulatory portion of Raf-1 and promote the biological and enzymatic activity of Raf-1. Furthermore, we show that Zn(2+) inhibits Raf-1 binding to ZnT-1. We propose a model in which CDF protein binding facilitates Raf-1 activation.

Output from GPT:

1. CDF-1!binds to!amino-terminal regulatory portion of Raf-1
2. ZnT-1!binds to!amino-terminal regulatory portion of Raf-1
3. CDF-1!promotes!biological activity of Raf-1
4. CDF-1!promotes!enzymatic activity of Raf-1
5. ZnT-1!promotes!biological activity of Raf-1
6. ZnT-1!promotes!enzymatic activity of Raf-1
7. Zn(2+)!inhibits!Raf-1 binding to ZnT-1

Statements GPT missed:

1. CDF proteins may be downstream of Ras
2. CDF proteins may be upstream of Raf-1 in Xenopus oocytes.
3. CDF protein binding facilitates Raf-1 activation.

Correct, Incorrect, Missing:

7, 0, 3

Remarks:

6. Pubmed ID: 12204276

The multigene family of intermediate filament (IF) proteins in Caenorhabditis elegans covers 11 members of which four (A1-3, B1) are essential for development. Suppression of a fifth gene (C2) results in a dumpy phenotype. Expression patterns of three essential genes (A1, A3, B1) were already reported. To begin to analyze the two remaining RNAi phenotypes we followed the expression of the A2 and C2 proteins. Expression of A2 mRNA starts in larval stage L1 and continues in the adult. Transgenic A2 promoter/gfp larvae strongly display GFP in the main body hypodermis but not in seam cells. This pattern and the muscle displacement/paralysis induced by RNAi silencing are consistent with the role of this protein in keeping the correct hypodermis/muscle relationship during development. IF protein C2 occurs in the cytoplasm and desmosomes of intestinal cells and in pharynx desmosomes. Expression of C2 starts in the late embryo and persists in all further stages.

Output from GPT:

1. A2 protein!expressed in!larval stage L1
2. A2 protein!expressed in!adult
3. A2 protein!expressed in!main body hypodermis
4. A2 protein!not expressed in!seam cells
5. A2 protein!involved in!keeping correct hypodermis/muscle relationship
6. C2 protein!expressed in!late embryo
7. C2 protein!expressed in!all further stages
8. C2 protein!found in!cytoplasm of intestinal cells
9. C2 protein!found in!desmosomes of intestinal cells
10. C2 protein!found in!pharynx desmosomes
11. Suppression of C2 gene!results in!dumpy phenotype
12. Expression of A1, A3, B1 genes!already reported
13. Expression profiles of A2 and C2 proteins!analyzed

Statements GPT missed:

Correct, Incorrect, Missing:

12, 1, 1

Remarks:

7. Pubmed ID: 32392040

AMP-activated protein kinase (AMPK in human and AAK in <i>C. elegans</i>) is a master regulator of metabolism. It has many isotypes, but its isotype-dependent functions are largely unknown. By developing real-time in-organism NMR metabolomics for <i>C. elegans</i>, we were able to study different roles of†the isotypic catalytic subunits of AAK/AMPK, AAK-1, and AAK-2 in live worms at the whole organism level. The <i>aak-1</i> knockout animals exhibited enhanced glucose production under starvation, strikingly opposite to <i>aak-2</i> knockout animals. Unusually high compensatory expression of the reciprocal isotypes in each KO strain and the results for the double KO animals suggested an unconventional phenotype-genotype relationship and the dominance of <i>aak-2</i> in glucose production. The gene expression patterns showed that the differential phenotypes of <i>aak-1</i> KO strain are due to reduced TCA and glycolysis and enhanced gluconeogenesis compared to the <i>aak-2</i> KO strain. Subsequent <sup>13</sup>C-isotope incorporation experiment showed that the glucose production in <i>aak-1</i> KO occurs through the activation of fatty acid oxidation and glyoxylate shunt. Revealing differential roles of the isotypes of AAK/AMPK, our convenient approach is readily applicable to many <i>C. elegans</i> models for human metabolic diseases.

Output from GPT:

1. AAK-1!enhances!glucose production
2. AAK-2!inhibits!glucose production
3. aak-1!knockout!enhanced glucose production under starvation
4. aak-2!knockout!opposite phenotype to aak-1 knockout
5. reciprocal isotypes!compensatory expression!in each KO strain
6. aak-2!dominant!in glucose production
7. aak-1 KO!reduced TCA and glycolysis!enhanced gluconeogenesis
8. aak-2 KO!enhanced TCA and glycolysis!reduced gluconeogenesis
9. aak-1 KO!activates!fatty acid oxidation and glyoxylate shunt for glucose production

Statements GPT missed:

1. AMP-activated protein kinase (AMPK in human and AAK in <i>C. elegans</i>) is a master regulator of metabolism.

Correct, Incorrect, Missing:

6, 3, 1

Remarks:

8. Pubmed ID: 18050404

TGF-beta superfamily ligands play fundamental roles in the development and physiology of diverse animal species. Genetic and genomic analyses in the model organism Caenorhabditis elegans have contributed to the understanding of TGF-beta-related signal transduction mechanisms. In this chapter, I describe the currently characterized TGF-beta-related signals and signal transduction cassettes in C. elegans. Homology searches of the genome identify five TGF-beta-related genes, for which functions have been identified for three. Two of the TGF-beta-related genes, daf-7 and dbl-1, function through conventional signaling pathways. These signaling pathways are comprised of ser/thr kinase receptors, Smads, and transcription co-factors. A third TGF-beta-related gene, unc-129, functions in axonal guidance using novel signaling mechanisms. Thus, TGF-beta-related signaling in C. elegans proceeds via both conserved and novel paradigms that can inform studies in other animal systems.

Output from GPT:

1. genome!identify!TGF-beta-related genes

2. genome!identify!five genes
3. functions!identified!for three genes
4. TGF-beta-related genes!function through!conventional signaling pathways
5. daf-7!function through!conventional signaling pathways
6. dbl-1!function through!conventional signaling pathways
7. conventional signaling pathways!comprised of!ser/thr kinase receptors
8. conventional signaling pathways!comprised of!Smads
9. conventional signaling pathways!comprised of!transcription co-factors

Statements GPT missed:

1. unc-129, functions in axonal guidance using novel signaling mechanisms.

Correct, Incorrect, Missing:

6, 3, 1.

Remarks:

9. Pubmed ID: 11691930

Members of the basic helix-loop-helix (bHLH) family of transcription factors regulate a wide array of developmental processes in many cell types, including cell fate specification, differentiation and morphogenesis. Our studies describe the cloning of a gene from the nematode Caenorhabditis elegans that is closely related to the vertebrate-activated B-cell factor (ABF) gene. The nematode gene product CeABF-1 was detected by northern blot analysis from RNA isolated from pooled nematodes representing different developmental stages. The developmental expression profile of CeABF-1 was shown by RT-PCR analysis to be predominantly expressed in the larval stages L3 and L4, with lower levels observed in the L2 larval stage and adult. We also show that CeABF-1 is capable of forming heterodimers with E2A proteins and binding E-box target sites. Mammalian cells transfected with CeABF-1 expression plasmids were capable of blocking E2A-mediated gene transcription, but full repression activity required the presence of two conserved amino acid residues found within the first helix of the CeABF-1 bHLH domain. These results suggest a conserved mechanism of gene repression between certain class II bHLH and class I bHLH proteins found in vertebrates and invertebrates.

Output from GPT:

1. gene from nematode Caenorhabditis elegans!is closely related to!vertebrate-activated B-cell factor (ABF) gene
2. CeABF-1!detected by!northern blot analysis
3. CeABF-1!predominantly expressed in!larval stages L3 and L4
4. CeABF-1!is capable of forming!heterodimers with E2A proteins
5. CeABF-1!is capable of binding!E-box target sites
6. CeABF-1!blocks!E2A-mediated gene transcription
7. presence of two conserved amino acid residues!required for!full repression activity
8. conserved mechanism of gene repression!between!class II bHLH and class I bHLH proteins

Statements GPT missed:

1. Members of the basic helix-loop-helix (bHLH) family of transcription factors regulate a wide array of developmental processes in many cell types, including cell fate specification, differentiation and morphogenesis

Correct, Incorrect, Missing:

5, 3, 1

Remarks:

10. Pubmed ID: 16256736

C. elegans SIR-2.1, a member of the Sir-2 family of NAD(+)-dependent protein deacetylases, has been shown to regulate nematode aging via the insulin/IGF pathway transcription factor daf-16. Treatment of C. elegans with the small molecule resveratrol, however, extends life span in a manner fully dependent upon sir-2.1, but independent of daf-16. Microarray analysis of worms treated with resveratrol demonstrates the transcriptional induction of a family of genes encoding prion-like glutamine/asparagine-rich proteins involved in endoplasmic reticulum (ER) stress response to unfolded proteins. RNA interference of abu-11, a member of this ER stress gene family, abolishes resveratrol-mediated life span extension, and overexpression of abu-11 extends the life span of transgenic animals. Furthermore, SIR-2.1 normally represses transcription of abu-11 and other ER stress gene family members, indicating that resveratrol extends life span by inhibiting sir-2.1-mediated repression of ER stress genes. Our findings demonstrate that abu-11 and other members of its ER stress gene family are positive determinants of C. elegans life span.

Output from GPT:

1. Microarray analysis!demonstrates!transcriptional induction of genes
2. resveratrol!treated with!worms
3. genes!encode!prion-like glutamine/asparagine-rich proteins
4. proteins!involved in!ER stress response
5. RNA interference!abolishes!resveratrol-mediated life span extension
6. abu-11!member of!ER stress gene family
7. overexpression of abu-11!extends!life span
8. SIR-2.1!represses!transcription of abu-11 and other ER stress gene family members
9. resveratrol!inhibits!sir-2.1-mediated repression of ER stress genes
10. abu-11 and other members of ER stress gene family!positive determinants of!C.elegans life span

Statements GPT missed:

1. SIR-2.1 regulate nematode aging via the insulin/IGF pathway transcription factor daf-16.

Correct, Incorrect, Missing:

6, 4, 1

Remarks:

11. Pubmed ID: 22372763

Blueberry is rich in bioactive phytochemicals with a wide of range of biological activities and health benefits. However, little is known about their effects on aging. The objectives of this study were to evaluate the effects of supplementation with a blueberry extract (BE) on lifespan and stress resistance using Caenorhabditis elegans (C. elegans) as a model. The mechanisms of these effects were explored using RNAi technology. The mean lifespan of C. elegans treated with BE at 50, 100, and 200 mg mL-1 was significantly increased by 22.2%, 36.5%, and 44.4%, respectively, in a dose-dependent manner. In addition, supplementation with BE improved motility and decreased lipofuscin accumulation. C. elegans pretreated with BE were more resistant than untreated C. elegans to stresses (heat, ultraviolet-B radiation, and paraquat). Treatment with BE resulted in up-regulation of genes related to antioxidant systems, including sod-3, cat-1, mev-1, skn-1, mek-1, nhr-8, and daf-16. Suppression of daf-16 by RNAi shortened the lifespan of C. elegans and inhibited the expression of sod-3, suggesting that BE may regulate sod-3 downstream of daf-16 to extend lifespan and stress resistance. Our findings revealed that, in C. elegans, BE can prolong the lifespan, improve health indexes, and enhance stress resistance.

Output from GPT:

1. Blueberry extract!promotes!longevity

2. Blueberry extract!promotes!stress tolerance
3. Blueberry extract!up-regulation of!genes related to antioxidant systems
4. Blueberry extract!regulates!sod-3 downstream of daf-16
5. RNAi!suppresses!daf-16
6. RNAi!inhibits!expression of sod-3
7. BE!extends!lifespan
8. BE!extends!stress resistance

Statements GPT missed:

1. BE improved motility and decreased lipofuscin accumulation

Correct, Incorrect, Missing:

6, 2, 1

Remarks:

12. Pubmed ID: 18701922

Mammalian neuronal DEG/ENaC channels known as ASICs (acid-sensing ion channels) mediate sensory perception and memory formation. ASICS are closed at rest and are gated by protons. Members of the DEG/ENaC family expressed in epithelial tissues are called ENaCs and mediate Na(+) transport across epithelia. ENaCs exhibit constitutive activity and strict Na(+) selectivity. We report here the analysis of the first DEG/ENaC in Caenorhabditis elegans with functional features of ENaCs that is involved in sensory perception. ACD-1 (acid-sensitive channel, degenerin-like) is constitutively open and impermeable to Ca(2+), yet it is required with neuronal DEG/ENaC channel DEG-1 for acid avoidance and chemotaxis to the amino acid lysine. Surprisingly, we document that ACD-1 is required in glia rather than neurons to orchestrate sensory perception. We also report that ACD-1 is inhibited by extracellular and intracellular acidification and, based on the analysis of an acid-hypersensitive ACD-1 mutant, we propose a mechanism of action of ACD-1 in sensory responses based on its sensitivity to protons. Our findings suggest that channels with ACD-1 features may be expressed in mammalian glia and have important functions in controlling neuronal function.

Output from GPT:

1. DEG-1!is required for!acid avoidance
2. DEG-1!is required for!chemotaxis to lysine
3. ACD-1!is constitutively open
4. ACD-1!is impermeable to Ca(2+)
5. ACD-1!is required in glia!for sensory perception
6. ACD-1!is inhibited by!extracellular acidification
7. ACD-1!is inhibited by!intracellular acidification
8. ACD-1!is sensitive to!protons
9. channels with ACD-1 features!may be expressed in!mammalian glia
10. channels with ACD-1 features!have important functions in!controlling neuronal function

Statements GPT missed:

1. ASICs (acid-sensing ion channels) mediate sensory perception and memory formation.
2. ENaCs exhibit constitutive activity and strict Na(+) selectivity.
3. ASICS are closed at rest and are gated by protons.

Correct, Incorrect, Missing:

10, 0, 3.

Remarks:

13. Pubmed ID: 3272166

In C. elegans, the newly identified ace-3 is the third gene affecting acetylcholinesterase (AChE) activity. ace-3 II specifically affects class C AChE and is unlinked to ace-1 X or ace-2 I, which affect the other two AChE classes (A and B, respectively). Strains homozygous for an ace-3 mutation have no apparent behavioral or developmental defect; ace-1 ace-3 and ace-2 ace-3 double mutants are also nearly wild type. In contrast, ace-1 ace-2 ace-3 triple mutant animals are paralyzed and developmentally arrested; their embryonic development is relatively unimpaired, but they are unable to grow beyond the hatching stage. Based on the analysis of genetic mosaics, we conclude that in the absence of ace-2 and ace-3 function, the expression of ace-1(+) in muscle cells, but not in neurons, is essential for postembryonic viability.

Output from GPT:

1. ace-3!affects!acetylcholinesterase activity
2. ace-3!affects!class C AChE
3. ace-3!unlinked to!ace-1 X
4. ace-3!unlinked to!ace-2 I
5. ace-3 mutation!no apparent behavioral or developmental defect
6. ace-1 ace-3 double mutants!nearly wild type
7. ace-2 ace-3 double mutants!nearly wild type
8. ace-1 ace-2 ace-3 triple mutants!paralyzed and developmentally arrested
9. ace-1 ace-2 ace-3 triple mutants!unable to grow beyond hatching stage
10. ace-1(+) expression in muscle cells!essential for postembryonic viability

Statements GPT missed:

Correct, Incorrect, Missing:

10, 0, 0

Remarks:

14. Pubmed ID: 11580201

Acetylcholinesterase (AChE, EC 3.1.1.7) is responsible for the termination of cholinergic nerve transmission. It is the target of organophosphates and carbamates, two types of chemical pesticides being used extensively in agriculture and veterinary medicine against insects and nematodes. Whereas there is usually one single gene encoding AChE in insects, nematodes are one of the rare phyla where multiple ace genes have been unambiguously identified. We have taken advantage of the nematode Caenorhabditis elegans model to identify the four genes encoding AChE in this species. Two genes, ace-1 and ace-2, encode two major AChEs with different pharmacological properties and tissue repartition: ace-1 is expressed in muscle cells and a few neurons, whereas ace-2 is mainly expressed in motoneurons. ace-3 represents a minor proportion of the total AChE activity and is expressed only in a few cells, but it is able to sustain double null mutants ace-1; ace-2. It is resistant to usual cholinesterase inhibitors. ace-4 was transcribed but the corresponding enzyme was not detected in vivo.

Output from GPT:

1. Caenorhabditis elegans model!identifies!four genes encoding AChE
2. ace-1!expressed in!muscle cells and a few neurons
3. ace-2!expressed in!motoneurons
4. ace-3!expressed in!a few cells
5. ace-3!sustains!double null mutants ace-1; ace-2
6. ace-3!resistant to!usual cholinesterase inhibitors
7. ace-4!transcribed but not detected in vivo

Statements GPT missed:

1. Acetylcholinesterase (AChE, EC 3.1.1.7) is responsible for the termination of cholinergic nerve transmission

Correct, Incorrect, Missing:

7, 0, 1

Remarks:

15. Pubmed ID: 14559923

Genome sequence analyses predict many proteins that are structurally related to proteases but lack catalytic residues, thus making functional assignment difficult. We show that one of these proteins (ACN-1), a unique multi-domain angiotensin-converting enzyme (ACE)-like protein from Caenorhabditis elegans, is essential for larval development and adult morphogenesis. Green fluorescent protein-tagged ACN-1 is expressed in hypodermal cells, the developing vulva, and the ray papillae of the male tail. The hypodermal expression of acn-1 appears to be controlled by nhr-23 and nhr-25, two nuclear hormone receptors known to regulate molting in C. elegans. acn-1(RNAi) causes arrest of larval development because of a molting defect, a protruding vulva in adult hermaphrodites, severely disrupted alae, and an incomplete seam syncytium. Adult males also have multiple tail defects. The failure of the larval seam cells to undergo normal cell fusion is the likely reason for the severe disruption of the adult alae. We propose that alteration of the ancestral ACE during evolution, by loss of the metallopeptidase active site and the addition of new protein modules, has provided opportunities for novel molecular interactions important for post-embryonic development in nematodes.

Output from GPT:

1. ACN-1!essential role in!molting and morphogenesis of Caenorhabditis elegans
2. ACN-1!lacks!metallopeptidase active site
3. proteins!structurally related to proteases!lack catalytic residues
4. ACN-1!essential for!larval development and adult morphogenesis
5. ACN-1!expressed in!hypodermal cells, developing vulva, and ray papillae of male tail
6. acn-1!controlled by!nhr-23 and nhr-25
7. acn-1(RNAi)!causes!arrest of larval development, protruding vulva, disrupted alae, incomplete seam syncytium
8. acn-1(RNAi)!affects!tail development in adult males

Statements GPT missed:

Correct, Incorrect, Missing:

7, 1, 0

Remarks:

16. Pubmed ID: 12507493

Apoptosis, a form of cellular suicide, involves the activation of CED-3-related cysteine proteases (caspases). The regulation of caspases by apoptotic signals and the precise mechanism by which they kill the cell remain unknown. In Drosophila, different death-inducing stimuli induce the expression of the apoptotic activator reaper. Cell killing by reaper and two genetically linked apoptotic activators, hid and grim, requires caspase activity. A Drosophila caspase, named Drosophila caspase-1 (DCP-1), was identified and found to be structurally and biochemically similar to Caenorhabditis elegans CED-3. Loss of zygotic DCP-1 function in Drosophila caused larval lethality and melanotic tumors, showing that this gene is essential for normal development.

Output from GPT:

1. DCP-1!essential for!development
2. Apoptosis!involves activation of!CED-3-related cysteine proteases
3. reaper!requires!caspase activity
4. hid!requires!caspase activity
5. grim!requires!caspase activity
6. DCP-1!similar to!CED-3
7. loss of zygotic DCP-1 function!causes!larval lethality
8. loss of zygotic DCP-1 function!causes!melanotic tumors
9. DCP-1!essential for!normal development

Statements GPT missed:

Correct, Incorrect, Missing:

9, 0, 0

Remarks:

17. Pubmed ID: 35021096

Mitochondria are known as the powerhouse of the cell. Dysfunction of mitochondria homeostasis induces the mitochondrial unfolded protein response (UPR<sup>mt</sup>), altering cellular metabolism. How cells sense the UPR<sup>mt</sup> to rewire metabolism is largely unknown. Here, we show that inactivation of either the citric/tricarboxylic acid (TCA) cycle enzymes aco-2 or idha-1, which encode aconitase and isocitrate dehydrogenase respectively, leads to citrate accumulation. In Caenorhabditis elegans, both in†vitro and in†vivo, citrate accumulation consequently triggers the UPR<sup>mt</sup> and also promotes lipid accumulation. The transcription factor DVE-1 binds to the promoter of the nuclear hormone receptor nhr-80 to transactivate its expression. NHR-80 then upregulates lipogenesis and lipid accumulation, shifting excess citrate for use in lipogenesis and for storage as triacylglycerol in lipid droplets. Inactivation of DVE-1 or NHR-80 fully abolishes the citrate-induced lipid accumulation. Therefore, our work uncovers a DVE-1-NHR-80-lipogenesis axis linking the transmission of the mitochondrial stress signal to lipid metabolism.

Output from GPT:

1. cells!sense!UPRmt
2. cells!rewire metabolism
3. inactivation of aco-2!leads to!citrate accumulation
4. inactivation of idha-1!leads to!citrate accumulation
5. citrate accumulation!triggers!UPRmt
6. citrate accumulation!promotes!lipid accumulation

Statements GPT missed:

1. DVE-1 binds to the promoter of the nuclear hormone receptor nhr-80 to transactivate its expression.
2. Inactivation of DVE-1 or NHR-80 fully abolishes the citrate-induced lipid accumulation.
3. NHR-80 then upregulates lipogenesis and lipid accumulation.

Correct, Incorrect, Missing:

4, 2, 3

Remarks:

18. Pubmed ID: 8581398

A number of putative nicotinic acetylcholine receptor subunit clones were isolated by screening a lambda library of Caenorhabditis elegans genomic DNA with a probe derived from the Drosophila melanogaster ard gene (a non-alpha nicotinic acetylcholine receptor subunit clone). Studies on one of these loci, acr-2, are described; acr-2 is located between sup-7 and unc-6 on the X chromosome. A full-length cDNA was isolated and sequenced. The cDNA encodes a putative non-alpha subunit of a nicotinic acetylcholine receptor that shows many of the conserved features of vertebrate and invertebrate non-alpha nicotinic acetylcholine receptor subunits. To investigate the functional expression of the subunit, the corresponding cRNA was produced, in vitro, and micro-injected into Xenopus oocytes. When expressed alone acr-2 shows no levamisole-gated channel activity. When co-expressed with a C. elegans alpha subunit (unc-38), which is itself unable to form functional homo-oligomers, acr-2 contributed to the formation of a functional channel. This is the first functional expression of a nematode nicotinic acetylcholine receptor and supports the interpretation that the differentiation between alpha and non-alpha subunits dates back to the earliest stages of the evolution of the metazoa.

Output from GPT:

1. acr-2!located between!sup-7 and unc-6
2. acr-2!isolated and sequenced
3. acr-2 cRNA!micro-injected into!Xenopus oocytes
4. acr-2!co-expressed with!unc-38
5. acr-2!contributes to!formation of functional channel
6. Differentiation between alpha and non-alpha subunits!dates back to!earliest stages of metazoa evolution

Statements GPT missed:

1. acr-2 shows no levamisole-gated channel activity

Correct, Incorrect, Missing:

5, 1, 1

Remarks:

19. Pubmed ID: 8617198

ATP binding cassette (ABC) transporters define a family of proteins with strong structural similarities conserved across evolution and devoted to the translocation of a variety of substrates across cell membranes. A few members of the family are known in mammals, but although all of them are medically relevant proteins, knowledge of their molecular function remains scanty. We report here a morphological and functional study of the recently identified mammalian ABC transporter, ABC1. Its expression during embryonic development correlates spatially and temporally with the areas of programmed cell death. More specifically, ABC1 is expressed in macrophages engaged in the engulfment and clearance of dead cells. Moreover, ABC1 transporter is required for engulfment since the ability of macrophages to ingest apoptotic bodies is severely impaired after antibody-mediated steric blockade of ABC1. A structural homologue of ABC1 has been identified in the Caenorhabditis elegans genome and maps close to the ced-7 locus. Since ced-7 phenotype is precisely defined by an imparied engulfment of cell corpses, it is tempting to surmise that ABC1 might be a mammalian homologue of ced-7.

Output from GPT:

1. ABC1 transporter!is required for!engulfment
2. macrophages!ingest!apoptotic bodies
3. antibody-mediated steric blockade!impairs!ingestion of apoptotic bodies
4. ABC1!is a structural homologue of!ABC1 in Caenorhabditis elegans genome
5. ABC1!maps close to!ced-7 locus
6. ced-7 phenotype!is defined by!impaired engulfment of cell corpses
7. ABC1!might be a mammalian homologue of!ced-7

Statements GPT missed:

Correct, Incorrect, Missing:

6, 1, 0

Remarks:

20. Pubmed ID: 9024666

Programmed cell death (apoptosis) is an evolutionarily conserved process used by multicellular organisms to eliminate cells that are not needed or are potentially detrimental to the organism. Members of the Bcl-2 family of mammalian proteins are intimately involved in the regulation of apoptosis, but, their precise mechanism of action remains unresolved. In Caenorhabditis elegans, the Bcl-2 homologue CED-9 prevents cell death by antagonizing the death-promoting activities of CED-3, a member of the Caspase family of death proteases, and of CED-4, a protein with no known mammalian homologue. Here we show that CED-9 interacts physically with CED-4. Mutations that reduce or eliminate CED-9 activity also disrupt its ability to bind CED-4, suggesting that this interaction is important for CED-9 function. Thus, CED-9 might control C. elegans cell death by binding to and regulating CED-4 activity. We propose that mammalian Bcl-2 family members might control apoptosis in a similar way through interaction and regulation of CED-4 homologues or analogues.

Output from GPT:

1. CED-9!interacts with!CED-4
2. Programmed cell death!is regulated by!Bcl-2 family proteins
3. CED-9!prevents!cell death
4. CED-9!antagonizes!CED-3
5. CED-9!antagonizes!CED-4
6. CED-9!binds to!CED-4
7. Mutations in CED-9!disrupt!binding to CED-4
8. CED-9!regulates!CED-4 activity
9. Bcl-2 family members!might control!apoptosis
10. Bcl-2 family members!interact with!CED-4 homologues or analogues

Statements GPT missed:

Correct, Incorrect, Missing:

10, 0, 0

Remarks:

21. Pubmed ID: 15242600

The DNA binding domains of human p53 and Cep-1, its C. elegans ortholog, recognize essentially identical DNA sequences despite poor sequence similarity. We solved the three-dimensional structure of the Cep-1 DNA binding domain in the absence of DNA and compared it to that of human p53. The two domains have similar overall folds. However, three loops, involved in DNA and Zn binding in human p53, contain small alpha helices in Cep-1. The alpha helix in loop L3 of Cep-1 orients the side chains of two conserved arginines toward DNA; in human p53, both arginines are mutation hotspots, but only one contacts DNA. The alpha helix in loop L1 of Cep-1 repositions the entire loop, making it unlikely for residues of this loop to contact bases in the major groove of DNA, as occurs in human p53. Thus, during evolution there have been considerable changes in the structure of the p53 DNA binding domain.

Output from GPT:

1. human p53!has structural differences with!C.elegans ortholog Cep-1
2. DNA binding domains of human p53!recognize!identical DNA sequences
3. Cep-1 DNA binding domain!has similar overall fold with!human p53
4. Cep-1 DNA binding domain!contains alpha helices in!loops involved in DNA and Zn binding
5. Cep-1 alpha helix in loop L3!orients!side chains of conserved arginines towards DNA
6. human p53!has mutation hotspots in!arginines that contact DNA
7. Cep-1 alpha helix in loop L1!repositions!the entire loop, making it unlikely to contact DNA bases
8. evolution!caused considerable changes in the structure of!p53 DNA binding domain

Statements GPT missed:

1. human p53 and Cep-1 are orthologs

Correct, Incorrect, Missing:

7, 1, 1

Remarks:

22. Pubmed ID: 9724641

Expression of a constitutively activated version of the heterotrimeric G protein alpha-subunit Galphas results in the swelling and vacuolization of a specific subset of ventral nerve cord motoneurons of Caenorhabditis elegans. A second site modifier (sgs-1) that completely suppresses this neuronal degeneration has been isolated. sgs-1 was cloned and was shown to encode an adenylyl cyclase which is most similar to mammalian adenylyl cyclase type IX. Mutations in sgs-1 change residues that are conserved among different adenylyl cyclases. These mutations are located in the two catalytic domains and in the first multiple transmembrane spanning region of the predicted protein. An sgs-1 reporter construct shows a general neuronal expression pattern, demonstrating that sgs-1 is expressed in the neurons that are susceptible to activated Galphas-induced cell death. A second C.elegans adenylyl cyclase gene (acy-2) was analyzed as well. In contrast to sgs-1, acy-2 shows a restricted expression pattern and loss of acy-2 function results in early larval lethality. These results suggest that SGS-1 is a target of Galphas signaling in motoneurons, whereas an interaction of Galphas with ACY-2, probably in the canal-associated neurons, is required for viability.

Output from GPT:

1. sgs-1!is expressed in!neurons susceptible to activated Galphas-induced cell death
2. acy-2!shows a restricted expression pattern
3. loss of acy-2 function!results in!early larval lethality
4. SGS-1!is a target of!Galphas signaling in motoneurons
5. interaction of Galphas!with!ACY-2 is required for viability

Statements GPT missed:

1. sgs-1 similar to mammalian adenylyl cyclase type IX.

Correct, Incorrect, Missing:

5, 0, 1

Remarks:

23. Pubmed ID: 10339392

In screens for mutants defective in vulval morphogenesis, multiple mutants were isolated in which the uterus and the vulva fail to make a proper connection. We describe five alleles that define the gene cog-2, for connection of gonad defective. To form a functional connection between the vulva and the uterus, the anchor cell must fuse with the multinucleate uterine seam cell, derived from uterine cells that adopt a (pi) lineage. In cog-2 mutants, the anchor cell does not fuse to the uterine seam cell and, instead, remains at the apex of the vulva, blocking the connection between the vulval and uterine lumens, resulting in an egg-laying defective phenotype. According to lineage analysis and expression assays for two (pi)-cell-specific markers, induction of the (pi) fate occurs normally in cog-2 mutants. We have cloned cog-2 and shown that it encodes a Sox family transcription factor that is expressed in the (pi) lineage. Thus, it appears that COG-2 is a transcription factor that regulates a late-stage aspect of uterine seam cell differentiation that specifically affects anchor cell-uterine seam cell fusion.

Output from GPT:

1. COG-2!necessary for!establishing functional vulval-uterine connection
2. mutants!fail to make!proper connection between uterus and vulva
3. cog-2!encodes!Sox family transcription factor
4. anchor cell!must fuse with!multinucleate uterine seam cell
5. cog-2 mutants!anchor cell does not fuse to!uterine seam cell
6. cog-2 mutants!block!connection between vulval and uterine lumens
7. cog-2!regulates!late-stage aspect of uterine seam cell differentiation
8. COG-2!affects!anchor cell-uterine seam cell fusion

Statements GPT missed:

Correct, Incorrect, Missing:

8, 0, 0

Remarks:

24. Pubmed ID: 22293500

The transmembrane protein MIG-13 is a key regulator required for anterior migration of neural cells in Caenorhabditis elegans, but the signaling mechanisms involved remain unknown. Here, we isolated a suppressor mutation in the unc-71/adm-1 gene, which rescued the AVM neuron migration defect in mig-13 mutants. Genetic analyses revealed that UNC-71 at least partly acts downstream of MIG-13 and has an inhibitory effect on the anterior cell migration. The unc-71 mutation also rescued the anterior migration defect of AVM neuron in src-1 mutants. These findings suggest that MIG-13 controls anteroposterior cell migration by interacting with UNC-71 and SRC-1 in C. elegans.

Output from GPT:

1. MIG-13!controls!anteroposterior cell migration
2. MIG-13!interacts with!UNC-71/ADM-1
3. MIG-13!interacts with!SRC-1
4. MIG-13!is required for!anterior migration of neural cells
5. UNC-71/ADM-1!acts downstream of!MIG-13
6. UNC-71/ADM-1!has an inhibitory effect on!anterior cell migration

Statements GPT missed:

1. unc-71 mutation also rescued the anterior migration defect of AVM neuron in src-1 mutants

Correct, Incorrect, Missing:

6, 0, 1

Remarks:

25. Pubmed ID: 37823127

Human milk oligosaccharides (HMOs) belong to a group of multifunctional glycans that are abundantly present in human breast milk. While health effects of neutral oligosaccharides have been investigated extensively, a lot remains unknown regarding health effects of acidic oligosaccharides, such as the two sialyllactoses (SLs), 3'sialyllactose (3'SL), and 6'sialyllactose (6'SL). We utilized <i>Caenorhabditis elegans</i> (<i>C. elegans</i>) to investigate the effects of SLs on exercise performance. Using swimming as an endurance-type exercise, we found that SLs decrease exhaustion, signifying an increase in endurance that is strongest for 6'SL. Through an unbiased metabolomics approach, we identified changes in energy metabolism that correlated with endurance performance. Further investigation suggested that these metabolic changes were related to adaptations of muscle mitochondria that facilitated a shift from beta oxidation to glycogenolysis during exercise. We found that the effect of SLs on endurance performance required AMPK- (<i>aak-1/aak-2</i>) and adenosine receptor (<i>ador-1</i>) signaling. We propose a model where SLs alter the metabolic status in the gut, causing a signal from the intestine to the nervous system toward muscle cells, where metabolic adaptation increases exercise performance. Together, our results underline the potential of SLs in exercise-associated health and contribute to our understanding of the molecular processes involved in nutritionally-induced health benefits.

Output from GPT:

1. metabolic changes!related to!muscle mitochondria
2. metabolic changes!facilitated shift!beta oxidation to glycogenolysis
3. SLs!require!AMPK signaling
4. SLs!require!adenosine receptor signaling
5. SLs!alter metabolic status!in the gut
6. metabolic status!causes signal!from intestine to nervous system
7. signal!from intestine to nervous system!increases exercise performance

Statements GPT missed:

Correct, Incorrect, Missing:

6, 1, 0

Remarks:

26. Pubmed ID: 12426375

Here we take advantage of the well-characterized and simple nervous system of Caenorhabditis elegans to further our understanding of the functions of RNA editing. We describe the two C.elegans ADAR genes, adr-1 and adr-2, and characterize strains containing homozygous deletions in each, or both, of these genes. We find that adr-1 is expressed in most, if not all, cells of the C.elegans nervous system and also in the developing vulva. Using chemotaxis assays, we show that both ADARs are important for normal behavior. Biochemical, molecular and phenotypic analyses indicate that ADR-1 and ADR-2 have distinct roles in C.elegans, but sometimes act together.

Output from GPT:

1. C.elegans!has!well-characterized and simple nervous system
2. RNA editing!is studied!in C.elegans
3. C.elegans!has!two ADAR genes: adr-1 and adr-2
4. strains!contain!deletions in adr-1 and/or adr-2 genes
5. adr-1!is expressed!in most cells of C.elegans nervous system and developing vulva
6. ADARs!are important!for normal behavior
7. ADR-1 and ADR-2!have distinct roles!in C.elegans
8. ADR-1 and ADR-2!sometimes act together

Statements GPT missed:

Correct, Incorrect, Missing:

6, 2, 0

Remarks:

27. Pubmed ID: 18719245

Adenosine deaminases that act on RNA (ADARs) are editing enzymes that convert adenosine to inosine in double-stranded RNA (dsRNA). ADARs sometimes target codons so that a single mRNA yields multiple protein isoforms. However, ADARs most often target noncoding regions of mRNAs, such as untranslated regions (UTRs). To understand the function of extensive double-stranded 3' UTR structures, and the inosines within them, we monitored the fate of reporter and endogenous mRNAs that include structured 3' UTRs in wild-type Caenorhabditis elegans and in strains with mutations in the ADAR genes. In general, we saw little effect of editing on stability or translatability of mRNA, although in one case an ADR-1 dependent effect was observed. Importantly, whereas previous studies indicate that inosine-containing RNAs are retained in the nucleus, we show that both C. elegans and Homo sapiens mRNAs with edited, structured 3' UTRs are present on translating ribosomes.

Output from GPT:

1. extensive double-stranded 3' UTR structures!monitored!fate of reporter and endogenous mRNAs
2. structured 3' UTRs!included in!wild-type Caenorhabditis elegans
3. mutations in ADAR genes!affected!stability or translatability of mRNA
4. ADR-1!caused!dependent effect
5. inosine-containing RNAs!retained in!nucleus
6. C. elegans!showed!retention of inosine-containing RNAs in the cytoplasm

Statements GPT missed:

Correct, Incorrect, Missing:

3, 3, 0

Remarks:

28. Pubmed ID: 11804572

Retrograde signaling from postsynaptic cells to presynaptic neurons is essential for regulation of synaptic development, maintenance, and plasticity. Here we report that the novel protein AEX-1 controls retrograde signaling at neuromuscular junctions in C. elegans. aex-1 mutants show neural defects including reduced presynaptic activity and abnormal localization of the synaptic vesicle fusion protein UNC-13. Muscle-specific AEX-1 expression rescues these defects but neuron-specific expression does not. AEX-1 has an UNC-13 homologous domain and appears to regulate exocytosis in muscles. This retrograde signaling requires prohormone-convertase function in muscles, suggesting that a peptide is the retrograde signal. This signal regulates synaptic vesicle release via the EGL-30 Gq(alpha) protein at presynaptic terminals.

Output from GPT:

1. AEX-1!controls!retrograde signaling
2. postsynaptic cells!send!retrograde signal
3. presynaptic neurons!receive!retrograde signal
4. AEX-1!regulates!neuromuscular junctions
5. AEX-1 mutants!show!neural defects
6. AEX-1!rescues!defects in muscle
7. AEX-1!does not rescue!defects in neurons
8. AEX-1!regulates!exocytosis in muscles
9. retrograde signaling!requires!prohormone-convertase function
10. muscles!produce!peptide retrograde signal

Statements GPT missed:

1. AEX-1 has an UNC-13 homologous domain

Correct, Incorrect, Missing:

10, 0, 1

Remarks:

29. Pubmed ID: 17488621

Cell fusion is fundamental for reproduction and organ formation. Fusion between most C. elegans epithelial cells is mediated by the EFF-1 fusogen. However, fusion between the anchor cell and the utse syncytium that establishes a continuous uterine-vulval tube proceeds normally in eff-1 mutants. By isolating mutants where the anchor-cell fails to fuse, we identified aff-1. AFF-1 ectopic expression results in fusion of cells that normally do not fuse in C. elegans. The fusogen activity of AFF-1 was further confirmed by its ability to fuse heterologous cells. AFF-1 and EFF-1 differ in their fusogenic activity and expression patterns but share eight conserved predicted disulfide bonds in their ectodomains, including a putative TGF-beta-type-I-Receptor domain. We found that FOS-1, the Fos transcription factor ortholog that controls anchor-cell invasion during nematode development, is a specific activator of aff-1-mediated anchor-cell fusion. Thus, FOS-1 links cell invasion and fusion in a developmental cascade.

Output from GPT:

1. AFF-1!mediates fusion!anchor cell
2. FOS-1!regulates!AFF-1
3. eff-1 mutants!do not affect!fusion between anchor cell and utse syncytium
4. mutants!anchor cell fails to fuse!identified aff-1
5. AFF-1!ectopic expression results in!fusion of cells that normally do not fuse
6. AFF-1!can fuse!heterologous cells
7. AFF-1 and EFF-1!differ in fusogenic activity and expression patterns
8. AFF-1 and EFF-1!share!eight conserved predicted disulfide bonds in their ectodomains
9. FOS-1!activates!aff-1-mediated anchor cell fusion
10. FOS-1!links!cell invasion and fusion in a developmental cascade

Statements GPT missed:

1. Cell fusion is fundamental for reproduction and organ formation.

Correct, Incorrect, Missing:

10, 0, 1

Remarks:

30. Pubmed ID: 21432011

In normal development cell fusion is essential for organ formation and sexual reproduction. The nematode Caenorhabditis elegans has become an excellent system to study the mechanisms and developmental functions of cell-to-cell fusion. In this review we focus on the heterochronic regulation of cell fusion. Heterochronic genes control the timing of specific developmental events in C. elegans. The first microRNAs discovered were found as mutations that affect heterochronic development and cell-cell fusions. In addition numerous heterochronic transcription factors also control specific cell fusion events in C. elegans. We describe what is known about the heterochronic regulation of cell fusion of the epidermal seam cells. The fusogen AFF-1 was previously shown to mediate the fusion of the lateral epidermal seam cells. Here we provide evidence supporting the model in which LIN-29, the heterochronic Zinc-finger transcription factor that controls the terminal fusion of the seam cells, stimulates AFF-1 expression in the seam cells before they fuse. Therefore, the heterochronic gene LIN-29 controls AFF-1-mediated cell-cell fusion as part of the terminal differentiation program of the epidermal seam cells.

Output from GPT:

1. LIN-29!controls!AFF-1-mediated cell-cell fusion
2. LIN-29!stimulates!AFF-1 expression
3. AFF-1!mediates!fusion of lateral epidermal seam cells
4. LIN-29!controls!terminal fusion of seam cells
5. LIN-29!part of!terminal differentiation program of seam cells

Statements GPT missed:

1. Heterochronic genes control the timing of specific developmental events in C. elegans.
2. cell fusion is essential for organ formation and sexual reproduction

Correct, Incorrect, Missing:

5, 0, 2

Remarks:

31. Pubmed ID: 32671785

Senescence is a progressive process of degeneration that occurs when cells and organisms mature. Many studies have shown that autophagy is closely related to senescence. Autophagy gradually decreases with the senescence activity of cells, and vice versa. Therefore, moderate autophagy can protect the body and inhibit cell senescence. The inactivation of genes encoding nematode insulin-like tyrosine kinase receptor (daf-2) inhibited the activity of type I PI3K (age-1), Akt molecules (akt1, akt2), PDK (pdk-1), and TOR, and increased the lifespan and autophagy of Caenorhabditis elegans.

Output from GPT:

1. moderate autophagy!protects the body
2. moderate autophagy!inhibits cell senescence
3. inactivation of daf-2!inhibits activity of age-1
4. inactivation of daf-2!inhibits activity of akt1
5. inactivation of daf-2!inhibits activity of akt2
6. inactivation of daf-2!inhibits activity of pdk-1
7. inactivation of daf-2!inhibits activity of TOR
8. inactivation of daf-2!increases lifespan of Caenorhabditis elegans
9. inactivation of daf-2!increases autophagy of Caenorhabditis elegans

Statements GPT missed:

Correct, Incorrect, Missing:

9, 0, 0

Remarks:

32. Pubmed ID: 9501178

Several genes and partial cDNAs encoding cuticle collagens have been isolated from the sheep parasitic nematode Haemonchus contortus. DNA sequencing and Southern blot hybridization studies reveal that H. contortus collagens comprise a large family of related, but non-identical genes. The genes appear to be dispersed throughout the genome. The predominant size of collagen mRNA in molting worms was found to be between 1.0 and 1.2 kb. The one complete gene that was sequenced contains two short introns and encodes a protein of about 300 amino acids. The predicted protein sequence contain several (Gly-X-Y)n triple helix-coding domains that are interrupted by short stretches of non-helix-coding amino acids. The size of the predicted protein and the organization of the triple-helix coding domains are similar to that of Caenorhabditis elegans collagens. All the H. contortus genes studied show a striking homology to the C. elegans collagen gene subfamily represented by col-1. In particular, the amino acid sequence of the carboxy-terminal non-(Gly-X-Y)n region and the positions of cysteine residues flanking the (Gly-X-Y)n domains were found to be highly conserved in the collagens of these two nematodes.

Output from GPT:

1. contortus genes!show homology to!C.elegans collagen gene subfamily
2. contortus genes!represented by!col-1
3. amino acid sequence!highly conserved in!carboxy-terminal non-(Gly-X-Y)n region
4. cysteine residues!flanking!positions of (Gly-X-Y)n domains
5. collagens!highly conserved in!these two nematodes

Statements GPT missed:

Correct, Incorrect, Missing:

5, 0, 0

Remarks:

33. Pubmed ID: 14757639

The aryl hydrocarbon receptors (AHR) are bHLH-PAS domain containing transcription factors. In mammals, they mediate responses to environmental toxins such as 2, 3, 7, 8-tetrachlorodibenzo-p-dioxin (TCDD). Such functions of AHRs require a cofactor, the aryl hydrocarbon receptor nuclear translocator (ARNT), and the cytoplasmic chaperonins HSP90 and XAP2. AHR homologs have been identified throughout the animal kingdom. We report here that the C. elegans orthologs of AHR and ARNT, ahr-1 and aha-1, regulate GABAergic motor neuron fate specification. Four C. elegans neurons known as RMED, RMEV, RMEL and RMER express the neurotransmitter GABA and control head muscle movements. ahr-1 is expressed in RMEL and RMER neurons. Loss of function in ahr-1 causes RMEL and RMER neurons to adopt a RMED/RMEV-like fate, whereas the ectopic expression of ahr-1 in RMED and RMEV neurons can transform them into RMEL/RMER-like neurons. This function of ahr-1 requires aha-1, but not daf-21/hsp90. Our results demonstrate that C. elegans ahr-1 functions as a cell-type specific determinant. This study further supports the notion that the ancestral role of the AHR proteins is in regulating cellular differentiation in animal development.

Output from GPT:

1. AHR-1!specifies!GABAergic neuron cell fate
2. AHA-1!specifies!GABAergic neuron cell fate
3. ahr-1!regulates!GABAergic motor neuron fate specification
4. aha-1!regulates!GABAergic motor neuron fate specification
5. ahr-1!causes!RMEL and RMER neurons to adopt RMED/RMEV-like fate
6. ahr-1!transforms!RMED and RMEV neurons into RMEL/RMER-like neurons
7. ahr-1!requires!aha-1
8. ahr-1!does not require!daf-21/hsp90

Statements GPT missed:

1. ahr-1 functions as a cell-type specific determinant.
2. ahr-1 is expressed in RMEL and RMER neurons

Correct, Incorrect, Missing:

8, 0, 2

Remarks:

34. Pubmed ID: 16109369

In metazoans, microRNAs (miRNAs) carry out various regulatory functions through association with multiprotein miRNA-induced silencing complexes (miRISCs) that contain Dicer and Argonaute proteins. How miRNAs regulate the expression of their mRNA targets remains a major research question. We have identified the C. elegans ain-1 gene through a genetic suppressor screen and shown that it functions with the heterochronic genetic pathway that regulates developmental timing. Biochemical analysis indicates that AIN-1 interacts with protein complexes containing an Argonaute protein, Dicer, and miRNAs. AIN-1 shares homology with the candidate human neurological disease protein GW182, shown to localize in cytoplasmic processing bodies that are sites of mRNA degradation and storage. A functional AIN-1::GFP also localizes at the likely worm processing bodies. When coexpressed from transgenes, AIN-1 targets ALG-1 to the foci. These results suggest a model where AIN-1 regulates a subset of miRISCs by localization to the processing bodies, facilitating degradation or translational inhibition of mRNA targets.

Output from GPT:

1. AIN-1!interacts with!miRISCs
2. AIN-1!may target!ALG-1
3. AIN-1!functions with!heterochronic genetic pathway
4. AIN-1!interacts with!Argonaute protein, Dicer, and miRNAs
5. AIN-1!shares homology with!GW182
6. AIN-1::GFP!localizes at!processing bodies
7. AIN-1!targets!ALG-1 to foci
8. AIN-1!regulates!subset of miRISCs
9. AIN-1!facilitates degradation or translational inhibition of mRNA targets

Statements GPT missed:

Correct, Incorrect, Missing:

9, 0, 0

Remarks:

35. Pubmed ID: 11694581

Cellular junctions are critical for intercellular communication and for the assembly of cells into tissues. Cell junctions often consist of tight junctions, which form a permeability barrier and prevent the diffusion of lipids and proteins between cell compartments, and adherens junctions, which control the adhesion of cells and link cortical actin filaments to attachment sites on the plasma membrane. Proper tight junction formation and cell polarity require the function of membrane-associated guanylate kinases (MAGUKs) that contain the PDZ protein-protein interaction domain. In contrast, less is known about how adherens junctions are assembled. Here we describe how the PDZ-containing protein DLG-1 is required for the proper formation and function of adherens junctions in Caenorhabditis elegans. DLG-1 is a MAGUK protein that is most similar in sequence to mammalian SAP97, which is found at both synapses of the CNS, as well as at cell junctions of epithelia. DLG-1 is localized to adherens junctions, and DLG-1 localization is mediated by an amino-terminal domain shared with SAP97 but not found in other MAGUK family members. DLG-1 recruits other proteins and signaling molecules to adherens junctions, while embryos that lack DLG-1 fail to recruit the proteins AJM-1 and CPI-1 to adherens junctions. DLG-1 is required for the proper organization of the actin cytoskeleton and for the morphological elongation of embryos. In contrast to other proteins that have been observed to affect adherens junction assembly and function, DLG-1 is not required to maintain cell polarity. Our results suggest a new function for MAGUK proteins distinct from their role in cell polarity.

Output from GPT:

1. DLG-1!is localized to!adherens junctions
2. DLG-1 localization!is mediated by!amino-terminal domain shared with SAP97
3. DLG-1!recruits!other proteins and signaling molecules to adherens junctions
4. embryos that lack DLG-1!fail to recruit!proteins AJM-1 and CPI-1 to adherens junctions
5. DLG-1!is required for!proper organization of actin cytoskeleton
6. DLG-1!is required for!morphological elongation of embryos

Statements GPT missed:

1. DLG-1 is not required to maintain cell polarity.
2. Cell junctions often consist of tight junctions.

Correct, Incorrect, Missing:

6, 0, 2

Remarks:

36. Pubmed ID: 23804750

Duchenne muscular dystrophy (DMD) is a neuromuscular disease caused by mutations in the dystrophin gene. The subcellular mechanisms of DMD remain poorly understood and there is currently no curative treatment available. Using a Caenorhabditis elegans model for DMD as a pharmacologic and genetic tool, we found that cyclosporine A (CsA) reduces muscle degeneration at low dose and acts, at least in part, through a mitochondrial cyclophilin D, CYN-1. We thus hypothesized that CsA acts on mitochondrial permeability modulation through cyclophilin D inhibition. Mitochondrial patterns and dynamics were analyzed, which revealed dramatic mitochondrial fragmentation not only in dystrophic nematodes, but also in a zebrafish model for DMD. This abnormal mitochondrial fragmentation occurs before any obvious sign of degeneration can be detected. Moreover, we demonstrate that blocking/delaying mitochondrial fragmentation by knocking down the fission-promoting gene drp-1 reduces muscle degeneration and improves locomotion abilities of dystrophic nematodes. Further experiments revealed that cytochrome c is involved in muscle degeneration in C. elegans and seems to act, at least in part, through an interaction with the inositol trisphosphate receptor calcium channel, ITR-1. Altogether, our findings reveal that mitochondria play a key role in the early process of muscle degeneration and may be a target of choice for the design of novel therapeutics for DMD. In addition, our results provide the first indication in the nematode that (i) mitochondrial permeability transition can occur and (ii) cytochrome c can act in cell death.

Output from GPT:

1. DMD!poorly understood
2. DMD!no curative treatment available
3. Caenorhabditis elegans model!tool for DMD
4. CsA!reduces muscle degeneration
5. CsA!acts through CYN-1
6. CsA!acts on mitochondrial permeability modulation

Statements GPT missed:

1. knocking down the fission-promoting gene drp-1 reduces muscle degeneration and improves locomotion abilities of dystrophic nematodes.
2. mitochondria play a key role in the early process of muscle degeneration

Correct, Incorrect, Missing:

6, 0, 2

Remarks:

37. Pubmed ID: 8694762

Cyclosporin A (CsA) is the most widely used immunosuppressive agent, whose properties are exerted via an interaction with cyclophilin, resulting in down-regulation of signal-transduction events in the T-cell. Cyclophilin is identical with peptidylprolyl cis-trans isomerase (PPI; EC 5.2.1.8), an enzyme which catalyses the isomerization between the two proline conformations in proteins, thereby acting as a catalyst in protein-folding events. Several reports indicate that CsA has potent anti-parasitic activity, effective against both protozoan and helminth species. In order to understand the various biological roles that cyclophilins play we have initiated a study of these proteins in the genetically tractable nematode Caenorhabditis elegans. Here we describe the cloning and characterization of 11 cyclophilin genes (cyp-1 to -11) derived from this nematode; this is currently the greatest number of isoforms described in a single species. Southern blotting and physical mapping indicated that these genes are dispersed throughout the nematode genome. A high degree of conservation exists between several isoforms, which also share characteristics with the ubiquitous isoforms previously described. The remaining isoforms are divergent, having altered CsA-binding domains and additional non-cyclophilin domains, which may impart compartmental specificity. Ten of these isoforms have been expressed in Escherichia coli, and the resultant fusion proteins have been examined biochemically for PPI activity, which they all possess. Isomerase activity is highest in the conserved and lowest in divergent isoforms, perhaps indicating a more specific substrate for the latter. Analysis of the C. elegans cyp genes will provide answers as to the roles played by cyclophilins in protein folding and signal transduction.

Output from GPT:

1. cyclophilins!play roles!Caenorhabditis elegans
2. study!initiated!cyclophilins, Caenorhabditis elegans
3. cloning!described!11 cyclophilin genes
4. 11 cyclophilin genes!derived from!Caenorhabditis elegans
5. greatest number of isoforms!described in!single species
6. genes!dispersed throughout!nematode genome
7. Southern blotting!indicated!dispersion of genes
8. physical mapping!indicated!dispersion of genes

Statements GPT missed:

1. Cyclosporin A (CsA) is the most widely used immunosuppressive agent

Correct, Incorrect, Missing:

2, 6, 1

Remarks:

38. Pubmed ID: 17276923

The cellular response to genotoxic stress involves the integration of multiple prosurvival and proapoptotic signals that dictate whether a cell lives or dies. In mammals, AKT/PKB regulates cell survival by modulating the activity of several apoptotic proteins, including p53. In Caenorhabditis elegans, akt-1 and akt-2 regulate development in response to environmental cues by controlling the FOXO transcription factor daf-16, but the role of these genes in regulating p53-dependent apoptosis is not known. In this study, we show that akt-1 and akt-2 negatively regulate DNA-damage-induced apoptosis in the C. elegans germline. The antiapoptotic activity of akt-1 is independent of its target gene daf-16 but dependent on cep-1/p53. Although only akt-1 regulates the apoptotic activity of cep-1, both akt-1 and akt-2 modulate the intensity of the apoptotic response independently of the transcriptional activity of CEP-1. Finally, we show that AKT-1 regulates apoptosis but not cell-cycle progression downstream of the HUS-1/MRT-2 branch of the DNA damage checkpoint.

Output from GPT:

1. AKT-1!regulates!DNA-damage-induced germline apoptosis
2. AKT/PKB!regulates!cell survival
3. AKT/PKB!modulates!apoptotic proteins
4. AKT/PKB!modulates!p53
5. akt-1!regulates!development in response to environmental cues
6. akt-1!regulates!FOXO transcription factor daf-16
7. akt-1!regulates!p53-dependent apoptosis
8. akt-1!negatively regulates!DNA-damage-induced apoptosis
9. akt-1!independent of!daf-16
10. akt-1!dependent on!cep-1/p53
11. akt-1!regulates!apoptotic activity of cep-1
12. akt-1 and akt-2!modulate!intensity of apoptotic response
13. akt-1 and akt-2!independently of!transcriptional activity of CEP-1
14. AKT-1!regulates!apoptosis downstream of HUS-1/MRT-2 branch of DNA damage checkpoint

Statements GPT missed:

1. AKT-1 regulates apoptosis

Correct, Incorrect, Missing:

13, 1, 1

Remarks:

39. Pubmed ID: 10200558

ALG-2 is a EF hand calcium binding protein with sequence homologies to calmodulin. Vito et al have shown that ALG-2 expression is required for apoptosis following a number of death stimuli,1 although nothing is known about the effectors which underlie ALG-2 function. Here we have used ALG-2 as bait in a yeast two hybrid screen of a mouse brain cDNA library. We found that ALG-2 binds to itself and to a novel protein that we call ALG-2 interacting protein X, Alix. Using co-immunoprecipitation experiments, we confirmed ALG-2/ALG-2 binding and demonstrated that this interaction is calcium independent. ALG-2/Alix interaction was also validated by co-immunoprecipitation, but in this case, the binding was found to be strictly calcium dependent. Alix seems highly conserved throughout evolution since it shows significant homologies to a putative C. elegans protein (YNK-1) and to proteins of A. nidulans (PalA) and S. cerevisiae (BRO1). Alix is a potential regulator or downstream effector of ALG-2 action.

Output from GPT:

1. Alix!undergoes calcium-dependent interaction with!ALG-2
2. ALG-2!is a EF hand calcium binding protein with sequence homologies to!calmodulin
3. ALG-2 expression!is required for apoptosis following a number of death stimuli
4. ALG-2!binds to itself
5. ALG-2!binds to a novel protein called!ALG-2 interacting protein X, Alix
6. ALG-2/ALG-2 binding!is calcium independent
7. ALG-2/Alix interaction!is calcium dependent
8. Alix!shows significant homologies to a putative C.cerevisiae (BRO1)
9. Alix!is a potential regulator or downstream effector of!ALG-2 action

Statements GPT missed:

1. Alix!shows significant homologies to a putative C. elegans protein (YNK-1) and to proteins of A. nidulans (PalA).

Correct, Incorrect, Missing:

9, 0, 1

Remarks:

40. Pubmed ID: 11461699

RNAi is a gene-silencing phenomenon triggered by double-stranded (ds) RNA and involves the generation of 21 to 26 nt RNA segments that guide mRNA destruction. In Caenorhabditis elegans, lin-4 and let-7 encode small temporal RNAs (stRNAs) of 22 nt that regulate stage-specific development. Here we show that inactivation of genes related to RNAi pathway genes, a homolog of Drosophila Dicer (dcr-1), and two homologs of rde-1 (alg-1 and alg-2), cause heterochronic phenotypes similar to lin-4 and let-7 mutations. Further we show that dcr-1, alg-1, and alg-2 are necessary for the maturation and activity of the lin-4 and let-7 stRNAs. Our findings suggest that a common processing machinery generates guide RNAs that mediate both RNAi and endogenous gene regulation.

Output from GPT:

1. lin-4!regulates!stage-specific development
2. let-7!regulates!stage-specific development
3. dcr-1!inactivation causes!heterochronic phenotypes
4. alg-1!inactivation causes!heterochronic phenotypes
5. alg-2!inactivation causes!heterochronic phenotypes
6. dcr-1!necessary for!maturation and activity of lin-4 and let-7 stRNAs
7. alg-1!necessary for!maturation and activity of lin-4 and let-7 stRNAs
8. alg-2!necessary for!maturation and activity of lin-4 and let-7 stRNAs
9. common processing machinery!generates!guide RNAs for RNAi and endogenous gene regulation

Statements GPT missed:

Correct, Incorrect, Missing:

9, 0, 0

Remarks:

41. Pubmed ID: 15790968

Mutations in the highly conserved Aristaless-related homeodomain protein ARX have been shown to underlie multiple forms of X-linked mental retardation. Arx knockout mice exhibit thinner cerebral cortices because of decreased neural precursor proliferation, and also exhibit defects in the differentiation and migration of GABAergic interneurons. However, the role of ARX in the observed behavioral and developmental abnormalities is unclear. The regulatory functions of individual homeodomain proteins and the networks in which they act are frequently highly conserved across species, although these networks may be deployed in different developmental contexts. In Drosophila, aristaless mutants exhibit defects in the development of terminal appendages, and Aristaless has been shown to function with the LIM-homeodomain protein LIM1 to regulate leg development. Here, we describe the role of the Aristaless/Arx homolog alr-1 in C. elegans. We show that alr-1 acts in a pathway with the LIM1 ortholog lin-11 to regulate the development of a subset of chemosensory neurons. Moreover, we demonstrate that the differentiation of a GABAergic motoneuron subtype is affected in alr-1 mutants, suggesting parallels with ARX functions in vertebrates. Investigating ALR-1 functions in C. elegans may yield insights into the role of this important protein in neuronal development and the etiology of mental retardation.

Output from GPT:

1. C.elegans Aristaless/Arx homolog alr-1!regulates!chemosensory neuron development
2. C.elegans Aristaless/Arx homolog alr-1!regulates!GABAergic motor neuron development
3. ARX!underlies!X-linked mental retardation
4. aristaless mutants!exhibit!defects in terminal appendage development
5. Aristaless!functions with!LIM1 to regulate leg development
6. alr-1!acts in a pathway with!lin-11 to regulate chemosensory neuron development
7. alr-1 mutants!affect!differentiation of GABAergic motoneuron subtype
8. ALR-1 functions!yield insights into!neuronal development and mental retardation etiology

Statements GPT missed:

Correct, Incorrect, Missing:

8, 0, 0

Remarks:

42. Pubmed ID: 35122183

Neurodegenerative diseases with tau pathology, or tauopathies, include Alzheimer's disease and related dementia disorders. Previous work has shown that loss of the poly(A) RNA-binding protein gene sut-2/MSUT2 strongly suppressed tauopathy in Caenorhabditis elegans, human cell culture, and mouse models of tauopathy. However, the mechanism of suppression is still unclear. Recent work has shown that MSUT2 protein interacts with the THO complex and ALYREF, which are components of the mRNA nuclear export complex. Additionally, previous work showed ALYREF homolog Ref1 modulates TDP-43 and G<sub>4</sub>C<sub>2</sub> toxicity in Drosophila melanogaster models. We used transgenic C. elegans models of tau or TDP-43 toxicity to investigate the effects of loss of ALYREF function on tau and TDP-43 toxicity. In C. elegans, three genes are homologous to human ALYREF: aly-1, aly-2, and aly-3. We found that loss of C. elegans aly gene function, especially loss of both aly-2 and aly-3, suppressed tau-induced toxic phenotypes. Loss of aly-2 and aly-3 was also able to suppress TDP-43-induced locomotor behavior deficits. However, loss of aly-2 and aly-3 had divergent effects on mRNA and protein levels as total tau protein levels were reduced while mRNA levels were increased, but no significant effects were seen on total TDP-43 protein or mRNA levels. Our results suggest that although aly genes modulate both tau and TDP-43-induced toxicity phenotypes, the molecular mechanisms of suppression are different and separated from impacts on mRNA and protein levels. Altogether, this study highlights the importance of elucidating RNA-related mechanisms in both tau and TDP-43-induced toxicity.

Output from GPT:

1. C.elegans!has homologous genes to!human ALYREF
2. aly-1, aly-2, aly-3!are homologous genes to!human ALYREF
3. loss of C.elegans aly gene function!suppresses!tau-induced toxic phenotypes
4. loss of aly-2 and aly-3!suppresses!tau-induced toxic phenotypes
5. loss of aly-2 and aly-3!suppresses!TDP-43-induced locomotor behavior deficits
6. loss of aly-2 and aly-3!reduces!total tau protein levels
7. loss of aly-2 and aly-3!increases!mRNA levels
8. loss of aly-2 and aly-3!has no significant effects on!total TDP-43 protein levels
9. loss of aly-2 and aly-3!has no significant effects on!TDP-43 mRNA levels
10. aly genes!modulate!both tau and TDP-43-induced toxicity phenotypes
11. molecular mechanisms of suppression!are different and separated from impacts on!mRNA and protein levels

Statements GPT missed:

1. MSUT2 protein interacts with the THO complex and ALYREF.
2. ALYREF homolog Ref1 modulates TDP-43.

Correct, Incorrect, Missing:

11, 0, 2

Remarks:

43. Pubmed ID: 2586513

Two genomic sequences that share homology with Rp11215, the gene encoding the largest subunit of RNA polymerase II in Drosophila melanogaster, have been isolated from the nematode Caenorhabditis elegans. One of these sequences was physically mapped on chromosome IV within a region deleted by the deficiency mDf4, 25 kilobases (kb) from the left deficiency breakpoint. This position corresponds to ama-1 (resistance to alpha-amanitin), a gene shown previously to encode a subunit of RNA polymerase II. Northern (RNA) blotting and DNA sequencing revealed that ama-1 spans 10 kb, is punctuated by 11 introns, and encodes a 5.9-kb mRNA. A cDNA clone was isolated and partially sequenced to confirm the 3' end and several splice junctions. Analysis of the inferred 1,859-residue ama-1 product showed considerable identity with the largest subunit of RNAP II from other organisms, including the presence of a zinc finger motif near the amino terminus, and a carboxyl-terminal domain of 42 tandemly reiterated heptamers with the consensus Tyr Ser Pro Thr Ser Pro Ser. The latter domain was found to be encoded by four exons. In addition, the sequence oriented ama-1 transcription with respect to the genetic map. The second C. elegans sequence detected with the Drosophila probe, named rpc-1, was found to encode a 4.8-kb transcript and hybridized strongly to the gene encoding the largest subunit of RNA polymerase III from yeast, implicating rpc-1 as encoding the analogous peptide in the nematode. By contrast with ama-1, rpc-1 was not deleted by mDf4 or larger deficiencies examined, indicating that these genes are no closer than 150 kb. Genes flanking ama-1, including two collagen genes, also have been identified.

Output from GPT:

1. ama-1!encodes!largest subunit of Caenorhabditis elegans RNA polymerase II
2. ama-1!mapped on!chromosome IV
3. ama-1!spans!10 kb
4. ama-1!encodes!5.9-kb mRNA
5. ama-1!encoded by!four exons
6. ama-1!shows identity with!largest subunit of RNAP II from other organisms
7. ama-1!oriented with respect to!genetic map
8. rpc-1!encodes!analogous peptide in the nematode
9. rpc-1!not deleted by!mDf4 or larger deficiencies
10. ama-1!flanked by!two collagen genes

Statements GPT missed:

Correct, Incorrect, Missing:

10, 0, 0

Remarks:

44. Pubmed ID: 21055941

Polar body extrusion (PBE) is the specialized asymmetric division by which oocytes accomplish reduction in ploidy and retention of cytoplasm. During maternal gametogenesis, as in male meiosis and mitosis, cytokinesis is accomplished by a ring rich in active Rho, myosin, and formin-nucleated F-actin [1-7]. However, unlike mitosis, wherein the contractile ring encircles the cell equator, the polar body ring assembles as a discoid cortical washer. Here we show that in Caenorhabditis elegans, the meiotic contractile ring transforms during closure from a disc above the spindle to a cylinder around the spindle midzone. The meiotic midbody tube comprises stacked cytoskeletal rings. This topological transition suggests a novel mechanism for constriction of an initially discoid cytokinetic ring. Analysis of mouse PBE indicates that midbody tube formation is a conserved process. Depletion of the scaffold protein anillin (ANI-1) from C. elegans results in large and unstable polar bodies that often fuse with the oocyte. Anillin is dispensable for contractile ring assembly, initiation, and closure but is required for the meiotic contractile ring to transform from a disc into a tube. We propose that cytoskeletal bundling by anillin promotes formation of the midbody tube, which ensures the fidelity of PBE.

Output from GPT:

1. mouse PBE!indicates!midbody tube formation
2. scaffold protein anillin (ANI-1)!depletion results in!large and unstable polar bodies
3. scaffold protein anillin (ANI-1)!depletion results in!polar bodies fusion with oocyte

Statements GPT missed:

1. Anillin is dispensable for contractile ring assembly, initiation, and closure.
2. Anillin is required for the meiotic contractile ring to transform from a disc into a tube.

Correct, Incorrect, Missing:

3, 0, 2

Remarks:

45. Pubmed ID: 12640035

Migration of cells through the reorganization of the actin cytoskeleton is essential for morphogenesis of multicellular animals. In a cell culture system, the actin-related protein (Arp) 2/3 complex functions as a nucleation core for actin polymerization when activated by the members of the WASP (Wiskott-Aldrich syndrome protein) family. However, the regulation of cell motility in vivo remains poorly understood. Here we report that homologues of the mammalian Arp2/3 complex and N-WASP in Caenorhabditis elegans play an important role in hypodermal cell migration during morphogenesis, a process known as ventral enclosure. In the absence of one of any of the C. elegans Arp2/3 complex subunits (ARX-1, ARX-2, ARX-4, ARX-5, ARX-6 or ARX-7) or of N-WASP (WSP-1), hypodermal cell migration led by actin-rich filopodia formation is inhibited during ventral enclosure owing to the reduction of filamentous actin formation. However, there is no effect on differentiation of hypodermal cells and dorsal intercalation. Disruption of the function of ARX-1 and WSP-1 in hypodermal cells also resulted in hypodermal cell arrest during ventral enclosure, suggesting that their function is cell autonomous. WSP-1 protein activated Arp2/3-mediated actin polymerization in vitro. Consistent with these results, the Arp2/3 complex and WSP-1 colocalized at the leading edge of migrating hypodermal cells. The stable localization of WSP-1 was dependent on the presence of Arp2/3 complex, suggesting an interaction between the Arp2/3 complex and WSP-1 in vivo.

Output from GPT:

1. C.elegans Arp2/3 complex subunits!inhibit!hypodermal cell migration
2. N-WASP!inhibits!hypodermal cell migration
3. hypodermal cell migration!inhibited by!reduction of filamentous actin formation
4. ARX-1!disrupts!hypodermal cell migration
5. WSP-1!disrupts!hypodermal cell migration
6. ARX-1!disrupts!hypodermal cell arrest
7. WSP-1!disrupts!hypodermal cell arrest
8. ARX-1!disrupts!cell autonomous function
9. WSP-1!disrupts!cell autonomous function
10. WSP-1!activates!Arp2/3-mediated actin polymerization

Statements GPT missed:

Correct, Incorrect, Missing:

7, 3, 0

Remarks:

46. Pubmed ID: 17223323

The developmental timing of all types of cells must be synchronized and spatially coordinated to achieve the organized development of a multicellular organism. Previously, we found RNAi of asb-1, encoding a germline-specific isoform of mitochondrial ATP synthase b subunit, caused 100% penetrant sterility in Caenorhabditis elegans. ATP synthase is one of the five complexes of the mitochondrial respiratory chain, and defects in some of the components of the chain are known to slow the growth and extend the lifespan of worms. We found that development of asb-1 mutant germ line was not arrested at any stage, but did slow to half the rate of wild type, whereas the rate of somatic development was the same in asb-1 mutants as that of wild type, indicating that asb-1 is required to maintain the rate of germline development but has no effect on somatic development. Among ATP synthase subunit genes, RNAi of asg-1, encoding a germline-specific isoform of the g subunit, also caused asb-1-like sterility, indicating that some other germline-specific components are also required to maintain the rate of germline development. Both asb-1 and asg-1 are located on autosomes while they possess counterparts, asb-2 and asg-2, respectively, on X chromosome, which are both required for somatic development. Chromosomal locations of the genes may be the basis of the segregation of germline/somatic functions of each gene, as were demonstrated for other autosomal/X-linked duplicated gene pairs.

Output from GPT:

1. ASB-1!is required to maintain!rate of germline development
2. ASB-1!is required to maintain!rate of germline development
3. RNAi of asb-1!caused!100% penetrant sterility
4. ATP synthase!is one of!five complexes of mitochondrial respiratory chain
5. Defects in components of mitochondrial respiratory chain!slow growth and extend lifespan of worms
6. Development of asb-1 mutant germ line!slows to half the rate!of wild type
7. ASB-1!has no effect on!somatic development
8. RNAi of asg-1!caused!asb-1-like sterility
9. ASB-1 and ASG-1!are located on!autosomes
10. ASB-2 and ASG-2!are located on!X chromosome
11. ASB-2 and ASG-2!are required for!somatic development
12. Chromosomal locations of genes!may be the basis of!segregation of germline/somatic functions

Statements GPT missed:

Correct, Incorrect, Missing:

12, 0, 0

Remarks:

47. Pubmed ID: 26234645

Caenorhabditis elegans BLMP-1 is the homolog of mammalian PRDM1/BLIMP1 transcriptional repressor and a component of the heterochronic pathway regulating developmental timing. We found that BLMP-1 positively and directly regulates the bed-3 gene required for vulval cell division and molting. blmp-1 mutation or RNAi reduces bed-3 reporter expression and causes phenotype similar to bed-3 mutations. We mapped an enhancer element responsible for bed-3 expression in the vulva and the hypodermis to a 200 bp region in the third intron. Using EMSA, we identified BLMP-1 binding sites within this region. Mutating these sites abolished both in vitro BLMP-1 binding and in vivo enhancer activity. Thus, BLMP-1 directly and positively regulates bed-3 transcription, connecting the heterochronic pathway to regulation of vulval cell division and molting. Recently, C. elegans blmp-1 was found to act as a transcriptional repressor. Thus, C. elegans BLMP-1 acts as a transcriptional activator or a repressor depending on the context.

Output from GPT:

1. PRDM1/BLIMP1 ortholog BLMP-1!directly regulates!Caenorhabditis elegans bed-3
2. BLMP-1!homolog of!mammalian PRDM1/BLIMP1 transcriptional repressor
3. BLMP-1!component of!heterochronic pathway regulating developmental timing
4. blmp-1 mutation or RNAi!reduces!bed-3 reporter expression
5. blmp-1 mutation or RNAi!causes!phenotype similar to bed-3 mutations
6. Enhancer element!responsible for!bed-3 expression in vulva and hypodermis
7. Enhancer element!mapped to!200 bp region in third intron
8. EMSA!identified!BLMP-1 binding sites within enhancer region
9. Mutating binding sites!abolished!in vitro BLMP-1 binding and in vivo enhancer activity
10. BLMP-1!directly and positively regulates!bed-3 transcription
11. BLMP-1!connects!heterochronic pathway to regulation of vulval cell division and molting

Statements GPT missed:

1. C. elegans BLMP-1 acts as a transcriptional activator or a repressor depending on the context

Correct, Incorrect, Missing:

11, 0, 1

Remarks:

48. Pubmed ID: 17135259

A mutation in the Caenorhabditis elegans bre-1 gene was isolated in a screen for Bacillus thuringiensis toxin-resistant (bre) mutants to the Cry5B crystal toxin made by B. thuringiensis. bre-1 mutant animals are different from the four other cloned bre mutants in that their level of resistance is noticeably lower. bre-1 animals also display a significantly reduced brood size at 25 degrees C. Here we cloned the bre-1 gene and characterized the bre-1 mutant phenotype. bre-1 encodes a protein with significant homology to a GDP-mannose 4,6-dehydratase, which catalyzes the first step in the biosynthesis of GDP-fucose from GDP-mannose. Injection of GDP-fucose but not fucose into C. elegans intestinal cells rescues bre-1 mutant phenotypes. Thus, C. elegans lacks a functional fucose salvage pathway. Furthermore, we demonstrate that bre-1 mutant animals are defective in production of fucosylated glycolipids and that bre-1 mutant animals make quantitatively reduced levels of glycolipid receptors for Cry5B. We finally show that bre-1 mutant animals, although viable, show a lack of fucosylated N- and O-glycans, based on mass spectrometric evidence. Thus, C. elegans can survive with little fucose and can develop resistance to crystal toxin by loss of a monosaccharide biosynthetic pathway.

Output from GPT:

1. Resistance to Bacillus thuringiensis toxin: !caused by! loss of fucose
2. Caenorhabditis elegans: !mutant! in bre-1 gene
3. Caenorhabditis elegans bre-1 mutant animals: !show! lower level of resistance
4. Caenorhabditis elegans bre-1 mutant animals: !display! reduced brood size at 25 degrees C
5. bre-1 gene: !encodes! protein with homology to GDP-mannose 4,6-dehydratase
6. GDP-mannose 4,6-dehydratase: !catalyzes! first step in biosynthesis of GDP-fucose
7. Injection of GDP-fucose: !rescues! bre-1 mutant phenotypes in C. elegans intestinal cells
8. C. elegans: !lacks! functional fucose salvage pathway
9. bre-1 mutant animals: !defective in! production of fucosylated glycolipids
10. bre-1 mutant animals: !make! reduced levels of glycolipid receptors for Cry5B
11. bre-1 mutant animals: !lack! fucosylated N- and O-glycans

Statements GPT missed:

1. C. elegans can survive with little fucose.
2. C. elegans can develop resistance to crystal toxin by loss of a monosaccharide biosynthetic pathway.

Correct, Incorrect, Missing:

10, 1, 2

Remarks:

49. Pubmed ID: 16079230

Interactions with bacteria play a major role in immune responses, ecology, and evolution of all animals, but they have been neglected until recently in the case of C. elegans. We report a genetic investigation of the interaction of C. elegans with the nematode-specific pathogen Microbacterium nematophilum, which colonizes the rectum and causes distinctive tail swelling in its host. A total of 121 mutants with altered response to infection were isolated from selections or screens for a bacterially unswollen (Bus) phenotype, using both chemical and transposon mutagenesis. Some of these correspond to known genes, affecting either bacterial adhesion or colonization (srf-2, srf-3, srf-5) or host swelling response (sur-2, egl-5). Most mutants define 15 new genes (bus-1-bus-6, bus-8, bus-10, bus-12-bus-18). The majority of these mutants exhibit little or no rectal infection when challenged with the pathogen and are probably altered in surface properties such that the bacteria can no longer infect worms. A number have corresponding alterations in lectin staining and cuticle fragility. Most of the uninfectable mutants grow better than wild type in the presence of the pathogen, but the sur-2 mutant is hypersensitive, indicating that the tail-swelling response is associated with a specific defense mechanism against this pathogen.

Output from GPT:

1. srf-2, srf-3: affect bacterial adhesion
2. srf-5: affects bacterial colonization
3. sur-2, egl-5: affect host swelling response
4. bus-1-bus-6, bus-8, bus-10, bus-12-bus-18: define 15 new genes
5. mutants: exhibit little or no rectal infection
6. mutants: altered in surface properties
7. bacteria: can no longer infect worms

Statements GPT missed:

1. sur-2 mutant is hypersensitive

Correct, Incorrect, Missing:

6, 1, 1

Remarks:

50. Pubmed ID: 15914661

The pharynx of Caenorhabditis elegans is a tubular muscle controlled by its own set of neurons. We developed a technique to voltage clamp the pharyngeal muscle and demonstrate by analyzing mutants that the pharyngeal action potential is regulated by three major voltage-gated currents, conducted by a T-type calcium channel CCA-1, an L-type calcium channel EGL-19 and a potassium channel EXP-2. We show that CCA-1 exhibits T-type calcium channel properties: activation at -40 mV and rapid inactivation. Our results suggest that CCA-1's role is to accelerate the action potential upstroke in the pharyngeal muscle in response to excitatory inputs. Similarly to other L-type channels, EGL-19 activates at high voltages and inactivates slowly; thus it may maintain the plateau phase of the action potential. EXP-2 is a potassium channel of the kV family that shows inward rectifier properties when expressed in Xenopus laevis oocytes. We show that endogenous EXP-2 is not a true inward rectifier--it conducts large outward currents at potentials up to +20 mV and is therefore well suited to trigger rapid repolarization at the end of the action potential plateau phase. Our results suggest that EXP-2 is a potassium channel with unusual properties that uses a hyperpolarization threshold to activate a regenerative hyperpolarizing current.

Output from GPT:

1. CCA-1!shapes!action potentials
2. EGL-19!shapes!action potentials
3. EXP-2!shapes!action potentials
4. CCA-1!conducted by!T-type calcium channel
5. EGL-19!conducted by!L-type calcium channel
6. EXP-2!conducted by!potassium channel
7. CCA-1!regulates!pharyngeal action potential
8. EGL-19!regulates!pharyngeal action potential
9. CCA-1!accelerates!action potential upstroke
10. EGL-19!maintains!plateau phase of action potential

Statements GPT missed:

1. EXP-2 is a potassium channel of the kV family that shows inward rectifier properties when expressed in Xenopus laevis oocytes.

Correct, Incorrect, Missing:

7, 3, 1

Remarks:
