## Supplementary material for "Harnessing full-text publications for deep insights into *C. elegans* and *Drosophila* connectomes": Document S3

**Document Supplemental 3**

**Manual accuracy assessment on a random sample of 50 abstracts for the *Drosophila* Connectome**

1. Pubmed ID: 19323850

The host range of major fruit fly pests in Central Tanzania was evaluated from October 2004 to October 2006. Samples of 48 potential hosts were collected and incubated for fruit fly emergence. Bactrocera invadens was the dominant species in incidence expressed as the ratio of infested to total number samples collected, as well as infestation rate, expressed as number of flies emerging per unit weight. Eight new host fruits are reported. Infestation by native pests, such as Ceratitis capitata and C. cosyra, was minor compared to B. invadens. Ceratitis rosa was the dominant species in temperate fruits, and Cucurbitaceae were mainly infested by Bactrocera cucurbitae, a specialized cucurbit feeder. Among commercial fruits, high infestation incidences were observed in mango and guava, but they decreased throughout the fruiting season. Low infestation rates were observed in all Citrus species and in avocado, indicating these fruits as poor hosts for the studied fruit fly pests in this region. Widespread availability and abundance of fruit species studied here ensures year-round breeding of B. invadens. Seasonal infestation differs, with mango being the most important host in October to January, while guava being important from February to August. Tropical almond showed very high incidence and infestation rate for B. invadens and might act as an important reservoir host, bridging the fruiting seasons of mango and guava. Soursop acts as an important host for C. cosyra after the mango season. Ceratitis capitata is a pest of minor importance of the commercial fruits studied in this region.

Output from GPT:

1. invadens.Ceratitis rosa!dominant species!temperate fruits

2. Cucurbitaceae!infested by!Bactrocera cucurbitae

3. commercial fruits!high infestation incidences!mango

4. commercial fruits!high infestation incidences!guava

5. infestation incidences!decreased!fruiting season

Statements GPT missed:

1. Low infestation rates were observed in all Citrus species and in avocado

2. Soursop acts as an important host for C. cosyra after the mango season

Correct, incorrect, missing:

1, 4, 2

Remarks:

Statement 1 is wrong. GPT read B.invadens together with Ceratitis rosa.

Statements 3 and 4 are poorly interpreted.

Statement 5 is wrong. Infestation incidences decreased THROUGHOUT fruiting season.

Missing statements are likely from GPT not having been given that portion of the abstract.

2. Pubmed ID: 24896539

Discrimination of particular species within the species complexes of tephritid fruit flies is a very challenging task. In this fruit-fly family, several complexes of cryptic species have been reported, including the African cryptic species complex (FAR complex). Cuticular hydrocarbons (CHCs) appear to be an excellent tool for chemotaxonomical discrimination of these cryptic species. In the present study, CHC profiles have been used to discriminate among three important agricultural pests from the FAR complex, Ceratitis fasciventris, Ceratitis anonae and Ceratitis rosa. Hexane body surface extracts of mature males and females were analyzed by two-dimensional gas chromatography with mass spectrometric detection and differences in CHC profiles between species and sexes tested through multivariate statistics and compared with species identification by means of microsatellite markers. Quantitative as well as qualitative CHC profile differences between sexes and species are reported. The CHC profiles consisted of a mixture of linear, internally methyl-branched and mono-, di- and tri-unsaturated alkanes. Twelve compounds were pinpointed as potential chemotaxonomical markers. The present study shows that presence or absence of particular CHCs might be used in the chemical diagnosis of the FAR complex. Moreover, our results represent an important first step in the development of a useful chemotaxonomic tool for cryptic species identification of these important agricultural pests.

Output from GPT:

1. anonae!is discriminated from!Ceratitis fasciventris

2. anonae!is discriminated from!Ceratitis rosa

3. Ceratitis fasciventris!is discriminated from!Ceratitis rosa

4. Differences in CHC profiles!are tested!between species and sexes

5. Cuticular hydrocarbons (CHCs)!are used to discriminate!anonae, Ceratitis fasciventris, and Ceratitis rosa

6. CHC profiles!are analyzed!in mature males and females

7. CHC profiles!are compared with!species identification by microsatellite markers

8. Quantitative and qualitative CHC profile differences!are reported!between sexes and species

Statements GPT missed:

Correct, incorrect, missing:

8, 0, 0

Remarks:

Statements 1,2 and 5 are partially correct and should have included “Ceratitis” for anonae.

Missing statements are likely from GPT not having been given that portion of the abstract.

3. Pubmed ID: 19326842

The effect of expression of the enzyme producing nitric oxide on Drosophila melanogaster morphogenesis was studied using ectopic expression of a truncated splice form of the enzyme that has a dominant negative effect on the full-length enzyme form. The suppressed enzyme expression was shown to affect the cell polarity and movement as well as the structural pattern of organs during Drosophila development. The effect of nitric oxide production on the intercellular distribution of significant factors of cell polarity, protein products of the Strabismus and Prospero genes, has been revealed.

Output from GPT:
1. enzyme expression!affects!cell polarity

2. enzyme expression!affects!structural pattern of organs

3. nitric oxide production!affects!intercellular distribution

4. enzyme expression!affects!cell movement

5. nitric oxide production!affects!protein products of Strabismus gene

6. nitric oxide production!affects!protein products of Prospero gene

Statements GPT missed:

1. effect of nitric oxide production on the intercellular distribution of significant factors of cell polarity

Correct, incorrect, missing:

3, 3, 1

Remarks:

Enzyme was never named in the abstract, should have been nitric oxide synthase. GPT missed the enzyme name for statements 1, 2 and 4.

Statements 3, 5 and 6 are partially should have been read together. (e.g. nitric oxide production!affects!intercellular movement of protein products of Strabismus gene).

4. Pubmed ID: 17333936

General transcription factors (GTFs) were tested for the presence on the promoter of the Drosophila melanogaster hsp70 gene in vivo. TBP, TBP-associated TAF proteins, TFIIB, TFIIF (RAP30), TFIIH (XPB), the TFTC complex (GCN5 and TRRAP), and a Mediator complex subunit (MEDI 3) were detected on the promoter before heat induction. Heat exposure significantly reduced the contents of TBP, TAF proteins, TFIIB, and TFIIF (RAP30), while these proteins were detected in ecdysone--ucible loci. It was assumed on the basis of these findings that a special mechanism induces transcription from the hsp70 promoter and that the apparent presence or absence of GTFs does not always reflect the transcriptional status of a gene.

Output from GPT:

1. GTFs!tested for presence on promoter!hsp70 gene

Statements GPT missed:

1. TBP, TBP-associated TAF proteins, TFIIB, TFIIF (RAP30), TFIIH (XPB), the TFTC complex (GCN5 and TRRAP), and a Mediator complex subunit (MEDI 3) were detected on the promoter before heat induction

2. Heat exposure significantly reduced the contents of TBP, TAF proteins, TFIIB, and TFIIF (RAP30) and these proteins were detected in ecdysone--ucible loci

Correct, incorrect, missing:

1, 0, 2

Remarks:

Missing statements are likely from GPT not having been given that portion of the abstract.

5. Pubmed ID: 1876836

Recombinant cDNA clones that encode two distinct subunits of the transcription factor GA binding protein (GABP) have been isolated. The predicted amino acid sequence of one subunit, GABP alpha, exhibits similarity to the sequence of the product of the ets-1 protooncogene in a region known to encompass the Ets DNA binding domain. The sequence of the second subunit, GABP beta, contains four 33-amino acid repeats located close to the NH2-terminus of the subunit. The sequences of these repeats are similar to repeats in several transmembrane proteins, including Notch from Drosophila melanogaster and Glp-1 and Lin-12 from Caenorhabditis elegans. Avid, sequence-specific binding to DNA required the presence of both polypeptides, revealing a conceptual convergence of nuclear transforming proteins and membrane-anchored proteins implicated in developmentally regulated signal transduction processes.

Output from GPT:

1. GABP beta!contains!four 33-amino acid repeats

2. GABP beta!located close to!NH2-terminus of the subunit

3. repeats!similar to!repeats in several transmembrane proteins

4. repeats!similar to!Notch from Drosophila melanogaster

5. repeats!similar to!Lin-12 from Caenorhabditis elegans

6. binding to DNA!requires!presence of both polypeptides

7. nuclear transforming proteins!converge with!membrane-anchored proteins

8. nuclear transforming proteins!implicated in!developmentally regulated signal transduction processes

9. repeats!similar to!Glp-1 from Caenorhabditis elegans

Statements GPT missed:

1. The predicted amino acid sequence of one subunit, GABP alpha, exhibits similarity to the sequence of the product of the ets-1 protooncogene in a region known to encompass the Ets DNA binding domain.

Correct, incorrect, missing:

8, 1, 1

Remarks:

6. Pubmed ID: 2105491

From a group of 436 Drosophila melanogaster cDNA clones, we selected 39 that are expressed exclusively or predominantly in the adult visual system. By sequence analysis, 20 of the clones appear to represent previously unreported distinct cDNAs. The corresponding transcripts are detected in the retina and optic lobes. The genes are scattered throughout the genome, some near mutations known to affect eye function. One of these clones has been identified, by sequence analysis, as the structural gene (Arr) for a Drosophila homolog of human arrestin. Vertebrate arrestin interacts with rhodopsin in phototransduction and has been associated with an autoimmune form of uveitis in primates. The presence of an arrestin homolog in Drosophila suggests that both the vertebrate and invertebrate phototransduction cascades are regulated in a similar manner.

Output from GPT:

1. Drosophila visual system cDNA clones!include!homolog of human arrestin

2. Vertebrate arrestin!interacts with!rhodopsin in phototransduction

3. 436 Drosophila melanogaster cDNA clones!selected as!expressed in adult visual system

4. 39 selected clones!scattered throughout!genome

5. Some clones!near!mutations affecting eye function

6. One clone identified as!structural gene (Arr)!for Drosophila homolog of human arrestin

7. Presence of arrestin homolog in Drosophila!suggests!similar regulation of phototransduction cascades

Statements GPT missed:

1. Vertebrate arrestin has been associated with an autoimmune form of uveitis in primates

Correct, incorrect, missing:

4, 3, 1

Remarks:

7. Pubmed ID: 10727946

The soluble N-ethylmaleimide-sensitive-factor-attachment proteins (SNAP) are eukaryotic soluble proteins required for membrane fusion. Based on their initial identification in bovine brain cytosol, they are divided in alpha/beta and gamma subfamilies. SNAPs act as adapters between N-ethylmaleimide-sensitive factor (NSF), a hexameric ATPase, and membrane SNARE proteins (SNAP receptors). Within the NSF/SNAP/SNARE complex, SNAPs contribute to the catalysis of an ATP-driven conformational change in the SNAREs, resulting in dissociation of the complex. We have constructed a Dictyostelium discoideum strain overexpressing a c-myc-tagged form of D. discoideum NSF (NSF-myc). Its immunoprecipitation from detergent-solubilized membrane extracts reveals two associated polypeptides with apparent molecular masses of 33 and 36 kDa (p33 and p36) that are absent in NSF-myc immunoprecipitates from cytosol. Analysis of trypsin-digested peptides by microsequencing and mass spectrometry and comparison with cDNA sequences identify p33 and p36 as the D. discoideum homologues of alpha- and gamma-SNAP, respectively. The alpha-/gamma-SNAP molar ratio is close to 3 in vegetative amoebae from this organism. The molecular identification of gamma-SNAP in plants (Arabidopsis thaliana) and insects (Drosophila melanogaster) documents, for the first time, the wide distribution of the gamma subtype. Altogether, these results suggest a specific role for gamma-SNAP, distinct from that of alpha-SNAP.

Output from GPT:

1. bovine brain cytosol!divided into!gamma subfamilies

2. SNAPs!act as adapters between!N-ethylmaleimide-sensitive factor (NSF)

3. SNAPs!act as adapters between!membrane SNARE proteins (SNAP receptors)

4. Dictyostelium discoideum strain!overexpressing!c-myc-tagged form of D

5. Dictyostelium discoideum strain!overexpressing!NSF-myc

6. bovine brain cytosol!divided into!alpha/beta subfamilies

7. SNAPs!contribute to!catalysis of ATP-driven conformational change

8. SNAPs!contribute to!dissociation of NSF/SNAP/SNARE complex

9. NSF-myc!immunoprecipitation reveals!p33 and p36 polypeptides

10. p33 and p36!identified as!D. discoideum proteins

Statements GPT missed:

1. SNAPs are eukaryotic soluble proteins SNAPs are required for membrane fusion

2. N-ethylmaleimide-sensitive factor (NSF) is a hexameric ATPase

Correct, incorrect, missing:

3, 7, 6

Remarks:

Statements 1 and 6 is wrong. It is the SNAPs that are divided into subfamilies. Additionally alpha beta and gamma subfamilies are the 3 families, not alpha/beta and gamma as 2 families.

Statements 2 and 3 should be one statement. SNAPs adapt between NSF and SNAREs.

Statements 4 and 5 should be one statement.

Statement 7 is partially correct and should include that the conformational change occurs in SNAREs.

8. Pubmed ID: 10483093

We present a global analysis of the distribution of 43 transposable elements (TEs) in 228 species of the Drosophila genus from our data and data from the literature. Data on chromosome localization come from in situ hybridization and presence/absence of the elements from southern analyses. This analysis shows great differences between TE distributions, even among closely related species. Some TEs are distributed according to the phylogeny of their host species; others do not entirely follow the phylogeny, suggesting horizontal transfers. A higher number of insertion sites for most TEs in the genome of D. melanogaster is observed when compared with that in D. simulans. This suggests either intrinsic differences in genomic characteristics between the two species, or the influence of differing effective population sizes, although biases due to the use of TE probes coming mostly from D. melanogaster and to the way TEs are initially detected in species cannot be ruled out. Data on TEs more specific to the species under consideration are necessary for a better understanding of their distribution in organisms and populations.

Output from GPT:

1. transposable elements!distributed in!Drosophila species

2. some TEs!do not follow!phylogeny, suggesting horizontal transfers

3. differences!in genomic characteristics!between the two species

4. analysis!shows differences!in TE distributions

5. some TEs!follow phylogeny!of host species

6. higher number of insertion sites!in D.melanogaster!compared with D

7. data on species-specific TEs!necessary!for better understanding of distribution

Statements GPT missed:

Correct, incorrect, missing:

6, 1, 0

Remarks:

Statement 6 misses the full species name.

9. Pubmed ID: 28197917

The ubiquitin -like protein SUMO is conjugated covalently to hundreds of target proteins in organisms throughout the eukaryotic domain. Genetic and biochemical studies using the model organism Drosophila melanogaster are beginning to reveal many essential functions for SUMO in cell biology and development. For example, SUMO regulates multiple signaling pathways such as the Ras/MAPK, Dpp, and JNK pathways. In addition, SUMO regulates transcription through conjugation to many transcriptional regulatory proteins, including Bicoid, Spalt , Scm, and Groucho. In some cases, conjugation of SUMO to a target protein inhibits its normal activity, while in other cases SUMO conjugation stimulates target protein activity. SUMO often modulates a biological process by altering the subcellular localization of a target protein. The ability of SUMO and other ubiquitin-like proteins to diversify protein function may be critical to the evolution of developmental complexity.

Output from GPT:

1. SUMO!is conjugated to!target proteins

2. SUMO!modulates!biological process

3. SUMO!diversifies!protein function

4. SUMO!is critical to!evolution of developmental complexity

5. SUMO!alters!subcellular localization of target protein

Statements GPT missed:

1. SUMO regulates multiple signaling pathways such as the Ras/MAPK, Dpp, and JNK pathways

2. SUMO regulates transcription through conjugation to many transcriptional regulatory proteins, including Bicoid, Spalt , Scm, and Groucho

Correct, incorrect, missing:

5, 0, 2

Remarks:

10. Pubmed ID: 6235226

The DNA polymerase activity of the near homogeneous, multisubunit DNA polymerase-primase from Drosophila melanogaster embryos has been compared to Escherichia coli DNA polymerase III core, DNA polymerase III, and DNA polymerase III holoenzyme. The rate of deoxynucleotide incorporation by the Drosophila polymerase on singly primed phi X174 DNA is similar to that observed with equivalent levels of DNA polymerase III holoenzyme in the absence of E. coli single-stranded DNA binding protein. However, analysis of the DNA products indicates that the Drosophila polymerase is less processive than DNA polymerase III holoenzyme, and closely resembles DNA polymerase III. The Drosophila polymerase-primase contains neither 3'-5' exonuclease nor RNase H-like activities, and catalyzes no significant pyrophosphate exchange. There is a low level of DNA-dependent ATPase activity which can be eliminated by a second glycerol gradient sedimentation (Kaguni, L.S., Rossignol, J.-M., Conaway, R.C., and Lehman, I.R. (1983) Proc. Natl. Acad. Sci. U. S. A. 80, 2221-2225). Although lacking a 3'-5' exonuclease, the replication fidelity of the D. melanogaster polymerase is similar to that of E. coli DNA polymerase III holoenzyme which possesses such an activity.

Output from GPT:

1. Drosophila polymerase-primase: !contains! 3'-5' exonuclease

2. Drosophila polymerase-primase: !contains! RNase H-like activities

3. Drosophila polymerase-primase: !catalyzes! no significant pyrophosphate exchange

4. Drosophila polymerase-primase: !has! low level of DNA-dependent ATPase activity

5. DNA-dependent ATPase activity: !can be eliminated by! second glycerol gradient sedimentation

Statements GPT missed:

1. replication fidelity of the D. melanogaster polymerase is similar to that of E. coli DNA polymerase III holoenzyme

Correct, incorrect, missing:

3, 2, 1

Remarks:

Statements 1 and 2 are wrong. The primase does not contain either.

11. Pubmed ID: 2120680

The molecular organization of the dunce gene of Drosophila melanogaster has proved to be particularly complex, with two divergently transcribed genes, Sgs-4 and Pig-1, nested within its 79 kb intron (1). Here we report the identification and the molecular characterization of a third gene nested within the transcription unit of dunce. This newly identified gene is located nearly 6 kb downstream Pig-1, within a more upstream dunce intron. The gene is developmentally regulated and transcribed with the same polarity of dunce; several lines of evidence indicate that it might encode for a salivary gland secreted (Sgs) protein.

Output from GPT:

1. dunce gene!contains!new gene

2. new gene!located within!upstream dunce intron

3. dunce gene!contains!Sgs-4 gene

4. dunce gene!contains!Pig-1 gene

5. new gene!located within!dunce transcription unit

6. new gene!might encode for!salivary gland secreted protein

Statements GPT missed:

1. located nearly 6 kb downstream Pig-1

Correct, incorrect, missing:

6, 0, 1

Remarks:

12. Pubmed ID: 6417663

Recombinant plasmids in which the sequence encoding the bacterial chloramphenicol acetyltransferase (CAT; acetyl-CoA:chloramphenicol 3-O-acetyltransferase, EC 2.3.1.28) has been placed under the control of Drosophila heat shock protein 70 (hsp 70) or copia promoters have been introduced into cultured cells of two Drosophila species (Schneider II line of Drosophila melanogaster and D. immigrans) as calcium-phosphate complexes. Within 1-2 days after transfection functional CAT enzyme was detected in cells exposed to either CAT recombinant. The expression of the bacterial information depends on the activity of the Drosophila promoters because plasmids in which the Drosophila DNA fragments were fused to the CAT coding sequence in inverted orientation did not support the synthesis of CAT enzyme activity. Low levels of CAT activity and of hybrid mRNA were detected in cells transformed with hsp-cat recombinants when the cells were maintained at room temperature, and both mRNA levels and CAT activity increased substantially after a brief exposure to 37 degrees C. hsp-cat mRNA has the same 5' terminus as authentic Drosophila hsp 70 messenger. These experiments document a practical system for the introduction and expression of isolated genes in cultured cells of Drosophila.

Output from GPT:

1. Transient expression of genes!introduced into!cultured cells of Drosophila

2. Recombinant plasmids!placed under the control of!Drosophila heat shock protein 70 (hsp 70) or copia promoters

3. Recombinant plasmids!introduced into!cultured cells of two Drosophila species

4. Plasmids!fused to!CAT coding sequence

5. Drosophila promoters!control the expression of!bacterial information

6. mRNA levels and CAT activity!increased after!exposure to 37 degrees C

7. hsp-cat mRNA!has the same 5' terminus as!Drosophila hsp 70 messenger

8. Practical system!for the introduction and expression of!isolated genes in cultured cells of Drosophila

9. Low levels of CAT activity and hybrid mRNA!detected in!cells transformed with hsp-cat recombinants at room temperature

Statements GPT missed:

Correct, incorrect, missing:

6, 3, 0

Remarks:

Not sure where GPT got ‘transient’ for statement 1.

Statements 2 and 3 are partially correct and could be combined into one statement.

13. Pubmed ID: 1843280

Furin, the translational product of the recently discovered fur gene, appears to be the first known mammalian member of the subtilisin family of serine proteases and the first known mammalian proprotein-processing enzyme with cleavage selectivity for paired basic amino acid residues. Structurally and functionally, it resembles the prohormone-processing enzyme, kexin (EC 3.4.21.61), which is encoded by the KEX2 gene of yeast Saccharomyces cerevisiae. Most likely, furin is primarily involved in the processing of precursors of proteins that are secreted via the constitutive secretory pathway. Here, we review the discovery of the fur gene and describe the isolation of cDNA clones corresponding to human and mouse fur and to two fur-like genes of Drosophila melanogaster, Dfur1 and Dfur2. We also compare the structural organization of the various deduced furin proteins to that of yeast kexin, and of other members of the subtilisin family of serine proteases. Furthermore, the biosynthesis of biologically active human and mouse furin is evaluated. Finally, the cleavage specificity for paired basic amino acid residues of human and mouse furin is demonstrated by the correct processing of the precursor for von Willebrand factor.

Output from GPT:

1. cDNA clones!correspond to!mouse fur

2. furin proteins!have structural organization similar to!yeast kexin

3. furin!is involved in!processing of precursors of secreted proteins

4. cDNA clones!correspond to!human fur

5. cDNA clones!correspond to!Dfur1 gene of Drosophila melanogaster

6. cDNA clones!correspond to!Dfur2 gene of Drosophila melanogaster

7. furin proteins!have structural organization similar to!other subtilisin family serine proteases

Statements GPT missed:

1. Furin is a mammalian member of the subtilisin family of serine proteases

2. Furin functionally resembles kexin

Correct, incorrect, missing:

3, 4, 2

Remarks:

Statement 3 is partially correct and should include that the secretion is done via the constitution secretory pathway.

Statement 7 is wrong. The text only explained that the structures were compared but did not state anything about their actual similarity.

14. Pubmed ID: 20870963

How functional diversification affects the organization of the transcriptome is a central question in systems genetics. To explore this issue, we sequenced all six Odorant binding protein (Obp) genes located on the X chromosome, four of which occur as a cluster, in 219 inbred wild-derived lines of Drosophila melanogaster and tested for associations between genetic and phenotypic variation at the organismal and transcriptional level. We observed polymorphisms in Obp8a, Obp19a, Obp19b, and Obp19c associated with variation in olfactory responses and polymorphisms in Obp19d associated with variation in life span. We inferred the transcriptional context, or "niche," of each gene by identifying expression polymorphisms where genetic variation in these Obp genes was associated with variation in expression of transcripts genetically correlated to each Obp gene. All six Obp genes occupied a distinct transcriptional niche. Gene ontology enrichment analysis revealed associations of different Obp transcriptional niches with olfactory behavior, synaptic transmission, detection of signals regulating tissue development and apoptosis, postmating behavior and oviposition, and nutrient sensing. Our results show that diversification of the Obp family has organized distinct transcriptional niches that reflect their acquisition of additional functions.

Output from GPT:

1. functional diversification!affects!organization of the transcriptome

2. functional diversification!affects!transcriptional niches

3. transcriptome!contains!six Odorant binding protein (Obp) genes

4. Obp genes!located on!X chromosome

5. 219 inbred wild-derived lines!sequenced for!genetic and phenotypic variation

6. polymorphisms in Obp8a, Obp19a, Obp19b, and Obp19c!associated with!olfactory responses

7. all six Obp genes!occupy!distinct transcriptional niches

8. four Obp genes!occur as!cluster

9. polymorphisms in Obp19d!associated with!life span variation

10. genetic variation in Obp genes!associated with!variation in expression of correlated transcripts

11. different Obp transcriptional niches!associated with!olfactory behavior, synaptic transmission, tissue development, apoptosis, postmating behavior, oviposition, and nutrient sensing

12. diversification of Obp family!organized!distinct transcriptional niches

Statements GPT missed:

Correct, incorrect, missing:

10, 2, 0

Remarks:

Statement 5 is wrong as the text never stated if sequencing was done.

15. Pubmed ID: 11520357

We cloned and sequenced a full length cDNA coding for [Arg7]-corazonin in the greater wax moth Galleria mellonella. The deduced corazonin preprohormone consists of a nineteen amino acid signal peptide, the actual eleven amino acid corazonin sequence, followed by a Gly serving for amidation, a Lys-Arg processing site and an eighty amino acid corazonin precursor-related peptide. The data confirm the phylogenetic conservation of the actual corazonin sequence. The signal peptide and the precursor-related peptide exhibit a similar spacing of a few amino acids as detected in the corazonin preprohormone of Drosophila melanogaster. Northern blots and in situ hybridization experiments revealed that the G. mellonella corazonin gene is tissue-specifically expressed in four pairs of lateral neurosecretory cells in the brains of penultimate and last instar larvae, as well as of pupae and adults. No corazonin mRNA was detected in other cells of the nervous system, fat body, gut, and several other organs.

Output from GPT:

1. We!cloned and sequenced!full length cDNA coding for [Arg7]-corazonin

2. Corazonin preprohormone!consists of!nineteen amino acid signal peptide

3. Corazonin preprohormone!consists of!eleven amino acid corazonin sequence

4. Corazonin preprohormone!consists of!Gly serving for amidation

5. Corazonin preprohormone!consists of!Lys-Arg processing site

6. Corazonin preprohormone!consists of!eighty amino acid corazonin precursor-related peptide

7. Data!confirm!phylogenetic conservation of actual corazonin sequence

Statements GPT missed:

1. G. mellonella corazonin gene is tissue-specifically expressed in four pairs of lateral neurosecretory cells in the brains of penultimate and last instar larvae, as well as of pupae and adults

Correct, incorrect, missing:

7, 0, 1

Remarks:

Missing statements are likely from GPT not having been given that portion of the abstract.

16. Pubmed ID: 24748742

Objectives: The antimutagenic effect of caffeine is evaluated against ethyl methanesulfonate (EMS)-induced mutation rate in Drosophila.

Materials and methods: The mutation rate is evaluated using wing mosaic assay. In transheterozygous larvae, multiple wing hair (mwh 0.3-3) and flare (flr 3-38.8) genes were used as markers of the extent of mutagenicity.

Results: The results at 0.5 and 1.0 mM EMS concentration at both 48 ± 4 and 72 ± 4 h have shown consistent increase in mutation rate, which was being measured as frequency of clone formation per 105 cells. Toxicity of caffeine at 5 mM concentration was parallel to that of distilled water alone. At 0.5 mM EMS concentration at 42 ± 4 and 72 ± 4 h, Drosophila larvae mutation rate was significantly increased. Although caffeine prevented mutation rate in all pre, post, and combined treatment, it was more significant in pretreatment experiments where it was found to be effective in reducing the genotoxicity of EMS. However, the concentration of caffeine as recommended in dietary allowance did not induce the frequency of mutant clones in somatic mutation and recombination test (SMART) recorded.

Conclusion: This study shows that caffeine significantly reduced the genotoxicity induced by EMS. However, the limitation in completely abolishing genotoxicity induced by EMS as observed at the dietary allowance of caffeine makes it interesting for further in-depth study. Further studies on the molecular mechanism of antigenotoxic effect of caffeine will also be interesting.

Output from GPT:

1. mutation rate!evaluated using!wing mosaic assay

2. transheterozygous larvae!used as markers of!extent of mutagenicity

3. results!shown consistent increase in!mutation rate

4. results!measured as!frequency of clone formation per 105 cells

5. multiple wing hair genes!used as markers of!extent of mutagenicity

6. flare genes!used as markers of!extent of mutagenicity

7. EMS concentration!at 0.5 and 1.0 mM!results in consistent increase in mutation rate

8. EMS concentration!at 48 ± 4 and 72 ± 4 h!results in consistent increase in mutation rate

Statements GPT missed:

1. caffeine significantly reduced the genotoxicity induced by EMS

Correct, incorrect, missing:

2, 2, 1

Remarks:

Statement 2 is wrong. The mwh and flare genes are used as markers, not the larvae. (Statement 5 & 6)

Statement 4 is wrong. The mutation rate is measured as a frequency of clone formation, not the results.

17. Pubmed ID: 9891050

The genes of the trithorax group (trxG) in Drosophila melanogaster are required to maintain the pattern of homeotic gene expression that is established early in embryogenesis by the transient expression of the segmentation genes. The precise role of each of the diverse trxG members and the functional relationships among them are not well understood. Here, we report on the isolation of the trxG gene moira (mor) and its molecular characterization. mor encodes a fruit fly homolog of the human and yeast chromatin-remodeling factors BAF170, BAF155, and SWI3. mor is widely expressed throughout development, and its 170-kDa protein product is present in many embryonic tissues. In vitro, MOR can bind to itself and it interacts with Brahma (BRM), an SWI2-SNF2 homolog, with which it is associated in embryonic nuclear extracts. The leucine zipper motif of MOR is likely to participate in self-oligomerization; the equally conserved SANT domain, for which no function is known, may be required for optimal binding to BRM. MOR thus joins BRM and Snf5-related 1 (SNR1), two known Drosophila SWI-SNF subunits that act as positive regulators of the homeotic genes. These observations provide a molecular explanation for the phenotypic and genetic relationships among several of the trxG genes by suggesting that they encode evolutionarily conserved components of a chromatin-remodeling complex.

Output from GPT:

1. MOR!SANT domain!required for optimal binding to BRM

2. mor!encodes!fruit fly homolog of BAF170

3. mor!encodes!fruit fly homolog of BAF155

4. mor!encodes!fruit fly homolog of SWI3

5. mor!170-kDa protein product!present in embryonic tissues

6. MOR!leucine zipper motif!participates in self-oligomerization

Statements GPT missed:

1. mor is widely expressed throughout development

2. MOR can bind to itself and it interacts with Brahma (BRM), an SWI2-SNF2 homolog.

3. BRM and SNR1 act as positive regulators of the homeotic genes

Correct, incorrect, missing:

6, 0, 3

Remarks:

Statements 2, 3 and 4 could have also mentioned that those were yeast chromatin-remodelling factors.

18. Pubmed ID: 15479229

Mitochondrial biogenesis is a complex and highly regulated process that requires the controlled expression of hundreds of genes encoded in two separated genomes, namely the nuclear and mitochondrial genomes. To identify regulatory proteins involved in the transcriptional control of key nuclear-encoded mitochondrial genes, we have performed a detailed analysis of the promoter region of the alpha subunit of the Drosophila melanogaster F1F0 ATP synthase complex. Using transient transfection assays, we have identified a 56 bp cis-acting proximal regulatory region that contains binding sites for the GAGA factor and the alcohol dehydrogenase distal factor 1. In vitro mutagenesis revealed that both sites are functional, and phylogenetic footprinting showed that they are conserved in other Drosophila species and in Anopheles gambiae. The 56 bp region has regulatory enhancer properties and strongly activates heterologous promoters in an orientation-independent manner. In addition, Northern blot and RT-PCR analysis identified two alpha-F1-ATPase mRNAs that differ in the length of the 3' untranslated region due to the selection of alternative polyadenylation sites.

Output from GPT:

1. 56 bp region!has!regulatory enhancer properties

2. 56 bp region!activates!heterologous promoters

3. Northern blot!identified!two alpha-F1-ATPase mRNAs

4. two alpha-F1-ATPase mRNAs!differ in length!3' untranslated region

5. two alpha-F1-ATPase mRNAs!due to!selection of alternative polyadenylation sites

Statements GPT missed:

Correct, incorrect, missing:

4, 1, 0

Remarks:

Statement 2 could have also included that the activation is done in an orientation-independent manner.

Statement 3 failed to include the RT-PCR.

Statement 5 should have been part of statement 4.

Missing statements are likely from GPT not having been given that portion of the abstract.

19. Pubmed ID: 7631747

We have examined the process of membrane protein targeting in the polarized cells of the developing Drosophila melanogaster embryo. Human placental alkaline phosphatase (PLAP) is a glycosylphosphatidyl inositol-linked protein that accumulates at the apical membranes of mammalian epithelial cells. A chimeric construct composed of the transmembrane and cytosolic portions of the vesicular stomatitis virus G protein coupled to the ectodomain of PLAP, termed PLAPG, has been found to behave as a basolateral protein (D. A. Brown, B. Crise, and J. K. Rose. Science Wash. DC 232: 34-47, 1989). The subcellular distributions of these proteins were examined in the epithelial and neuronal tissues of transgenic Drosophila embryos. In the surface ectoderm, both PLAP and PLAPG were restricted to the basolateral membranes throughout development. Internal epithelia derived from the surface ectoderm accumulated PLAP at their apical surfaces, whereas PLAPG retained its basolateral distribution. The redistribution of PLAP from the basolateral to the apical plasma membrane was found to be coincident with the invagination of the surface epithelium to form internal structures, suggesting that the sorting pathways that function in the epithelium of the Drosophila embryo are developmentally regulated.

Output from GPT:

1. developing Drosophila melanogaster embryo!examined!process of membrane protein targeting

2. Human placental alkaline phosphatase (PLAP)!accumulates at!apical membranes of mammalian epithelial cells

3. PLAPG!retains!basolateral distribution

4. chimeric construct (PLAPG)!behaves as!basolateral protein

5. transgenic Drosophila embryos!examined!subcellular distributions of PLAP and PLAPG

6. surface ectoderm!restricts PLAP and PLAPG!to basolateral membranes

7. internal epithelia!accumulate PLAP!at apical surfaces

8. redistribution of PLAP!coincident with!invagination of surface epithelium

9. sorting pathways!function in!epithelium of Drosophila embryo

Statements GPT missed:

1. PLAP is a glycosylphosphatidyl inositol-linked protein

Correct, incorrect, missing:

7, 3, 1

Remarks:

Statement 3 could have included that the PLAPG was derived from surface ectoderm.

20. Pubmed ID: 18246067

In both yeast and mammals, uncapped telomeres activate the DNA damage response (DDR) and undergo end-to-end fusion. Previous work has shown that the Drosophila HOAP protein, encoded by the caravaggio (cav) gene, is required to prevent telomeric fusions. Here we show that HOAP-depleted telomeres activate both the DDR and the spindle assembly checkpoint (SAC). The cell cycle arrest elicited by the DDR was alleviated by mutations in mei-41 (encoding ATR), mus304 (ATRIP), grp (Chk1) and rad50 but not by mutations in tefu (ATM). The SAC was partially overridden by mutations in zw10 (also known as mit(1)15) and bubR1, and also by mutations in mei-41, mus304, rad50, grp and tefu. As expected from SAC activation, the SAC proteins Zw10, Zwilch, BubR1 and Cenp-meta (Cenp-E) accumulated at the kinetochores of cav mutant cells. Notably, BubR1 also accumulated at cav mutant telomeres in a mei-41-, mus304-, rad50-, grp- and tefu-dependent manner. Our results collectively suggest that recruitment of BubR1 by dysfunctional telomeres inhibits Cdc20-APC function, preventing the metaphase-to-anaphase transition.

Output from GPT:

1. SAC!partially overridden by!zw10

2. SAC!partially overridden by!bubR1

3. SAC!partially overridden by!mei-41

4. SAC!partially overridden by!mus304

5. SAC!partially overridden by!rad50

6. SAC!partially overridden by!grp

7. SAC!partially overridden by!tefu

8. SAC proteins!accumulate at!kinetochores of cav mutant cells

9. BubR1!accumulates at!cav mutant telomeres

10. BubR1!accumulates at!cav mutant telomeres in a mei-41-dependent manner

11. BubR1!accumulates at!cav mutant telomeres in a mus304-dependent manner

12. BubR1!accumulates at!cav mutant telomeres in a rad50-dependent manner

13. BubR1!accumulates at!cav mutant telomeres in a grp-dependent manner

14. BubR1!accumulates at!cav mutant telomeres in a tefu-dependent manner

Statements GPT missed:

1. uncapped telomeres activate the DNA damage response (DDR)

2. Drosophila HOAP protein is required to prevent telomeric fusions

3. HOAP-depleted telomeres activate both the DDR and the spindle assembly checkpoint (SAC).

4. recruitment of BubR1 by dysfunctional telomeres inhibits Cdc20-APC function, preventing the metaphase-to-anaphase transition

Correct, incorrect, missing:

14, 0 , 4

Remarks:

21. Pubmed ID: 1904540

We investigated the recognition of the conserved 5-bp repeated motif NGAAN, which occurs in heat shock gene promoters of Drosophila melanogaster and other eukaryotic organisms, by human heat shock transcription factor (HSF). Extended heat shock element mutants of the human HSP70 gene promoter, containing additional NGAAN blocks flanking the original element, showed significantly higher affinity than the wild-type promoter element for human HSF in vitro. Protein-DNA contact positions were identified by hydroxyl radical protection, diethyl pyrocarbonate interference, and DNase I footprinting. New contacts in the mutant HSE constructs corresponded to the locations of additional NGAAN motifs. The pattern of binding indicated the occurrence of multiple DNA binding modes for HSF with the various constructs and was consistent with an oligomeric, possibly trimeric, structure of the protein. In contrast to the improved binding, the extended heat shock element mutant constructs did not exhibit dramatically increased heat-inducible transcription in transient expression assays with HeLa cells.

Output from GPT:

1. 5-bp repeated motif NGAAN!recognized by!human heat shock transcription factor (HSF)

2. Protein-DNA contact positions!identified by!hydroxyl radical protection, diethyl pyrocarbonate interference, and DNase I footprinting

3. Extended heat shock element mutants of human HSP70 gene promoter!showed higher affinity!for human HSF

Statements GPT missed:

1. extended heat shock element mutant constructs did not exhibit dramatically increased heat-inducible transcription.

Correct, incorrect, missing:

3, 0, 1

Remarks:

22. Pubmed ID: 29449456

Animals in a natural environment confront many sensory cues. Some of these cues bias behavioral decisions independent of experience, and action selection can reveal a stimulus-response (S-R) connection. However, in a changing environment it would be a benefit for an animal to update behavioral action selection based on experience, and learning might modify even strong S-R relationships. How animals use learning to modify S-R relationships is a largely open question. Three sensory stimuli, air, light, and gravity sources were presented to individual Drosophila melanogaster in both naïve and place conditioning situations. Flies were tested for a potential modification of the S-R relationships of anemotaxis, phototaxis, and negative gravitaxis by a contingency that associated place with high temperature. With two stimuli, significant S-R relationships were abandoned when the cue was in conflict with the place learning contingency. The role of the dunce (dnc) cAMP-phosphodiesterase and the rutabaga (rut) adenylyl cyclase were examined in all conditions. Both dnc1 and rut2080 mutant flies failed to display significant S-R relationships with two attractive cues, and have characteristically lower conditioning scores under most conditions. Thus, learning can have profound effects on separate native S-R relationships in multiple contexts, and mutation of the dnc and rut genes reveal complex effects on behavior.

Output from GPT:

1. stimuli!abandoned!S-R relationships

2. cue!in conflict with!place learning contingency

3. learning!has profound effects on!native S-R relationships in multiple contexts

4. dunce (dnc) cAMP-phosphodiesterase!examined in!all conditions

5. rutabaga (rut) adenylyl cyclase!examined in!all conditions

6. dnc1 mutant flies!failed to display!significant S-R relationships with two attractive cues

7. rut2080 mutant flies!failed to display!significant S-R relationships with two attractive cues

8. dnc1 mutant flies!have!lower conditioning scores under most conditions

9. rut2080 mutant flies!have!lower conditioning scores under most conditions

10. mutation of dnc and rut genes!reveal!complex effects on behavior

Statements GPT missed:

10, 0, 0

Correct, incorrect, missing:

Remarks:

Statements 1 and 2 should be together.

23. Pubmed ID: 11014821

Activity of the Darkener of apricot (Doa) locus of Drosophila melanogaster is required for development of the embryonic nervous system, segmentation, photoreceptor maintenance, normal transcription, and sexual differentiation. The gene encodes a protein kinase, with homologues throughout eukaryotes known as the LAMMER kinases. We show here that DOA is expressed as at least two different protein isoforms of 105 and 55 kD throughout development, which are primarily localized to the cytoplasm and nucleus, respectively. Doa transcripts and protein are expressed in all cell types both during embryogenesis and in imaginal discs. Although it was recently shown that DOA kinase is essential for normal sexual differentiation, levels of both kinase isoforms are equal between the sexes during early pupal development. The presence of the kinase on the cell membrane and in the nuclei of polytene salivary gland cells, as well as exclusion from the nuclei of specific cells, may be indicative of regulated kinase localization. Mosaic analysis in both the soma and germline demonstrates that Doa function is essential for cell viability. Finally, in contrast to results reported in other systems and despite some phenotypic similarities, genetic data demonstrate that the LAMMER kinases do not participate in the ras-MAP kinase signal transduction pathway.

Output from GPT:

1. Mosaic analysis!demonstrates!Doa function essential for cell viability

2. genetic data!demonstrate!LAMMER kinases do not participate in ras-MAP kinase signal transduction pathway

Statements GPT missed:

1. Doa transcripts and protein are expressed in all cell types both during embryogenesis and in imaginal discs.

2. DOA kinase is essential for normal sexual differentiation

Correct, incorrect, missing:

2, 0, 2

Remarks:

Statements 1 and 2 are partially correct as they do not specify which neuron/in where.

Missing statements are likely from GPT not having been given that portion of the abstract.

24. Pubmed ID: 1551578

Polytene section 17 of the X chromosome of Drosophila melanogaster, previously known to contain six putative lethal complementation groups important in oogenesis and embryogenesis, has here been further characterized genetically and developmentally. We constructed fcl+Y, a duplication of this region, which allowed us to conduct mutagenesis screens specific for the region and to perform complementation analyses (previously not possible). We recovered 67 new lethal mutations which defined 15 complementation groups within Df(1)N19 which deletes most of polytene section 17. The zygotic lethal phenotypes of these and preexisting mutations within polytene section 17 were examined, and their maternal requirements were analysed in homozygous germline clones using the dominant female sterile technique. We present evidence that an additional gene, which produces two developmentally regulated transcripts, is located in this region and is involved in embryogenesis, although no mutations in this gene were identified. In this interval of 37 to 43 polytene chromosome bands we have defined 17 genes, 12 (71%) of which are of significance to oogenesis or embryogenesis.

Output from GPT:

1. fcl+Y!duplication of!polytene section 17

2. fcl+Y!allows!mutagenesis screens specific for the region

3. fcl+Y!allows!complementation analyses

4. 67 new lethal mutations!recovered from!fcl+Y

5. 67 new lethal mutations!define!15 complementation groups

6. Df(1)N19!deletes most of!polytene section 17

7. Polytene section 17 of X chromosome!contains!six putative lethal complementation groups

8. Polytene section 17 of X chromosome!important in!oogenesis and embryogenesis

Statements GPT missed:

Correct, incorrect, missing:

8, 0, 0

Remarks:

Statement 6 is wrong. It is not Df(1)N19 that deletes sections 17, but rather the 15 groups in Df(1)N19.

Missing statements are likely from GPT not having been given that portion of the abstract.

25. Pubmed ID: 16055700

The ESC-E(Z) complex of Drosophila melanogaster Polycomb group (PcG) repressors is a histone H3 methyltransferase (HMTase). This complex silences fly Hox genes, and related HMTases control germ line development in worms, flowering in plants, and X inactivation in mammals. The fly complex contains a catalytic SET domain subunit, E(Z), plus three noncatalytic subunits, SU(Z)12, ESC, and NURF-55. The four-subunit complex is >1,000-fold more active than E(Z) alone. Here we show that ESC and SU(Z)12 play key roles in potentiating E(Z) HMTase activity. We also show that loss of ESC disrupts global methylation of histone H3-lysine 27 in fly embryos. Subunit mutations identify domains required for catalytic activity and/or binding to specific partners. We describe missense mutations in surface loops of ESC, in the CXC domain of E(Z), and in the conserved VEFS domain of SU(Z)12, which each disrupt HMTase activity but preserve complex assembly. Thus, the E(Z) SET domain requires multiple partner inputs to produce active HMTase. We also find that a recombinant worm complex containing the E(Z) homolog, MES-2, has robust HMTase activity, which depends upon both MES-6, an ESC homolog, and MES-3, a pioneer protein. Thus, although the fly and mammalian PcG complexes absolutely require SU(Z)12, the worm complex generates HMTase activity from a distinct partner set.

Output from GPT:

1. E(Z) SET domain!requires!multiple partner inputs

2. MES-2!depends upon!MES-6

3. MES-2!depends upon!MES-3

4. fly and mammalian PcG complexes!require!SU(Z)12

5. worm complex!generates!HMTase activity

6. E(Z) homolog (MES-2)!has!robust HMTase activity

7. worm complex!generates!from distinct partner set

Statements GPT missed:

1. ESC-E(Z) complex of Drosophila melanogaster Polycomb group (PcG) repressors is a histone H3 methyltransferase

2. ESC and SU(Z)12 play key roles in potentiating E(Z) HMTase activity.

3. loss of ESC disrupts global methylation of histone H3-lysine 27

Correct, incorrect, missing:

7, 0, 3

Remarks:

Statement 1 is partially correct and could have included the fact it is to produce active HMTase.

26. Pubmed ID: 6451215

Mutant strains sensitive and resistant to the drug 5-flourouracil (FU) have been isolated from the wild-type Pac strain of Drosophila melanogaster. The resistant strain, termed flu, is resistant to at least o.00035%FU (2.7x10(-4)M) in the food media and exhibits cross-resistance to 5-fluorodeoxyuridine (FUdR) but not to 5-fluorouridine (FUR). The sensitive strain termed flus, exhibits over 90% mortality on 0.0008%FU (6x10(-15)M). Genetic analysis indicates that the flu gene is located on the third chromosome, which agrees with results of previous workers. An analysis of the enzyme thymidylate synthetase from the selected sensitive and resistant strains indicates that the resistant strain enzyme possesses an elevated specific activity. Levels 4 times that of the sensitive strain were observed when the enzymes were assayed at 20C. This increase is apparently not due to induction by FU in the food media. It is suggested that the enzyme thymidylate synthetase may be involved in the resistance process.

Output from GPT:

1. flu gene!located on!third chromosome

2. enzyme thymidylate synthetase!from!selected sensitive and resistant strains

3. resistant strain enzyme!possesses!elevated specific activity

4. increase!not due to!induction by FU in the food media

5. enzyme thymidylate synthetase!may be involved in!resistance process

6. levels 4 times!observed when!enzymes assayed at 20C

Statements GPT missed:

Correct, incorrect, missing:

6, 0, 0

Remarks:

27. Pubmed ID: 7816022

Drosophila melanogaster F elements are mobile, oligo(A)-terminated DNA sequences that probably propagate by the retrotranscription of RNA intermediates. Polyadenylated transcripts corresponding in size to full-length (4.7 kb) family members were detected in the Drosophila melanogaster Canton-S strain from 2nd larval instar to the adult stage. RNA accumulation reached a maximum in pupae. In the adult, F elements are transcribed in both sexes. F expression is directed in vivo by the intragenic promoter (Fin) located at the 5' end of F. Whole-mount hybridizations were carried out to define the site of synthesis of full-length transcripts found in the ovary. Selective RNA accumulation was not detected in the cytoplasm of any specific cell type. Stained nuclear dots were observed in nurse cells from stage 2-3 to the end of oogenesis. RNase treatment of egg chambers prior to the addition of the probe led to disappearance of the nuclear dots and appearance of a cytoplasmic hybridization signal suggesting leakage of nuclear transcripts. Transgenic lines harbouring the chloramphenicol acetyltransferase (CAT) gene under the control of the Fin promoter were obtained. In independent lines, CAT enzyme levels mirror the ontogenetic profile of F expression drawn from Northern RNA blotting data. An antisense promoter (Fout) that is located downstream from the Fin promoter and transcribe to bords the 5' end of F seems to be constitutively expressed in the fly.

Output from GPT:

1. F elements!are transcribed!in both sexes

2. Whole-mount hybridizations!define!site of synthesis of full-length transcripts

3. Transgenic lines!harbour!chloramphenicol acetyltransferase (CAT) gene

4. CAT enzyme levels!mirror!ontogenetic profile of F expression

5. F expression!is directed!by intragenic promoter (Fin)

6. Fout promoter!transcribes!to bords the 5' end of F

7. Fout promoter!is constitutively expressed!in the fly

Statements GPT missed:

1. Fin is located at the 5' end of F.

2. (Fout) that is located downstream from the Fin promoter

Correct, incorrect, missing:

7, 0, 2

Remarks:

28. Pubmed ID: 27623778

The silent information regulator 1 (Sirt1) has been shown to have negative effects on the Notch pathway in several contexts. We bring evidence that Sirt1 has a positive effect on Notch activation in Drosophila, in the context of sensory organ precursor specification and during wing development. The phenotype of Sirt1 mutant resembles weak Notch loss-of-function phenotypes, and genetic interactions of Sirt1 with the components of the Notch pathway also suggest a positive role for Sirt1 in Notch signalling. Sirt1 is necessary for the efficient activation of enhancer of split [E(spl)] genes by Notch in S2N cells. Additionally, the Notch-dependent response of several E(spl) genes is sensitive to metabolic stress caused by 2-deoxy-d-glucose treatment, in a Sirt1-dependent manner. We found Sirt1 associated with several proteins involved in Notch repression as well as activation, including the cofactor exchange factor Ebi (TBL1), the RLAF/LAF histone chaperone complex and the Tip60 acetylation complex. Moreover, Sirt1 participates in the deacetylation of the CSL transcription factor Suppressor of Hairless. The role of Sirt1 in Notch signalling is, therefore, more complex than previously recognized, and its diverse effects may be explained by a plethora of Sirt1 substrates involved in the regulation of Notch signalling.

Output from GPT:

1. E(spl) genes!are sensitive to!metabolic stress

2. 2-deoxy-d-glucose treatment!causes!metabolic stress

3. Sirt1!is associated with!proteins involved in Notch repression

4. Sirt1!is associated with!Ebi (TBL1)

5. Sirt1!is associated with!RLAF/LAF histone chaperone complex

6. Sirt1!is associated with!Tip60 acetylation complex

7. Notch-dependent response of E(spl) genes!is sensitive to!2-deoxy-d-glucose treatment

8. Sirt1!is associated with!proteins involved in Notch activation

9. Sirt1!participates in!deacetylation of CSL transcription factor Suppressor of Hairless

Statements GPT missed:

1. Sirt1 has a positive effect on Notch activation in Drosophila, in the context of sensory organ precursor specification and during wing development

Correct, incorrect, missing:

8, 1, 1

Remarks:

Statement 1 is wrong as it is the Notch-dependent response that is sensitive, not the gene itself.

29. Pubmed ID: 2353928

cDNA synthesised from liver Poly A+ RNA of estradiol-stimulated male Oreochromis aureus was ligated to PTZ-19R and transformed into E. coli. TG-1 to yield 6 x 10(4) clones/ug DNA, of which 6% was vitellogenin positive. Restriction maps indicate 4 possible subgroups of vitellogenin cDNA, with homology, observed by Southern hybridisation, at a fragment flanked by Pst 1 sites. When the amino acid sequence derived from the DNA sequence of pOA Vg 62 was aligned with that of S. gairdneri, X. laevis, C. elegans and D. melanogaster, homologies of 63%, 22%, 21% and 15%, respectively, were obtained. Estradiol induced a vitellogenin gene transcript of 6500 nucleotides at 6 h and reached a maximum at 72 h after stimulation.

Output from GPT:

1. vitellogenin gene!cloned and characterised!Oreochromis aureus

2. cDNA!synthesised from liver Poly A+ RNA!estradiol-stimulated male Oreochromis aureus

3. cDNA!ligated to PTZ-19R!estradiol-stimulated male Oreochromis aureus

4. cDNA!transformed into E. coli!estradiol-stimulated male Oreochromis aureus

Statements GPT missed:

Correct, incorrect, missing:

2, 2, 0

Remarks:

30. Pubmed ID: 7518199

It is proposed that aging is induced by somatic replication of transposable elements (TEs). Most transposable elements in Drosophila reproduce by reverse transcription. Therefore, inhibitors of reverse transcriptase were tested for their ability to retard aging in Drosophila melanogaster. Two inhibitors, phosphonoformic acid (PFA) and dideoxyinosine (ddI), were capable of prolonging life span when administered for the first half of the adult life. PFA was investigated further. It also produced a reduction in the rate of decline of behavior. PFA appeared not to act on an infectious agent in these experiments, nor did it alter the food intake. Analogues unable to inhibit RT had no life span prolonging effect at similar concentrations to that of PFA.

Output from GPT:

1. aging!induced by!somatic replication of transposable elements (TEs)

2. transposable elements (TEs)!reproduce by!reverse transcription

3. inhibitors of reverse transcriptase!tested for their ability to!retard aging

4. phosphonoformic acid (PFA)!capable of!prolonging life span

5. dideoxyinosine (ddI)!capable of!prolonging life span

6. inhibitors of reverse transcriptase!administered for!first half of the adult life

Statements GPT missed:

1. PFA produced a reduction in the rate of decline of behavior

Correct, incorrect, missing:

6, 0 , 1

Remarks:

Missing statements are likely from GPT not having been given that portion of the abstract.

31. Pubmed ID: 37693001

Biogenic amines are crucial signaling molecules that modulate various physiological life functions both in vertebrates and invertebrates. In humans, these neurotransmitters influence the innate and adaptive immunity systems. In this work, we analyzed whether the aminergic neurotransmission of dopamine, serotonin, and octopamine could have an impact on the humoral innate immune response of Drosophila melanogaster. This is a powerful model system widely used to uncover the insect innate immunity mechanisms which are also conserved in mammals. We found that the neurotransmission of all these amines positively modulates the Toll-responsive antimicrobial peptide (AMP) drosomycin (drs) gene in adult flies infected with the Micrococcus luteus bacterium. Indeed, we showed that either blocking the neurotransmission in their specific aminergic neurons by expressing shibirets (Shits) or silencing the vesicular monoamine transporter gene (dVMAT) by RNAi caused a significantly reduced expression of the Toll-responsive drs gene. However, upon M. luteus infection, the block of aminergic transmission did not alter the expression of AMP attacin genes responding to the immune deficiency (Imd) and Toll pathways. Overall, our results not only reveal a neuroimmune function for biogenic amines in humoral immunity but also further highlight the complexity of the network controlling AMP gene regulation.

Output from GPT:

1. amines!positively modulates!Toll-responsive AMP drosomycin (drs) gene

2. blocking neurotransmission!reduces expression!Toll-responsive drs gene

3. silencing vesicular monoamine transporter gene!reduces expression!Toll-responsive drs gene

4. Micrococcus luteus bacterium!infection!adult flies

Statements GPT missed:

Correct, incorrect, missing:

3, 1, 0

Remarks:

Statement 1 could have specified that the amines involved are dopamine, serotonin, and octopamine.

Statement 2 could have specified that the neurotransmission is blocked by expressing Shits.

Missing statements are likely from GPT not having been given that portion of the abstract.

32. Pubmed ID: 8537310

Three isozymic forms, alpha, beta, and gamma, of Drosophila melanogaster aldolase are produced from a single gene by alternative usage of the triple exons 4 (4 alpha, 4 beta, and 4 gamma) [Shaw-Lee et al. (1992) J. Biol. Chem. 267, 3959-3967; Kim et al. (1992) Mol. Cell. Biol. 12, 773-783; Kai et al. (1992) J. Biochem. 112,677-688]. The expression plasmids for the respective isozymes were transfected into Escherichia coli cells, and the isozymes alpha and beta were purified to homogeneity by a simple procedure, though isozyme gamma was only partially purified. These isozymes are active towards two substrates, fructose-1,6-bisphosphate (Fru-1,6-P2) and fructose-1-phosphate (Fru-1-P), with a preference for Fru-1,6-P2 over Fru-1-P, but they have different kcat/Km values towards these two substrates; isozyme alpha shows the highest value for Fru-1,6-P2. These isozymes show similarity in optimal pHs, thermal stability, and Km values for both Fru-1,6-P2 and Fru-1-P. They are composed of four identical subunits of 40 kDa, forming a tetramer with a molecular weight of approximately 160 kDa. The three isozymes are different in primary structure only at the carboxyl-terminal region encoded by the respective exon 4. Therefore, this region should be primarily responsible for the distinct characteristics of these isozymes.

Output from GPT:

1. Drosophila melanogaster aldolase: !characterization of! isozymes alpha, beta, and gamma

2. Drosophila melanogaster aldolase: !produced from! single gene

3. single gene: !generates! isozymes alpha, beta, and gamma

4. alternative usage: !produces! isozymes alpha, beta, and gamma

5. triple exons 4: !used for! isozymes 4 alpha, 4 beta, and 4 gamma

Statements GPT missed:

Correct, incorrect, missing:

5, 0, 0

Remarks:

Statement 2 is wrong as it is the isozymes that are produced from one gene, not aldolase itself.

Statement 4 should include that it is the alternative use of the triple exons 4.

Missing statements are likely from GPT not having been given that portion of the abstract.

33. Pubmed ID: 17322403

Embryogenesis is controlled by large gene-regulatory networks, which generate spatially and temporally refined patterns of gene expression. Here, we report the characteristics of the regulatory network orchestrating early mesodermal development in the fruitfly Drosophila, where the transcription factor Twist is both necessary and sufficient to drive development. Through the integration of chromatin immunoprecipitation followed by microarray analysis (ChIP-on-chip) experiments during discrete time periods with computational approaches, we identified >2000 Twist-bound cis-regulatory modules (CRMs) and almost 500 direct target genes. Unexpectedly, Twist regulates an almost complete cassette of genes required for cell proliferation in addition to genes essential for morophogenesis and cell migration. Twist targets almost 25% of all annotated Drosophila transcription factors, which may represent the entire set of regulators necessary for the early development of this system. By combining in vivo binding data from Twist, Mef2, Tinman, and Dorsal we have constructed an initial transcriptional network of early mesoderm development. The network topology reveals extensive combinatorial binding, feed-forward regulation, and complex logical outputs as prevalent features. In addition to binary activation and repression, we suggest that Twist binds to almost all mesodermal CRMs to provide the competence to integrate inputs from more specialized transcription factors.

Output from GPT:

1. Twist!targets!annotated Drosophila transcription factors

2. Network topology!reveals!combinatorial binding

3. Network topology!reveals!feed-forward regulation

4. Network topology!reveals!complex logical outputs

5. Twist!represents!entire set of regulators necessary for early development

6. Twist, Mef2, Tinman, Dorsal!combine!in vivo binding data

7. Twist, Mef2, Tinman, Dorsal!construct!initial transcriptional network of early mesoderm development

Statements GPT missed:

1. Twist regulates an almost complete cassette of genes required for cell proliferation in addition to genes essential for morophogenesis and cell migration

Correct, incorrect, missing:

6, 1, 1

Remarks:

Statement 5 is technically wrong as it is the transcription factors being represented, not twist.

Missing statements are likely from GPT not having been given that portion of the abstract.

34. Pubmed ID: 28003491

Stress granules (SGs) are cytosolic ribonucleoprotein aggregates that are induced during cellular stress. Several viruses modulate SG formation, suggesting that SGs have an impact on virus infection. However, the mechanisms and impact of modulating SG assembly in infected cells are not completely understood. In this study, we identify the dicistrovirus cricket paralysis virus 1A (CrPV-1A) protein that functions to inhibit SG assembly during infection. Moreover, besides inhibiting RNA interference, CrPV-1A also inhibits host transcription, which indirectly modulates SG assembly. Thus, CrPV-1A is a multifunctional protein. We identify a key R146A residue that is responsible for these effects, and mutant CrPV(R146A) virus infection is attenuated in Drosophila melanogaster S2 cells and adult fruit flies and results in increased SG formation. Treatment of CrPV(R146A)-infected cells with actinomycin D, which represses transcription, restores SG assembly suppression and viral yield. In summary, CrPV-1A modulates several cellular processes to generate a cellular environment that promotes viral translation and replication.IMPORTANCE RNA viruses encode a limited set of viral proteins to modulate an array of cellular processes in order to facilitate viral replication and inhibit antiviral defenses. In this study, we identified a viral protein, called CrPV-1A, within the dicistrovirus cricket paralysis virus that can inhibit host transcription, modulate viral translation, and block a cellular process called stress granule assembly. We also identified a specific amino acid within CrPV-1A that is important for these cellular processes and that mutant viruses containing mutations of CrPV-1A attenuate virus infection. We also demonstrate that the CrPV-1A protein can also modulate cellular processes in human cells, suggesting that the mode of action of CrPV-1A is conserved. We propose that CrPV-1A is a multifunctional, versatile protein that creates a cellular environment in virus-infected cells that permits productive virus infection.

Output from GPT:

1. Cricket Paralysis Virus 1A protein!disrupts!Stress Granule Formation

2. Stress Granules!are induced by!cellular stress

3. Several viruses!modulate!SG formation

4. SGs!have an impact on!virus infection

Statements GPT missed:

1. CrPV-1A modulates several cellular processes to generate a cellular environment that promotes viral translation and replication.

2. a viral protein, called CrPV-1A, within the dicistrovirus cricket paralysis virus that can inhibit host transcription, modulate viral translation, and block a cellular process called stress granule assembly

Correct, incorrect, missing:

4, 0, 2

Remarks:

Missing statements are likely from GPT not having been given that portion of the abstract.

35. Pubmed ID: 17721789

Transposable elements (TEs) are mobile repetitive DNA sequences that constitute a structurally dynamic component of genomes. In order to understand the dynamics of TEs it is necessary to have information about the control of transposition and its dependence of environmental factors. After a great deal of previous work on transposition conducted on long-term mutation accumulation (MA) lines of Drosophila melanogaster started in 1987, only roo out of 16 families was found active in this genotype. Here we test the effect of the modification of the genetic background by introducing a Cy chromosome, and the effect of extreme temperature (28 degrees C) on the transposition rate of roo. Thermal stress did not affect the transposition rate, whereas the presence of a Cy chromosome in heterozygosis lowered it. There was an excess of insertions in the X chromosome, with respect to autosomes, and in the proximal and distal regions of chromosome arms that can be interpreted as target site preference. One of the control lines became highly unstable with mean insertion and excision rates of 3.0 x 10(-3) and 8.5 x 10(-4), respectively. Instability arose spontaneously during generations of mutation accumulation, and can be attributed to "de novo" mutation. Transposition in the unstable line could be directly studied on the progeny of individual males and females, from where we deduced that transposition occurs mainly, if not exclusively, in males, with a rate of 1.125 insertions per gamete. In situ hybridization with an LTR probe showed that most excisions (12 out of 14) were precise. Our data show the prominent role of genotype in transposition control and can explain rapid turnovers in the genome without increasing the number of copies.

Output from GPT:

1. genotype!affects!transposition of roo

2. extreme temperature!does not affect!transposition rate

3. genetic background!affects!transposition rate of roo

4. presence of Cy chromosome!lowers!transposition rate

Statements GPT missed:

Correct, incorrect, missing:

4, 0, 0

Remarks:

36. Pubmed ID: 26395483

Yolk provides an important source of nutrients during the early development of oviparous organisms. It is composed mainly of vitellogenin proteins packed into membrane-bound compartments called yolk platelets. Catabolism of yolk is initiated by acidification of the yolk platelet, leading to the activation of Cathepsin-like proteinases, but it is unknown how this process is triggered. Yolk catabolism initiates at cellularization in Drosophila melanogaster embryos. Using maternal shRNA technology we found that yolk catabolism depends on the Tor pathway and on the autophagy-initiating kinase Atg1. Whereas Atg1 was required for a burst of spatially regulated autophagy during late cellularization, autophagy was not required for initiating yolk catabolism. We propose that the conserved Tor metabolic sensing pathway regulates yolk catabolism, similar to Tor-dependent metabolic regulation on the lysosome.

Output from GPT:

1. Atg1-Tor pathway!regulates!yolk catabolism

2. Yolk!provides!nutrients

3. Yolk catabolism!initiates at!cellularization

4. Atg1!required for!burst of spatially regulated autophagy

5. Autophagy!not required for!initiating yolk catabolism

6. Tor metabolic sensing pathway!regulates!yolk catabolism

7. Tor-dependent metabolic regulation!on!lysosome

8. Maternal shRNA technology!found that!yolk catabolism depends on Tor pathway

9. Maternal shRNA technology!found that!yolk catabolism depends on Atg1

Statements GPT missed:

Correct, incorrect, missing:

9, 0, 0

Remarks:

37. Pubmed ID: 14960014

In Drosophila melanogaster food search behaviour, groups of flies swarm around and aggregate on patches of food. We wondered whether flies explore their environment in a cooperative way as interactions between individual flies within a population might influence the flies' ability to locate food sources. We have shown that the food search behavior in the fruit fly Drosophila is a two-step process. Firstly, 'primer' flies search the environment and randomly land on different food patches. Secondly, the remaining group of flies move to the most favorable food source and aggregate there. We call this a 'search-aggregation' cycle. Our data demonstrate that flies do not individually assess all available food resources. Rather, social interactions between flies appear to affect their choice of a specific food patch. A genetic analysis of this 'search-aggregation' behavior shows that flies carrying mutations in specific genes (for example, the dunce (dnc) gene which codes for a phosphodiesterase) were defective in this search-aggregation behavior when compared to normal flies. Future investigations of the neuronal signaling involved in this behavior will help us to understand the complexities of this aspect of Drosophila social behaviour.

Output from GPT:

1. genetic analysis!shows!flies carrying mutations in specific genes

2. dunce (dnc) gene!codes for!phosphodiesterase

3. flies carrying mutations in specific genes!defective in!search-aggregation behavior

4. future investigations!will help us understand!neuronal signaling involved in this behavior

5. neuronal signaling!involved in!search-aggregation behavior

6. social interactions between flies!affect!choice of specific food patch

7. Drosophila social behavior!has complexities!in this aspect

Statements GPT missed:

Correct, incorrect, missing:

7, 0, 0

Remarks:

38. Pubmed ID: 27183194

Proper gene expression requires coordinated interplay among transcriptional coactivators, transcription factors and the general transcription machinery. We report here that MSL1, a central component of the dosage compensation complex in Drosophila melanogaster and Drosophila virilis, displays evolutionarily conserved sex-independent binding to promoters. Genetic and biochemical analyses reveal a functional interaction of MSL1 with CDK7, a subunit of the Cdk-activating kinase (CAK) complex of the general transcription factor TFIIH. Importantly, MSL1 depletion leads to decreased phosphorylation of Ser5 of RNA polymerase II. In addition, we demonstrate that MSL1 is a phosphoprotein, and transgenic flies expressing MSL1 phosphomutants show mislocalization of the histone acetyltransferase MOF and histone H4 K16 acetylation, thus ultimately causing male lethality due to a failure of dosage compensation. We propose that, by virtue of its interaction with components of the general transcription machinery, MSL1 exists in different phosphorylation states, thereby modulating transcription in flies.

Output from GPT:

1. MSL1 depletion!leads to!decreased phosphorylation of Ser5 of RNA polymerase II

2. MSL1!is a!phosphoprotein

3. transgenic flies expressing MSL1 phosphomutants!show mislocalization of!histone acetyltransferase MOF

4. transgenic flies expressing MSL1 phosphomutants!show mislocalization of!histone H4 K16 acetylation

5. male lethality!is due to a failure of!dosage compensation

6. MSL1!interacts with!components of the general transcription machinery

7. MSL1!modulates transcription in!flies

8. mislocalization of histone acetyltransferase MOF!causes!male lethality

9. mislocalization of histone H4 K16 acetylation!causes!male lethality

Statements GPT missed:

1. MSL1 displays evolutionarily conserved sex-independent binding to promoters

Correct, incorrect, missing:

9, 0, 1

Remarks:

Missing statements are likely from GPT not having been given that portion of the abstract.

39. Pubmed ID: 27986833

The occurrence and biological importance of sialic acid (Sia) and its metabolic enzymes in insects have been studied using Drosophila melanogaster. The most prominent feature of D. melanogaster CMP-Sia synthetase (DmCSS) is its Golgi-localization, contrasted with nuclear localization of vertebrate CSSs. However, it remains unclear if the Golgi-localization is common to other insect CSSs and why it happens. To answer these questions, Aedes aegypti (mosquito) CSS (AaCSS) and Tribolium castaneum (beetle) CSS (TcCSS) were cloned and characterized for their activity and subcellular localization. Our new findings show: (1) AaCSS and TcCSS share a common overall structure with DmCSS in terms of evolutionarily conserved motifs and the absence of the C-terminal domain typical to vertebrate CSSs; (2) when expressed in mammalian and insect cells, AaCSS and TcCSS showed in vivo and in vitro CSS activities, similar to DmCSS. In contrast, when expressed in bacteria, they lacked CSS activity because the N-terminal hydrophobic region appeared to induce protein aggregation; (3) when expressed in Drosophila S2 cells, AaCSS and TcCSS were predominantly localized in the ER, but not in the Golgi. Surprisingly, DmCSS was mainly secreted into the culture medium, although partially detected in Golgi. Consistent with these results, the N-terminal hydrophobic regions of AaCSS and TcCSS functioned as a signal peptide to render them soluble in the ER, while the N-terminus of DmCSS functioned as a membrane-spanning region of type II transmembrane proteins whose cytosolic KLK sequence functioned as an ER export signal. Accordingly, the differential subcellular localization of insect CSSs are distinctively more diverse than previously recognized.

Output from GPT:

1. DmCSS!has!Golgi-localization

2. vertebrate CSSs!have!nuclear localization

3. insect CSSs!have!diverse subcellular localizations

4. Golgi-localization!is common!to other insect CSSs

5. occurrence and biological importance of Sia and its metabolic enzymes!have been studied!using Drosophila melanogaster

Statements GPT missed:

Correct, incorrect, missing:

4, 1, 5

Remarks:

Statement 4 is wrong. Abstract stated that it remains unclear.

Missing statements are likely from GPT not having been given that portion of the abstract.

40. Pubmed ID: 28439025

Ecdysis-triggering hormone (ETH) was originally discovered and characterized as a molt termination signal in insects through its regulation of the ecdysis sequence. Here we report that ETH persists in adult Drosophila melanogaster, where it functions as an obligatory allatotropin to promote juvenile hormone (JH) production and reproduction. ETH signaling deficits lead to sharply reduced JH levels and consequent reductions of ovary size, egg production, and yolk deposition in mature oocytes. Expression of ETH and ETH receptor genes is in turn dependent on ecdysone (20E). Furthermore, 20E receptor knockdown specifically in Inka cells reduces fecundity. Our findings indicate that the canonical developmental roles of 20E, ETH, and JH during juvenile stages are repurposed to function as an endocrine network essential for reproductive success.

Output from GPT:

1. Endocrine network!essential for!reproductive success

2. Ecdysis-triggering hormone (ETH)!regulates!ecdysis sequence

3. ETH!functions as!allatotropin

4. ETH!promotes!juvenile hormone (JH) production

5. ETH signaling deficits!lead to!reduced JH levels

6. Expression of ETH and ETH receptor genes!depends on!ecdysone (20E)

7. 20E receptor knockdown!reduces!fecundity

8. ETH signaling deficits!lead to!reduced ovary size

9. ETH signaling deficits!lead to!reduced egg production

10. ETH signaling deficits!lead to!reduced yolk deposition

11. Canonical roles of 20E, ETH, and JH!repurposed as!endocrine network for reproductive success

Statements GPT missed:

1. ETH promotes reproduction

Correct, incorrect, missing:

11, 0, 1

Remarks:

41. Pubmed ID: 8307322

Variation of eight allozyme loci, Acph, Adh, Est-6, Est-C, alpha-Gpdh, Idh (NADP-dependent), 6-Pgd and Pgm, was studied in 20 populations of Drosophila melanogaster from the territory of the former USSR, including Eastern Europe, the Caucasus and Central Asia. Adh exhibited weak regression on latitude, longitude, and also on the temperature of the hottest calendar month in these localities. For seven other loci, no regular geographical pattern was found. Nei's coefficient of gene differentiation (GST) ranged from 0.010 (6-Pgd, statistically nonsignificant) to 0.178 (Idh) and was on average 0.058. The average Nei's genetic distance (D) between the populations was 0.011. It is suggested that this fairly low level of interpopulation differentiation for the examined loci except Idh may be explained by extensive gene flow in combination with natural selection.

Output from GPT:

1. Adh!exhibits weak regression on!latitude

2. Adh!exhibits weak regression on!longitude

3. Adh!exhibits weak regression on!temperature of the hottest calendar month

4. Extensive gene flow!explains!low level of interpopulation differentiation

5. Natural selection!explains!low level of interpopulation differentiation

6. - Acph!varies in!20 populations

7. - Adh!varies in!20 populations

8. - Est-6!varies in!20 populations

9. - Est-C!varies in!20 populations

10. - alpha-Gpdh!varies in!20 populations

11. - Idh (NADP-dependent)!varies in!20 populations

12. - 6-Pgd!varies in!20 populations

13. - Pgm!varies in!20 populations

Statements GPT missed:

Correct, incorrect, missing:

13, 0, 0

Remarks:

52. Pubmed ID: 16260610

Menin, the product of the multiple endocrine neoplasia type I gene, has been implicated in several biological processes, including the control of gene expression and apoptosis, the modulation of mitogen-activated protein kinase pathways, and DNA damage sensing or repair. In this study, we have investigated the function of menin in the model organism Drosophila melanogaster. We show that Drosophila lines overexpressing menin or an RNA interference for this gene develop normally but are impaired in their response to several stresses, including heat shock, hypoxia, hyperosmolarity and oxidative stress. In the embryo subjected to heat shock, this impairment was characterized by a high degree of developmental arrest and lethality. The overexpression of menin enhanced the expression of HSP70 in embryos and interfered with its down-regulation during recovery at the normal temperature. In contrast, the inhibition of menin with RNA interference reduced the induction of HSP70 and blocked the activation of HSP23 upon heat shock, Menin was recruited to the Hsp70 promoter upon heat shock and menin overexpression stimulated the activity of this promoter in embryos. A 70-kDa inducible form of menin was expressed in response to heat shock, indicating that menin is also regulated in conditions of stress. The induction of HSP70 and HSP23 was markedly reduced or absent in mutant embryos harboring a deletion of the menin gene. These embryos, which did not express the heat shock-inducible form of menin, were also hypersensitive to various conditions of stress. These results suggest a novel role for menin in the control of the stress response and in processes associated with the maintenance of protein integrity.

Output from GPT:

1. overexpression of menin: !enhanced expression of! HSP70 in embryos

2. overexpression of menin: !interfered with! down-regulation of HSP70 during recovery at normal temperature

3. inhibition of menin with RNA interference: !reduced induction of! HSP70

4. inhibition of menin with RNA interference: !blocked activation of! HSP23 upon heat shock

5. menin: !recruited to! Hsp70 promoter upon heat shock

6. menin overexpression: !stimulated activity of! Hsp70 promoter in embryos

7. 70-kDa inducible form of menin: !expressed in response to! heat shock

8. induction of HSP70 and HSP23: !markedly reduced or absent in! mutant embryos with deletion of menin gene

9. mutant embryos with deletion of menin gene: !hypersensitive to! various conditions of stress

Statements GPT missed:

1. Menin, the product of the multiple endocrine neoplasia type I gene

Correct, incorrect, missing:

9, 0, 3

Remarks:

Missing statements are likely from GPT not having been given that portion of the abstract.

43. Pubmed ID: 17725801

The midgut proteome of Drosophila melanogaster was compared in larvae fed dietary Bowman-Birk inhibitor (BBI) vs. larvae fed a control diet. By using two-dimensional gel electrophoresis, nine differentially expressed proteins were observed, which were associated with enzymes or transport functions such as sterol carrier protein X (SCPX), ubiquitin-conjugating enzyme, endopeptidase, receptor signalling protein kinase, ATP-dependent RNA helicase and alpha-tocopherol transport. Quantitative real-time PCR verified differential expression of transcripts coding for six of the proteins observed from the proteomic analysis. BBI evidently affects expression of proteins associated with protein degradation, transport and fatty acid catabolism. We then tested the hypothesis that SCPX was critical for the Drosophila third instars' response to BBI treatment. Inhibition of SCPX caused the third instars to become more susceptible to dietary BBI.

Output from GPT:

1. larvae fed a control diet!observed!nine differentially expressed proteins

2. differentially expressed proteins!associated with!enzymes or transport functions

3. enzymes or transport functions!include!sterol carrier protein X (SCPX)

4. enzymes or transport functions!include!ubiquitin-conjugating enzyme

5. enzymes or transport functions!include!endopeptidase

6. enzymes or transport functions!include!receptor signalling protein kinase

7. enzymes or transport functions!include!ATP-dependent RNA helicase

8. enzymes or transport functions!include!alpha-tocopherol transport

9. quantitative real-time PCR!verified!differential expression of transcripts

10. proteomic analysis!observed!differential expression of transcripts coding for six proteins

Statements GPT missed:

1. Inhibition of SCPX caused the third instars to become more susceptible to dietary BBI

Correct, incorrect, missing:

10, 0, 1

Remarks:

44. Pubmed ID: 16328894

This study investigated the working hypothesis that two widely used organophosphate pesticides; Nuvan and Dimecron, exert toxic effects in Drosophila. Transgenic D. melanogaster (hsp70-lacZ) was used as a model for assaying stress gene expression and AchE activity as an endpoint for toxicity and also to evaluate whether stress gene expression is sufficient to protect against toxic insult of the chemicals and to prevent tissue damage. The study was extended to investigate the effect of the pesticides on the life cycle and reproduction of the organism. The study showed that Nuvan affected emergence of the exposed flies more drastically than Dimecron and the effect was lethal at the highest tested concentration (0.075 ppm). While Nuvan at 0.0075 and 0.015 ppm concentrations affected reproduction of the flies significantly, the effect of Dimecron was significant only at 0.015 and 0.075 ppm. Nuvan-exposed third-instar larvae exhibited a 1.2-fold to 1.5-fold greater hsp70 expression compared to Dimecron at concentrations ranging from 0.0075 to 0.075 ppm following 12 and 18 h exposure. While maximum expression of hsp70 was observed in Nuvan-exposed third-instar larval tissues following 18 h exposure at 0.075 ppm, Dimecron at the same dietary concentration induced a maximum expression of hsp70 following 24 h exposure. Further, concomitant with a significant induction of hsp70, significant inhibition of AchE was observed following chemical exposure and temperature shock. Concurrent with a significant decline in hsp70 expression in Nuvan-exposed larvae after 48 h at 0.075 ppm, tissue damage was evident. Dimecron-exposed larvae exhibited a plateau in hsp70 induction even after 48 h exposure and moderate tissue damage was observed in these larvae. The present study suggests that Nuvan is more cytotoxic than Dimecron in transgenic Drosophila melanogaster.

Output from GPT:

1. Nuvan!exhibits greater hsp70 expression!compared to Dimecron

2. Chemical exposure!inhibits!AchE activity

3. Nuvan!affects!reproduction of flies

4. Dimecron!affects!reproduction of flies

5. Nuvan!exhibits maximum hsp70 expression!at 0.075 ppm and 18 h exposure

6. Dimecron!exhibits maximum hsp70 expression!at 0.075 ppm and 24 h exposure

7. Nuvan!causes tissue damage!at 0.075 ppm after 48 h exposure

8. Nuvan!is more cytotoxic!than Dimecron in Drosophila melanogaster

Statements GPT missed:

1. pesticides; Nuvan and Dimecron, exert toxic effects in Drosophila

Correct, incorrect, missing:

8, 0, 1

Remarks:

45. Pubmed ID: 1981765

The metallothionein genes of Drosophila melanogaster, Mtn and Mto, may play an important role in heavy metal detoxification. Several different tandem duplications of Mtn have been shown to increase cadmium and copper tolerance, as well as Mtn expression. In order to investigate the possibility of increased selection for duplications of these genes in natural populations exposed to high levels of heavy metals, we compared the frequencies of such duplications among flies collected from metal-contaminated and non-contaminated orchards in Pennsylvania, Tennessee and Georgia. Restriction enzyme analysis was used to screen 1666 wild third chromosomes for Mtn duplications and a subset (327) of these lines for Mto duplications. The frequency of pooled Mtn duplications found ranged from 0% to 20%, and was not significantly higher at the contaminated sites. No Mto duplications were identified. Estimates of sequence diversity at the Mtn locus among a subsample (92) of the duplication survey were obtained using four-cutter analysis. This analysis revealed a low level of polymorphism, consistent with both selection at the Mtn locus, and a fairly recent origin for the duplications. To further examine this hypothesis, we sequenced an Mtn allele of Drosophila simulans and measured the amount of nucleotide sequence divergence between D. simulans and its sibling species D. melanogaster. The levels of silent nucleotide polymorphism and divergence in the Mtn region were compared with those in the Adh region, using the neutrality test of R.R. Hudson, M. Kreitman and M. Aguadé.

Output from GPT:

1. Drosophila melanogaster!has!metallothionein genes

2. Mtn!may play a role in!heavy metal detoxification

3. Mto!may play a role in!heavy metal detoxification

4. Mtn duplications!increase!cadmium and copper tolerance

5. Mtn duplications!increase!Mtn expression

6. Restriction enzyme analysis!was used to screen!wild third chromosomes for Mtn duplications

7. Frequency of pooled Mtn duplications!ranged from!0% to 20%

8. Mtn!has!several tandem duplications

9. Natural populations!may have increased selection for!duplications of Mtn genes

10. Flies from metal-contaminated and non-contaminated orchards!were compared for!Mtn duplications

11. Subset of lines!were screened for!Mto duplications

12. Frequency of pooled Mtn duplications!was not significantly higher!at contaminated sites

13. Sequence diversity at Mtn locus!was estimated using!four-cutter analysis

Statements GPT missed:

Correct, incorrect, missing:

13, 0, 0

Remarks:

46. Pubmed ID: 18694958

Stress-activated protein kinases such as p38 regulate the activity of transcription factor ATF-2. However, the physiological role of ATF-2, especially in the brain, is unknown. Here, we found that Drosophila melanogaster ATF-2 (dATF-2) is expressed in large ventral lateral neurons (l-LN(v)s) and also, to a much lesser extent, in small ventral lateral neurons, the pacemaker neurons. Only l-LN(v)s were stained with the antibody that specifically recognizes phosphorylated dATF-2, suggesting that dATF-2 is activated specifically in l-LN(v)s. The knockdown of dATF-2 in pacemaker neurons using RNA interference decreased sleep time, whereas the ectopic expression of dATF-2 increased sleep time. dATF-2 knockdown decreased the length of sleep bouts but not the number of bouts. The ATF-2 level also affected the sleep rebound after sleep deprivation and the arousal threshold. dATF-2 negatively regulated locomotor activity, although it did not affect the circadian locomotor rhythm. The degree of dATF-2 phosphorylation was greater in the morning than at night and was enhanced by forced locomotion via the dp38 pathway. Thus, dATF-2 is activated by the locomotor while it increases sleep, suggesting a role for dATF-2 as a regulator to connect sleep with locomotion.

Output from GPT:

1. dATF-2!negatively regulates!locomotor activity

2. dATF-2!does not affect!circadian locomotor rhythm

3. dATF-2 phosphorylation!is greater in the morning!than at night

4. dATF-2 phosphorylation!is enhanced by forced locomotion!via the dp38 pathway

5. dATF-2!is activated by locomotion!while it increases sleep

Statements GPT missed:

1. p38 regulate the activity of transcription factor ATF-2

2. ATF-2 (dATF-2) is expressed in large ventral lateral neurons (l-LN(v)s)

3. dATF-2 is activated specifically in l-LN(v)s

4. dATF-2 knockdown decreased the length of sleep bouts but not the number of bouts

Correct, incorrect, missing:

5, 0 ,4

Remarks:

Missing statements are likely from GPT not having been given that portion of the abstract.

47. Pubmed ID: 35029786

3-Hydroxykynurenine as a Potential Ligand for Hsp70 Proteins and Its Effects on Drosophila Memory After Heat Shock Kynurenine products of tryptophan metabolism are modifiers of the nervous activity and oxidative processes in mammals and invertebrates. 3-Hydroxykynurenine (3HOK) in moderate concentrations is a lipid peroxidation inhibitor. However, its accumulation and oxidative auto-dimerization lead to oxidative stress development manifested in age-related neurodegenerative diseases (NDD) and neurological disorders provoked by acute stress. Different forms of stress, the mostly studied being heat shock response, rely on functioning of heat shock proteins of the Hsp70 superfamily. Since kynurenines are called "kids of stress," we performed computational estimation of affinity of 3HOK and other kynurenines binding to predicted ATP site of Drosophila melanogaster Hsp cognate 71 protein (Dhsp71) using AutoDock Vina. The binding energy of 3HOK dimer is - 9.4 kcal/mol; its orientation within the active site is close to that of ATP. This might be a new mechanism of producing a competitive inhibitor of Hsp70 chaperones that decreases organism ability to adapt to heat shock. We also showed that the Drosophila cardinal (cd1) mutant with 3HOK excess, serving as a model for Huntington's disease (HD), manifests severe defects of short-term memory after heat shock applied either in adults or at the prepupal stage.

Output from GPT:

1. 3-Hydroxykynurenine!is a potential ligand for!Hsp70 proteins

2. 3-Hydroxykynurenine!affects!Drosophila memory after heat shock

3. Kynurenine products of tryptophan metabolism!modify!nervous activity and oxidative processes

4. Different forms of stress!rely on!functioning of heat shock proteins

5. Heat shock response!relies on!functioning of heat shock proteins

6. 3HOK dimer!produces!a competitive inhibitor of Hsp70 chaperones

7. Competitive inhibitor of Hsp70 chaperones!decreases!organism ability to adapt to heat shock

8. Heat shock!can be applied!in adults or at the prepupal stage

9. Kynurenine accumulation!leads to!oxidative stress development

10. Kynurenine accumulation!leads to!age-related neurodegenerative diseases

11. Kynurenine accumulation!leads to!neurological disorders provoked by acute stress

12. Kynurenines!are called! "kids of stress"

13. 3HOK!binds to!predicted ATP site of Dhsp71

14. 3HOK dimer!has a binding energy of!-9.4 kcal/mol

15. 3HOK dimer!has an orientation close to!ATP within the active site

16. Drosophila cardinal (cd1) mutant!manifests!severe defects of short-term memory after heat shock

17. Drosophila cardinal (cd1) mutant!has 3HOK excess and!serves as a model for Huntington's disease (HD)

Statements GPT missed:

1. 3-Hydroxykynurenine (3HOK) in moderate concentrations is a lipid peroxidation inhibitor.

Correct, incorrect, missing:

17, 0, 1

Remarks:

48. Pubmed ID: 12223399

COP9 signalosome subunits 4 and 5 regulate multiple pleiotropic pathways in Drosophila melanogaster. The COP9 signalosome (CSN) is an essential eight-subunit repressor of light-regulated development in Arabidopsis. This complex has also been identified in animals, though its developmental role remains obscure. CSN subunits have been implicated in various cellular processes, suggesting a possible role for the CSN as an integrator of multiple signaling pathways. In order to elucidate the function of the CSN in animals, a Drosophila model system has previously been established. Gel-filtration analysis with antibodies against CSN subunits 4, 5 and 7 revealed that these proteins act as a complex in Drosophila that is similar in size to the plant and mammalian complexes. Null mutations in either one of two subunits, CSN4 or CSN5, are larval lethal. Successful embryogenesis appears to be a consequence of maternal contribution of the complex. Biochemical analysis indicates that the different subunits are found in both CSN-dependent and CSN-independent forms, and that these forms are differentially affected by the mutations. Phenotypic characterization of these two mutants indicates that they show both shared and unique phenotypes, which suggest specific roles for each subunit. Both mutants have defective oocyte and embryo patterning, and defects in response to DNA damage, while csn5 mutants develop melanotic tumors and csn4 mutants have phenotypes reminiscent of defects in ecdysone signaling.

Output from GPT:

1. COP9 signalosome subunits 4!regulate!multiple pleiotropic pathways

2. COP9 signalosome!has been identified!in animals

3. CSN subunits!have been implicated!in various cellular processes

4. CSN!acts as!a complex in Drosophila

5. Successful embryogenesis!is a consequence of!maternal contribution of the complex

6. CSN subunits!are found!in both CSN-dependent and CSN-independent forms

7. csn5 mutants!develop!melanotic tumors

8. csn4 mutants!have phenotypes reminiscent of!defects in ecdysone signaling

9. COP9 signalosome subunits 5!regulate!multiple pleiotropic pathways

Statements GPT missed:

1. The COP9 signalosome (CSN) is an essential eight-subunit repressor of light-regulated development

2. Null mutations in either one of two subunits, CSN4 or CSN5, are larval lethal

Correct, incorrect, missing:

8, 1, 2

Remarks:

Statement 4 should specify it is subunits 4 5 and 7.

49. Pubmed ID: 9829824

The Cbf5 protein of Saccharomyces cerevisiae was originally identified as a low-affinity centromeric DNA-binding protein, and chf5 mutants have a defect in rRNA synthesis. A closely related protein from mammals, NAP57, is a nucleolar protein that coimmunoprecipitates with the nucleolar phosphoprotein Nopp140. To study the function of this protein family in a higher eukaryote that is amenable to genetic approaches, the gene encoding a Drosophila melanogaster homolog, Nop60B, was identified. The predicted Drosophila protein shares a high degree of sequence identity over a 380-residue region with both the mammalian and yeast proteins, and shares several conserved motifs with the prokaryotic tRNA pseudouridine 55 synthases. Nop60B RNA is found at high levels in nurse cells and in the oocyte, and is present throughout development. Nop60B protein is localized primarily to the nucleolus of interphase cells, and is absent from the chromosomes during mitosis. Nop60B mutants were generated and shown to be homozygous lethal. The Drosophila gene can rescue the lethal phenotype of yeast chf5 mutations, showing that the function of this protein has been conserved from yeast to Drosophila.

Output from GPT:

1. Cbf5 protein!originally identified as!low-affinity centromeric DNA-binding protein

2. chf5 mutants!have a defect in!rRNA synthesis

3. gene encoding!Drosophila melanogaster homolog!identified

4. NAP57 protein!closely related to!mammals

5. NAP57 protein!coimmunoprecipitates with!Nopp140 protein

Statements GPT missed:

1. Nop60B RNA is found at high levels in nurse cells and in the oocyte, and is present throughout development

2. Nop60B protein is localized primarily to the nucleolus of interphase cells, and is absent from the chromosomes during mitosis.

Correct, incorrect, missing:

5, 0, 2

Remarks:

Statement 3 should specify Nop60B.

Missing statements are likely from GPT not having been given that portion of the abstract.

50. Pubmed ID: 24526728

Drosophila melanogaster is a model system for examining the mechanisms of action of neuropeptides. DPKQDFMRFamide was previously shown to induce contractions in Drosophila body wall muscle fibres in a Ca(2+)-dependent manner. The present study examined the possible involvement of a G-protein-coupled receptor and second messengers in mediating this myotropic effect after removal of the central nervous system. DPKQDFMRFamide-induced contractions were reduced by 70% and 90%, respectively, in larvae with reduced expression of the Drosophila Fmrf receptor (FR) either ubiquitously or specifically in muscle tissue, compared with the response in control larvae in which expression was not manipulated. No such effect occurred in larvae with reduced expression of this gene only in neurons. The myogenic effects of DPKQDFMRFamide do not appear to be mediated through either of the two Drosophila myosuppressin receptors (DmsR-1 and DmsR-2). DPKQDFMRFamide-induced contractions were not reduced in Ala1 transgenic flies lacking activity of calcium/calmodulin-dependent protein kinase (CamKII), and were not affected by the CaMKII inhibitor KN-93. Peptide-induced contractions in the mutants of the phospholipase C-β (PLCβ) gene (norpA larvae) and in IP3 receptor mutants were similar to contractions elicited in control larvae. The peptide failed to increase cAMP and cGMP levels in Drosophila body wall muscles. Peptide-induced contractions were not potentiated by 3-isobutyl-1-methylxanthine, a phosphodiesterase inhibitor, and were not antagonized by inhibitors of cAMP-dependent or cGMP-dependent protein kinases. Additionally, exogenous application of arachidonic acid failed to induce myogenic contractions. Thus, DPKQDFMRFamide induces contractions via a G-protein coupled FMRFamide receptor in muscle cells but does not appear to act via cAMP, cGMP, IP3, PLC, CaMKII or arachidonic acid.

Output from GPT:

1. DPKQDFMRFamide-induced contractions: !not reduced in! Ala1 transgenic flies lacking activity of calcium/calmodulin-dependent protein kinase (CamKII)

2. DPKQDFMRFamide-induced contractions: !not affected by! CaMKII inhibitor KN-93

3. Peptide-induced contractions: !similar to! contractions elicited in mutants of the phospholipase C-β (PLCβ) gene (norpA larvae)

4. Peptide-induced contractions: !similar to! contractions elicited in IP3 receptor mutants

5. Peptide: !failed to increase! cAMP levels in Drosophila body wall muscles

6. Peptide: !failed to increase! cGMP levels in Drosophila body wall muscles

Statements GPT missed:

1. DPKQDFMRFamide was previously shown to induce contractions in Drosophila body wall muscle fibres in a Ca(2+)-dependent manner

2. exogenous application of arachidonic acid failed to induce myogenic contractions

3. DPKQDFMRFamide induces contractions via a G-protein coupled FMRFamide receptor in muscle cells but does not appear to act via cAMP, cGMP, IP3, PLC, CaMKII or arachidonic acid.

Correct, incorrect, missing:

4, 2, 4

Remarks:

Missing statements are likely from GPT not having been given that portion of the abstract.
